# Leveraging Base Pair Mammalian Constraint to Understand Genetic Variation and Human Disease

**DOI:** 10.1101/2023.03.10.531987

**Authors:** Patrick F. Sullivan, Jennifer R. S. Meadows, Steven Gazal, BaDoi N. Phan, Xue Li, Diane P. Genereux, Michael X. Dong, Matteo Bianchi, Gregory Andrews, Sharadha Sakthikumar, Jessika Nordin, Ananya Roy, Matthew J. Christmas, Voichita D. Marinescu, Ola Wallerman, James R. Xue, Yun Li, Shuyang Yao, Quan Sun, Jin Szatkiewicz, Jia Wen, Laura M. Huckins, Alyssa J. Lawler, Kathleen C. Keough, Zhili Zheng, Jian Zeng, Naomi R. Wray, Jessica Johnson, Jiawen Chen, Zoonomia Consortium, Benedict Paten, Steven K. Reilly, Graham M. Hughes, Zhiping Weng, Katherine S. Pollard, Andreas R. Pfenning, Karin Forsberg-Nilsson, Elinor K. Karlsson, Kerstin Lindblad-Toh

**Affiliations:** Department of Genetics, University of North Carolina Medical School; Chapel Hill, NC 27599, USA; Department of Medical Epidemiology and Biostatistics, Karolinska Institutet; Stockholm, Sweden; Department of Medical Biochemistry and Microbiology, Science for Life Laboratory, Uppsala University; Uppsala, 751 32, Sweden; Keck School of Medicine, University of Southern California; Los Angeles, CA 90033, USA; Department of Computational Biology, School of Computer Science, Carnegie Mellon University; Pittsburgh, PA 15213, USA; Medical Scientist Training Program, University of Pittsburgh School of Medicine; Pittsburgh, PA 15261, USA; Neuroscience Institute, Carnegie Mellon University; Pittsburgh, PA 15213, USA; Broad Institute of MIT and Harvard; Cambridge, MA 02139, USA; Morningside Graduate School of Biomedical Sciences, UMass Chan Medical School; Worcester, MA 01605, USA; Program in Bioinformatics and Integrative Biology, UMass Chan Medical School; Worcester, MA 01605, USA; Department of Immunology, Genetics and Pathology, Science for Life Laboratory, Uppsala University; Uppsala, 751 85, Sweden; Department of Organismic and Evolutionary Biology, Harvard University; Cambridge, MA 02138, USA; Department of Biostatistics, University of North Carolina at Chapel Hill; Chapel Hill, NC, USA; Department of Psychiatry, Icahn School of Medicine at Mount Sinai; New York, NY 10029, USA; Department of Biological Sciences, Mellon College of Science, Carnegie Mellon University; Pittsburgh, PA 15213, USA; Department of Epidemiology & Biostatistics, University of California San Francisco; San Francisco, CA 94158, USA; Fauna Bio Incorporated; Emeryville, CA 94608, USA; Gladstone Institutes; San Francisco, CA 94158, USA; Institute for Molecular Bioscience, University of Queensland; Brisbane, Queensland, Australia; Queensland Brain Institute, University of Queensland; Brisbane, Queensland, Australia; Department of Genetics and Genomic Sciences, Icahn School of Medicine at Mount Sinai; New York, NY 10029, USA; Genomics Institute, University of California Santa Cruz; Santa Cruz, CA 95064, USA; Department of Genetics, Yale School of Medicine; New Haven, CT 06510, USA; School of Biology and Environmental Science, University College Dublin; Belfield, Dublin 4, Ireland; Chan Zuckerberg Biohub; San Francisco, CA 94158, USA; Biodiscovery Institute, University of Nottingham; Nottingham, UK; Program in Molecular Medicine, UMass Chan Medical School; Worcester, MA 01605, USA

## Abstract

Although thousands of genomic regions have been associated with heritable human diseases, attempts to elucidate biological mechanisms are impeded by a general inability to discern which genomic positions are functionally important. Evolutionary constraint is a powerful predictor of function that is agnostic to cell type or disease mechanism. Here, single base phyloP scores from the whole genome alignment of 240 placental mammals identified 3.5% of the human genome as significantly constrained, and likely functional. We compared these scores to large-scale genome annotation, genome-wide association studies (GWAS), copy number variation, clinical genetics findings, and cancer data sets. Evolutionarily constrained positions are enriched for variants explaining common disease heritability (more than any other functional annotation). Our results improve variant annotation but also highlight that the regulatory landscape of the human genome still needs to be further explored and linked to disease.

## Introduction

In the past 15 years, increasingly larger genomic studies have delivered many novel associations for a wide array of human diseases, disorders, biomarkers, and other traits. Approximately 200K genetic associations have been identified that span the allelic spectrum, from ultra-rare variants in large sequencing datasets to variants common in all humans, in both coding and regulatory regions (see *Supplementary Methods, Section 1*). Although these associations meet rigorous standards for statistical significance and replicability, their functional importance is generally unknown. Inferring functional importance is crucial to translating the results of rare and common variant association studies into the biological, clinical, and therapeutic knowledge required to understand and treat human disease. Exceptional efforts have been made to annotate the human genome using functional genomics—e.g., ENCODE *(1)* and GTEx *(2)*—as well as inferring deleterious effects from allele frequencies and location in coding sequence—e.g., gnomAD *(3)* and TOPMed *(4)*. Although these seminal projects greatly expanded knowledge, this “central problem in biology” is unresolved and motivated the NHGRI Impact of Genomic Variation on Function initiative.

Evolutionary constraint is complementary to these efforts. Functional importance is inferred from the signatures of evolution in the human genome: “constraint” indicates genomic positions that have changed more slowly than expected under neutral drift due to purifying selection. A key advantage of constraint lies in its mechanistic agnosticism; a highly constrained base has an impact on some biological process, in some cell, at some life stage (discussed in *Supplementary Methods, Section 2)*. Constraint has been used in efforts to understand the human genome for over 50 years beginning with cross-species protein sequence comparisons. More recently, at the extremes of the allelic spectrum, constraint is often used by clinical geneticists to prioritize potentially causal rare variants *(5, 6)*, and common variants in regions under constraint are highly enriched in genome-wide association study (GWAS) results *(7–9)*. Despite its reported importance, evolutionary constraint is not systematically leveraged in interpreting the function of GWAS loci *(10–15)*.

Our companion paper describes the Zoonomia reference-free alignment of 240 placental mammals spanning ∼100 million years of evolution *(Companion paper #1, Christmas et al*.). The analyses showed the unprecedented informativeness of this alignment at multiple scales: from exceptionally constrained 100 kb bins (e.g., all *HOX* clusters) to smaller ultra-conserved and human accelerated regions, non-coding regulatory regions, nuances of the genetic code, and specific base positions in binding motifs. These results strongly suggested the utility of constraint as a functional annotation that can be leveraged to deepen our understanding of heritable human diseases. In this paper, we demonstrate the importance of mammalian constraint for connecting genotype to phenotype for human disease.

## The properties of evolutionary constraint at single base resolution

### Defining constraint

Placental mammalian constraint was estimated using phyloP scores *(16)* across 240 species for 2,852,623,265 bases in the human genome (chr1-22, X, Y; *Supplementary Methods, Section 3)*. In our companion paper we estimated that ∼13% of the human genome is under some degree of constraint due to purifying selection; for these disease-focused analyses, we used an empirical subset with the strongest constraint signatures. We defined a base as constrained in mammals if its phyloP score was ≥ 2.27 (FDR 0.05 threshold, 100,651,377 bases or 3.53% of the genome). We defined constraint across 43 primates using a phastCons *(17)* threshold (≥ 0.961, 101,134,907 bases) selected to match the same fraction of the genome annotated as constrained in mammals. Mammalian and primate constraint overlapped significantly but not fully (Jaccard index 0.30). In *Supplementary Methods, Section 4*, we describe the properties of constrained genomic positions, from base level to higher order annotations. Briefly, we found that mammalian constrained bases had a marked tendency to cluster (median distance 2 bases) compared to random expectations (median 24 bases), and that specific genomic elements were highly enriched in constrained bases (particularly coding sequence, CDS, as expected) as well as multiple regulatory features (Figs. 1A and S1), and that constraint scores captured nuances of the genetic code (fig. S2).

**Fig. 1.**
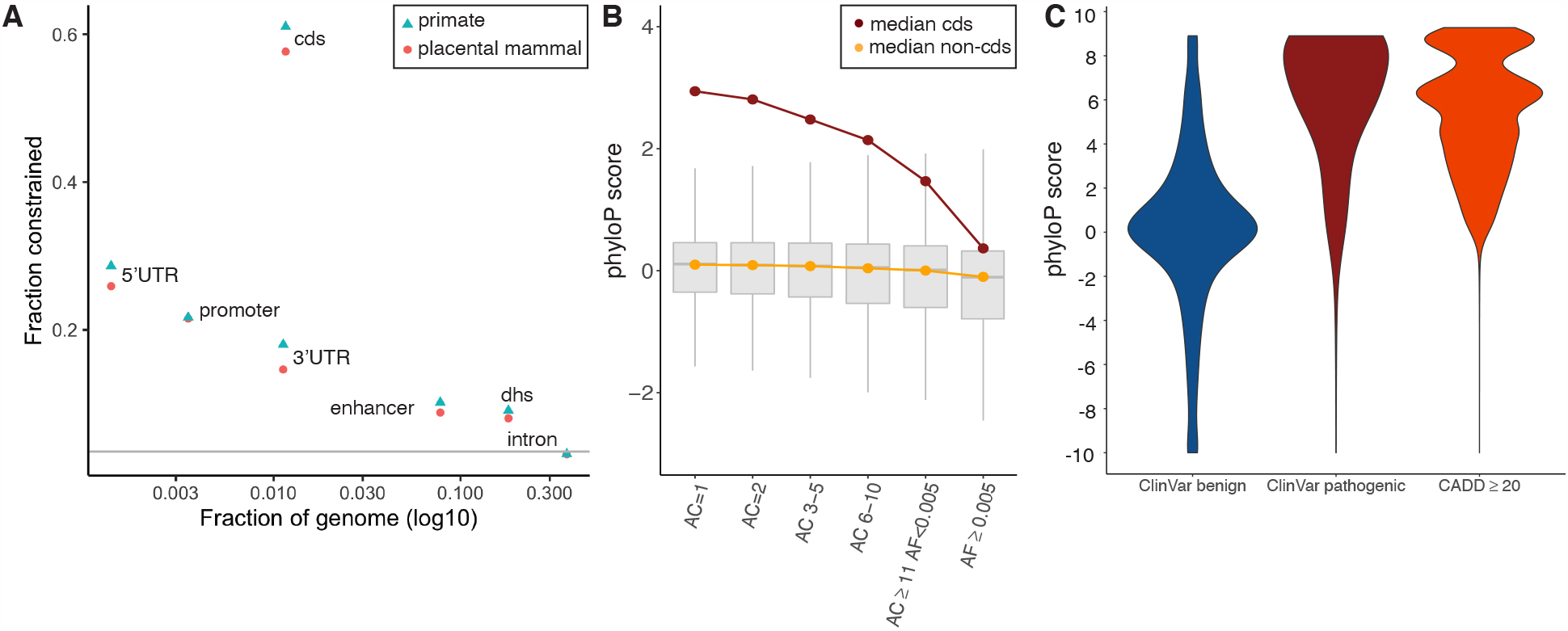
(A) Evolutionary constraint in multiple genomic partitions. X-axis=fraction of the genome occupied by a partition, Y-axis=fraction of partition under constraint in placental mammals (orange circles) and primates (blue triangles), grey line is the genome mean (0.035). The greatest constraint is found in CDS and key regulatory regions (5’UTR, ENCODE promoter-like elements, and 3’UTR). This figure is a subset of fig. S1 which shows more biotypes, protein-coding gene parts, and regulatory regions. (B) Whisker plots of constraint in variants from TOPMed WGS, stratified by CDS (red, 6.14 million biallelic SNPs) and non-CDS variants (orange, 549.64 million biallelic SNPs). X=six allele count (AC) bins, from singletons (AC=1, 44.8%) to common variants (allele frequency, AF ≥ 0.005, 1.4%). (C) PhyloP score density for ClinVar benign (N=231,642), ClinVar pathogenic (N=73,885), and gnomAD WGS variant positions with CADD ≥ 20 (N=3,958,488).

### Constraint across the allelic spectrum

Genetic variation is fundamental to heritable human diseases, disorders, and other traits. We thus evaluated the relationship between allele frequency and constraint (Fig. 1B). Using whole genome sequencing data from over 140K humans (TOPMed, v8) *(4)*, we observed an inverse correlation between allele count and phyloP score (rho = -0.07) with stronger correlations in CDS regions and for non-synonymous variants (rho = -0.12 and -0.18, all *P* < 2.2×10^−308^). As expected due to negative selection, common genetic variants were depleted for constrained bases (1.85% vs. 3.53% expected by chance, *P* < 2.2×10^−308^). However, this relatively high fraction of constrained bases highlights the ability of mammalian constraint to predict deleterious effects across the allele frequency spectrum. To evaluate these relations more formally, genome-wide models contrasting singletons (AC = 1) to common variants (AF ≥ 0.005) found that common variants had lower phyloP scores and a marked increase in CG context (fig. S3, *Supplementary Methods, Section 4)*. Models for CDS SNPs found an inverse association of AC with constraint, and that common SNPs had greater odds of occurring at a C or G base, and tend not to occur in important CDS positions (e.g., codon position 1 or 2, or at bases that could mutate to stop).

### Common constrained SNPs are relevant for human diseases

We conducted additional analyses of common SNPs (AF ≥ 0.005) as these variants are foundational for GWAS *(Supplementary Methods, Section 4)*. Of these 15,777,878 SNPs in TOPMed, 1.85% (N = 291,669) are constrained, far less than genome-wide constraint (3.53%). Our modeling showed that constrained SNPs were 22x more likely to occur in CDS bases, 3x more likely to occur in promoters, and ∼2x more likely to be a “fine-mapped” eQTL-SNP or to occur in open chromatin or an enhancer.

The strong tendency of these constrained SNPs to occur in CDS was unexpected given that (by definition) these positions are highly constrained in placental mammals and yet variable in humans. We hypothesized that this could occur if selection effects were variable across genes (some generate peptide variability whereas others are highly intolerant of CDS variation). We found that 37.8% of protein-coding (PC) genes had no constrained CDS SNPs and other genes had appreciable fractions (up to 10% of all CDS bases are common SNPs). The top 5% (N = 980) of genes with the most constrained CDS SNPs have medical relevance (131 have an OMIM entry including multiple neurological disorders) and were strongly enriched for G-protein coupled receptors (GPCR), “druggable” genes (both GPCR and non-GPCR) *(18)*, taste receptors, skin development, and multiple immune processes. These biological processes are at the interface of a mammal and its environment and allow adaptation to an environmental niche. We suggest that many of these genes could be prioritized for gene-environment interactions searches as constrained variants reaching high frequency in human populations are relevant for human diseases.

### Base pair resolution of deleterious effects

We contrasted constraint scores to metrics used to aid the interpretation of functional variation for human health. First, pathogenic ClinVar *(19)* variants were significantly skewed to higher phyloP in comparison to benign variants (two-tailed Wilcoxon rank sum test, *P* < 2.2×10^−16^, Fig. 1C), and phyloP scores were strongly associated with the improvement in annotations of variants in ClinVar from 2016 to 2021 (e.g. uncertain to benign or to pathogenic; *Supplementary Methods, Section 5)*. For a second metric, CADD *(6)*, which incorporates evolutionary constraint, we found variant positions with a higher likelihood of deleteriousness were also enriched for constrained phyloP scores (two-tailed Wilcoxon rank sum test, *P* < 2.2×10^−16^, Fig. 1C). A focused analysis of human non-synonymous variants at constrained sites across the mammalian tree using TOGA (Tool to infer Orthologs from Genome Alignments, *Companion paper #1, Christmas et al*.; *Companion paper #10, Kirilenko et al)*, identified 1,570 genes for which a non-synonymous change resulted in a ClinVar pathogenic or likely pathogenic phenotype in humans *(Supplementary Methods, Section 5)*. For example, the *CFTR* gene underlying cystic fibrosis *(20)* showed a high burden of pathogenic compared to benign sites (123 vs. 1 out of 1,585 alignment sites). A further 12,889 genes had identifiable constrained sites, but lacked records of non-synonymous pathogenic alterations *(Supplementary Methods, Section 5)*. Several of these constrained positions, currently lacking ClinVar pathogenic annotations, likely represent novel sources of deleterious variation resulting in a disease state. We tested this by leveraging functionally explored variation in two G-protein coupled receptors, *GPR75 (21)* and *ADRB2 (22)*, and showed that functionally important SNP or amino acid sites respectively, were marked by higher constraint scores *(Supplementary Methods, Section 5)*. Species alignments at this scale also allow for the identification of potential model systems, those for which a substitution may result in a human disease state, but is otherwise naturally occurring in non-human mammals. We found 697 such sites across 330 genes, including multiple positions in *SOD1* (pathogenic sites for amyotrophic lateral sclerosis). These observations open the avenue for natural adaptive variants to inform the development of new therapies for treatment *(Supplementary Methods, Section 5)*.

## Common variation and human diseases and complex traits

GWAS have found that the genetic architecture of human diseases and complex traits is highly polygenic and dominated by common variants with weak effects *(10)*. Here, we dissected the impact of common variants (defined in this section as AF ≥ 0.05) on this architecture via polygenic analyses of disease SNP-heritability *(h*^*2*^) using stratified LD score regression (S-LDSC) *(7, 23, 24)* using the results of 63 independent European ancestry GWAS *(25)* (mean N = 314K; table S1, *Supplemental Methods, Section 6)*.

### Constraint scores are proportional to common variant SNP-heritability enrichments

We first validated the relevance of our constraint scores to investigate the role of common variants in human diseases and complex traits. We found that common variants in the highest constraint score percentiles had greater enrichment for GWAS trait associated variants (measured by SNP-*h*^*2*^ enrichment, the proportion of *h*^*2*^ divided by the proportion of SNPs; Fig. 2A and table S2). We observed decreasing but significant enrichments *(P* < 0.05/15) for SNPs in the four first percentiles of mammalian constraint scores (phyloP) (in line with 3.53% of the genome bases being considered as constrained using a 5% FDR threshold), and in the first five percentiles of primate (phastCons) constraint scores. We justified the use of different scores to measure constraint in mammals and primates by the fact that phyloP scores were unable to detect single-base constraint in primates due to lack of power and were too noisy to lead to significant *h*^*2*^ enrichment (fig. S4). While both phyloP and phastCons scores performed similarly in heritability analyses, phyloP is superior for having single-base resolution (fig. S4 and additional justification in *Supplemental Methods, Section 6)*.

**Fig. 2.**
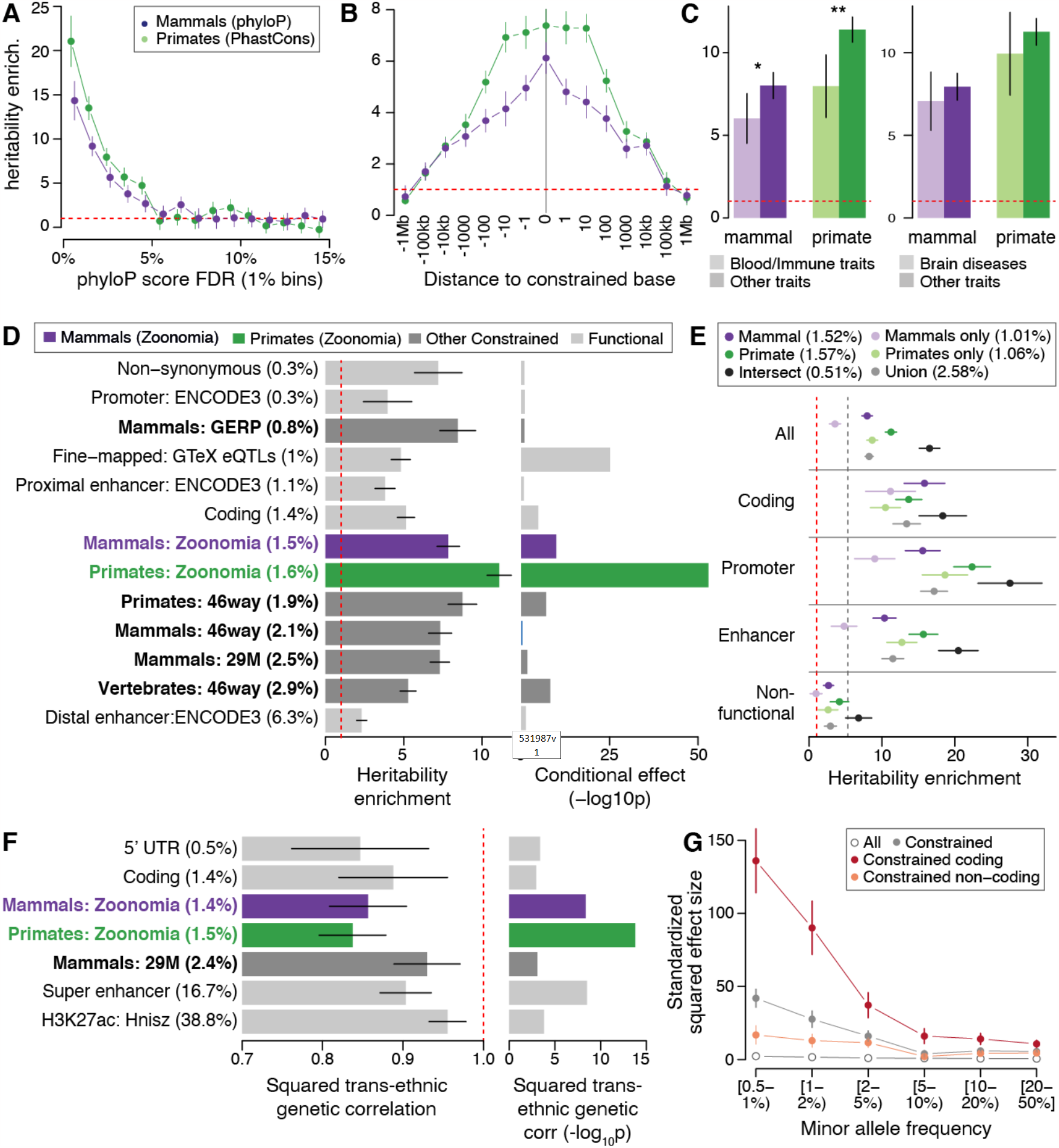
SNP-heritability analyses of variants at constrained positions in human complex traits and diseases. (A) Heritability enrichment of common SNPs in the top percentiles of constraint scores in placental mammals (phyloP) and primates (phastCons). (B) Heritability enrichment as a function of the distance to a constrained base. (C) Heritability enrichment of constrained annotations in 11 blood and immune traits and 9 brain diseases (light color) versus other types of traits (dark color). Asterisks indicate significance at P < 0.05 and double Asterisks indicate significance at P < 0.05 after Bonferroni correction (0.05/4). (D) Heritability enrichment of constrained and functional annotations (left), and corresponding significance of the conditional effect while considered in a joint model with 106 annotations (right). (E) Heritability enrichment of constrained annotations intersected together and stratified by their genomic function. The dashed grey line represents heritability enrichment in coding regions (plotted for comparison purposes). (F) Squared trans-ancestry genetic correlation enrichment (left) with corresponding significance (right) for 7 annotations with significant depletion of squared trans-ancestry genetic correlations. (G) Standardized squared effect sizes as a function of allele frequency. Results are meta-analyzed across 63 independent GWAS (A, B, C, E), 31 independent traits with GWAS available in European and Japanese populations (F), and 27 independent UK Biobank traits (G). Dashed red lines represent a null enrichment of 1 (A-E) and a null squared trans-ancestry genetic correlation (f). Error bars are 95% confidence intervals. Numerical results are reported in tables S2, S3, S4, S6, S7, S8, and S11.

### Mammal constraint scores are base pair specific

We evaluated the resolution of constraint scores by estimating SNP-*h*^*2*^ with different distances to a constrained base. First, we confirmed the base pair resolution of mammalian constraint scores by observing that SNPs ∼1 bp from a constrained variant were significantly less enriched than constrained SNPs *(P* ≤ 3.35×10^−3^) (Fig. 2B and table S3). We also observed log-linear decrease of *h*^*2*^ enrichment as a function of the distance to a constrained base, with significant *h*^*2*^ enrichment up to 100 kb from constrained bases, confirming the larger-scale clustering of constrained bases. Finally, demonstrating the power of a broad placental mammal-wide genome sampling, constraint scores obtained only from primate species have lower resolution (∼10 bp, Fig. 2B) as these are based on fewer species (43), from a single mammalian order, and thus less branch length.

### Zoonomia constraint is uniquely informative

Annotations derived from mammal and primate constrained positions were more informative for human diseases than key functional annotations, including previously published constrained annotations *(17, 26, 27)* (Fig. 2D and table S4). First, their degrees of enrichment (7.84 ± 0.37 fold for mammals and 11.10 ± 0.40 fold for primates) exceeded those of previously published constraint and key functional annotations, such as non-synonymous coding variants (7.20 ± 0.78 fold) or fine-mapped eQTL-SNPs (4.81 ± 0.31 fold) *(28)*. Second, in conditional analyses involving 106 annotations analyzed jointly *(Supplemental Methods, Section 6)*, we observed that these constrained annotations were among the most significant *(P* = 1.17×10^−10^ for mammals, and *P* = 1.19×10^−53^ for primates, respectively), and more significant than previously published constrained annotations (Fig. 2D and table S4).

### Variants at constrained positions are less enriched in blood and immune traits heritability than in other complex traits

We did not observe disease-specific patterns for our constrained annotations, without any trait exhibiting significantly higher *h*^2^ enrichment than the mean calculated for the mammal and primate constrained annotations (fig. S5 and table S5). However, we observed consistently lower *h*^2^ enrichments for constrained annotations in a meta-analysis of 11 blood and immune traits, as previously observed *(7)*, but no differential enrichment in 9 brain disorders (Fig. 2C, table S1, and table S6).

### Variants at positions constrained in primates are informative for non-coding common variants

Surprisingly, SNPs constrained in primates have greater SNP-*h*^2^ enrichment than SNPs constrained in mammals (Figs. 2A-C). To investigate, we intersected mammalian and primate constraint information, and observed significantly higher *h*^2^ enrichment in SNPs constrained in both mammals and primates (16.52 ± 0.73 fold), compared with constraint only in primates (8.66 ± 0.38 fold), or only in mammals (3.56 ± 0.40 fold) (Fig. 2E and table S7). We verified that these results are mostly driven by the intersection of mammal and primate constrained bases (and not due to the different scoring tests, fig. S6). By stratifying constrained mammalian bases by their primate constraint scores, we found that variants identified as constrained in mammals but not in primates are not significantly enriched in *h*^2^, whereas SNPs constrained in primates were significantly enriched regardless of their constraint scores in mammals (fig. S7). These results explain the lower SNP-*h*^2^ for constraint in mammals, and demonstrate increased informativeness when combining information from primates and mammals. Interestingly, we observed consistently higher *h*^2^ enrichment for SNPs that are constrained in both mammals and primates when stratifying by genomic function (i.e., coding regions, promoters, and enhancers), but that constraint is more informative in primates than in mammals only for non-coding variants (Fig. 2E). Strikingly, we observed that constrained SNPs defined as non-functional (see *Supplemental Methods, Section 6)* were still enriched in *h*^2^ (>2.67 fold with *P* < 1.22×10^−4^, except for SNPs constrained only in mammals or primates; Fig. 2E), emphasizing the informativeness of our constrained annotations to annotate non-coding variants with unknown function.

### Disease effect sizes of common variants at constrained positions differ across human populations

While our heritability analyses focused on European ancestry GWAS, variant effect sizes differ across human populations, especially for variants with stronger gene-environment interactions *(29)*. To quantify how effect sizes of constrained common variants differ across populations, we applied S-LDXR *(29)* on 31 diseases and complex traits with GWAS data from East Asian (mean *N* = 90K) and European (mean *N* = 267K) populations. Variants at constrained sites in mammals and primates were among the most significantly depleted in squared trans-ancestry genetic correlation *(P* = 4.38×10^−9^ and *P* = 1.63×10^−14^, the third and most significant investigated annotation, respectively; Fig. 2F and table S8). These results highlight more population-specific causal effect sizes for variants at constrained positions, in line with stronger gene-environment interactions at these loci, and potentially explain how genetic variations at constrained bases could have become common in human populations.

### Strong effect sizes for coding low-frequency variants at constrained positions

Annotations constrained by purifying selection tend to have low-frequency variants (0.5% ≤ AF < 5%) with larger effect sizes leading to higher enrichment in low-frequency variant *h*^2^ compared to common variant *h*^2^ *(8)*. We quantified low-frequency SNP-*h*^2^ enrichments of constrained annotations by analyzing 27 well-powered independent UK Biobank traits (same as in *(8)*; mean *N* = 355K; table S9). We observed that constrained annotations had consistently larger low-frequency *h*^2^ enrichment than common *h*^2^ enrichment, especially for variants at constrained sites in mammals (16.83 ± 0.92 vs. 8.70 ± 0.72 fold; *P* = 3.22×10^−11^ for difference) (fig. S8 and table S10) in line with greater effect sizes as allele frequency decreases (Fig. 2G and table S11). This enrichment difference was driven by coding variants at constrained sites in mammals (48.84 ± 3.10 vs. 19.42 ± 1.91 fold; *P* = 6.36×10^−16^ for difference); we note that the low-frequency *h*^2^ enrichment for these variants was similar to that of non-synonymous variants (40.38 ± 2.40 fold), suggesting that constraint information is as informative as protein change information at the coding level.

In conclusion, we observed that our mammalian constraint scores have unprecedented base pair resolution to investigate common variants in GWAS findings for human complex traits and diseases, are uniquely informative compared to known functional annotations and previously published constraint scores, are even more informative when combined with primate constraint scores, and could be utilized to investigate variants defined as non-functional.

## Leveraging constraint to move from prioritization to function

### Zoonomia constraint scores improve functionally-informed fine-mapping analyses

Based on our heritability results, we expect that our constraint scores will improve functionally-informed fine-mapping of constrained genetic variants associated with common traits. We compared PolyFun *(30)* fine-mapping results obtained with no annotations (non-functional model), with its default set of annotations (baseline-

LF model), and with an augmented baseline-LF annotations containing multiple Zoonomia constrained annotations (baseline-LF+Zoonomia model) on 24 well-powered UK Biobank diseases and complex traits *(30, 31)* (mean *N* = 440K; table S12 and *Supplemental Methods, Section 7)*. We observed significantly *(P* < 1.00×10^−4^) greater posterior inclusion probability (PIP) for variants at constrained sites in mammals and primates when using PolyFun with the baseline-LF+Zoonomia model compared to the non-functional and baseline-LF models (Figs. 3A and 3B). Notably, PolyFun with the baseline-LF+Zoonomia model detected 1,407 variants at constrained sites in mammals fine-mapped with high confidence (PIP > 0.75) across all the UK Biobank traits (32.80% of high confidence fine-mapped variants), against 732 and 1,216 when using the non-functional and baseline-LF and models, respectively (24.50% and 29.67% of high confidence fine-mapped variants, respectively) (fig. S9).

**Fig. 3.**
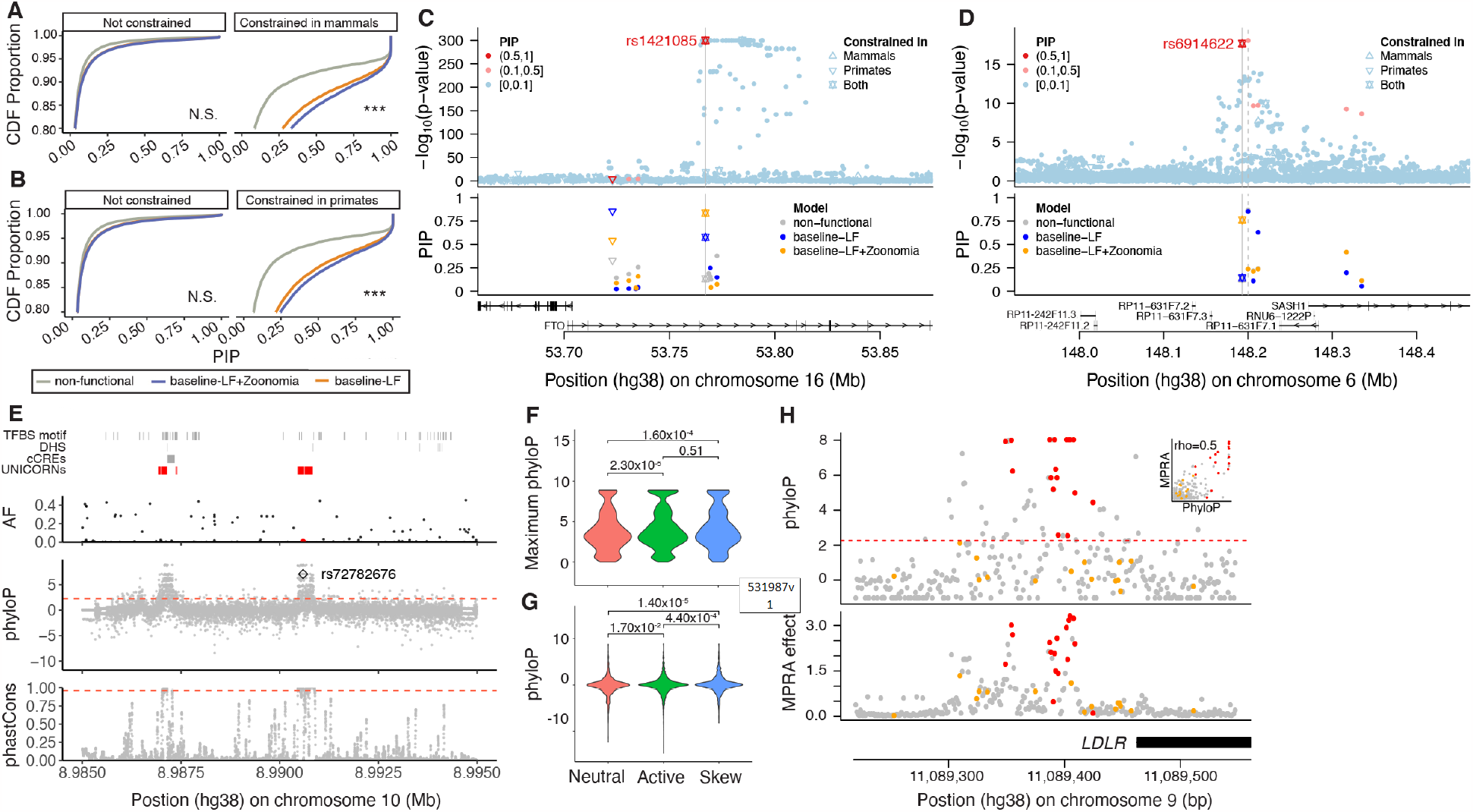
Leveraging constraint to move from prioritization to function. (A,B) We report the cumulative distribution function (CDF) of posterior inclusion probability (PIP) scores using functionally-informed fine-mapping with different models of functional annotations. Distribution functions are split into subpanels by whether the fine-mapped SNP overlaps high constraint scores in mammals (A) and primates (B). One-way Komolgorov-Smirnov tests that CDF for PIP obtained from the baselineLF model (gray) are lower (above) than the CDF for PIP obtained from the baseline-LF+Zoonomia model (orange) with Bonferroni correction for N=4 categories across panels (*** p/N < 0.0001, N.S. not significant). (C,D) Examples of constrained fine-mapped variants. We report GWAS P-values (upper panel) and corresponding PIP under different functionally informed fine-mapping models (lower panel). Shape of the dots corresponds to constraint information. (E) Fine-mapped variants are not limited to the annotated genome as exemplified by rs72782676 (red dot in AF panel) in the GATA3 UNannotated Intergenic COnstraint RegioN (UNICORN) locus. (F,G) Constraint is formally linked to function via massively parallel reporter assays (MPRAs) at the (F) regional oligo and (G) base pair level for neutral, active and allele specific skewed effect. (H) For the LDLR promoter locus, MPRA effect is strongly correlated with phyloP score. Constrained (red), and unconstrained (orange) ClinVar pathogenic variants are plotted to highlight known deleterious positions.

### Fine-mapping examples

We highlight the utility of evolutionary constraint scores in fine-mapping analyses. First, rs1421085 has a causal and experimentally validated association with BMI (the SNP is located in *FTO* but has regulatory effects on *IRX5* and *IRX3) (32, 33)*; this variant is extremely constrained in mammals (phyloP = 6.31) and primates (phastCons = 1.00), leading to a higher PIP when using the baseline-LF+Zoonomia model (0.84) than when using the non-functional and baseline-LF models (0.13 and 0.58, respectively; Fig. 3C). Interestingly, the fraction of CDS and promoter bases that are constrained for *IRX5* (0.79 and 0.58) and *IRX3* (0.74 and 0.34) were higher than for *FTO* (0.61 and 0.23), suggesting that constrained variant in regulatory regions could be more likely to target genes with constrained CDS and/or promoters (see below). Second, rs6914622 is constrained in mammals and primates (phyloP = 2.37 and phastCons = 1.00) and may be causal in hypothyroidism via the baseline-LF+Zoonomia model (PIP = 0.76; Fig. 3D) but not via the non-functional and baseline-LF models (PIP ≤ 0.14). Conversely, the sentinel variant rs9497965 is not evolutionarily constrained but has a notable PIP in the baseline-LF model (PIP ≥ 0.85) but not in the baseline-LF+Zoonomia model (PIP = 0.24). Using epigenetic marks from four thyroid cell types *(34)* (functional information not in the fine-mapping models), rs6914622 was in an active enhancer in all thyroid cell-types and rs9497965 was inferred as being in an enhancer in only one thyroid cell type (weak transcription and quiescent for the others), suggesting a causal role for rs6914622 over rs9497965. While functional follow-up is necessary, these examples illustrated how Zoonomia constraint scores can significantly impact fine-mapping. One method of functional follow-up, Cell-TACIT, is explored in a companion paper *(Companion paper #11, Phan et al*.), in which the conservation of human neural cell type-specific open chromatin across mammals is used to improve the fine-mapping of GWAS for brain disorders. Some regulatory elements may not be conserved at the nucleotide level but lie in a cell type regulatory element predicted to be conserved across mammalians. Fine-mapping genetic variants with constraint and Cell-TACIT provide examples of how mammalian genomes can be leveraged to discover nucleotide and regulatory conservation to link variation to function. Finally, as discussed in another companion paper, Human Accelerated Regions can also improve fine-mapping interpretation *(Companion paper #8, Keough et al*,).

### Measures of constraint can reveal unannotated variants impacting human health

Due to the challenge of generating functional datasets in all cell-types and cell-states, much of the genome’s regulatory space is still not fully annotated *(35)*. The high levels of constraint and low levels of variant diversity in UNannotated Intergenic COnstraint RegioNs (UNICORNs, *Supplemental Methods, Section 8, Companion paper #1, Christmas et al*.) suggest that they are likely of functional importance despite lacking functional annotations (consistent with our observation that non-functional constrained SNPs are enriched in *h*^2^, Fig. 2E). While fewer fine-mapped SNPs were located within UNICORNs (833 SNPs) compared to a matched set of random unannotated non-constrained intergenic regions (5,895 SNPs) and to SNPs located elsewhere in the genome (305,599 SNPs), those variants had higher mean PIP scores (0.15 UNICORNs vs 0.05 for the other two regions). This demonstrates that UNICORNs can reveal unannotated variants impacting human health and disease. UNICORNs contain fine-mapped SNPs with significantly higher PIP scores compared to the background sets across multiple traits (linear regression, *P* < 0.01 in all cases after correcting for multiple testing; table S13). For example, a 163 bp UNICORN contains rs72782676 with fine-mapping evidence for multiple traits (e.g., eosinophil count, asthma, eczema, respiratory and ENT diseases; AF_TOPMed_ = 0.005; PIP > 0.99 in all GWAS) (Fig. 3E). The nearest gene, *GATA3*, sits 915 kb upstream, and is a master transcriptional regulator for T Helper 2 lineage commitment *(36)*, and is known to play an important role in inflammatory disease *(37, 38)*. This UNICORN highlights a strong regulatory candidate for *GATA3* in a disease-relevant region currently lacking annotation.

### Predicted variant effect validated at single base resolution

Massively parallel reporter assays (MPRAs), have been used to rapidly test thousands of genomic variants for their potential regulatory effects on gene expression. While the functional output from these high-throughput methods are useful for localising putative causal alleles, overlaying constraint scores may help further elucidate functional variants *(Supplemental Methods, Section 8)*. To investigate this, we integrated our Zoonomia-derived phyloP scores with > 35,000 assayed variants from existing 3’UTR *(39)* and eQTL *(40)* MPRAs. Using the 3’UTR MPRA data to highlight our results, we found that phyloP scores could differentiate between sequence backgrounds with and without regulatory activity, (e.g. across multiple tissues, Neutral vs Active: *P*_olig_ = 2.32×10^−5^, Fig. 3F). PhyloP scores further highlighted variants with allele-specific regulatory effects (e.g. Neutral vs Skew: *P*_base_ = 1.4×10^−5^; Fig. 3G). Additionally, we found that selection on constrained phyloP positions enriched the allele-specific regulatory effects by 1.3 fold *(Supplemental Methods, Section 8)*. Similar trends were observed in promoter and enhancer saturation mutagenesis MPRAs *(41)*. For example, phyloP constraint was a strong predictor for variant effect within the *LDLR* promoter (Spearman rho = 0.51), with five of the most constrained sites providing the strongest regulatory effects and also tagging pathogenic ClinVar positions (Fig. 3H). Further, in our companion paper *(Companion paper #? CONDEL, Xue et al)*, we use MPRAs to directly assess the regulatory impacts of bases under high constraint that have been deleted specifically in the human lineage. For many we can precisely identify how the deletions impact transcription factor binding which is well correlated with the observed regulatory changes, linking sequence change to mechanism. We found these human-specific deletions were enriched to overlie psychiatric disease GWAS signals (i.e. Schizophrenia, Bipolar), and discovered 717 deletions with significant species-specific regulatory effects, providing candidates targets that may have contributed to the prevalence of human neurological disorders.

### Evolutionary constraint, protein-coding genes, and human disease

Gene-based measures of evolutionary constraint have an important role in understanding the impact of genetic variation on human disease (e.g., LOEUF) *(3)*. As detailed in *Supplementary Methods, Section 9*, we defined 7 measures of gene constraint based on the Zoonomia alignment including fraction of CDS constrained, normalization against 32.13 million CDS bases, a model-based approach adjusting for 12 covariates (codon information, mutational consequences, and positional features), and cross-species amino acid constraint (normalized Shannon entropy). After evaluation, we selected the fraction of constrained CDS bases per gene (fracCdsCons) as a simple measure of gene constraint, given its continuous distribution, low missingness, high correlations with more complex measures of gene constraint, and external validation (Fig. 4A). These gene-based constraint metrics are provided in table S14.

**Fig. 4.**
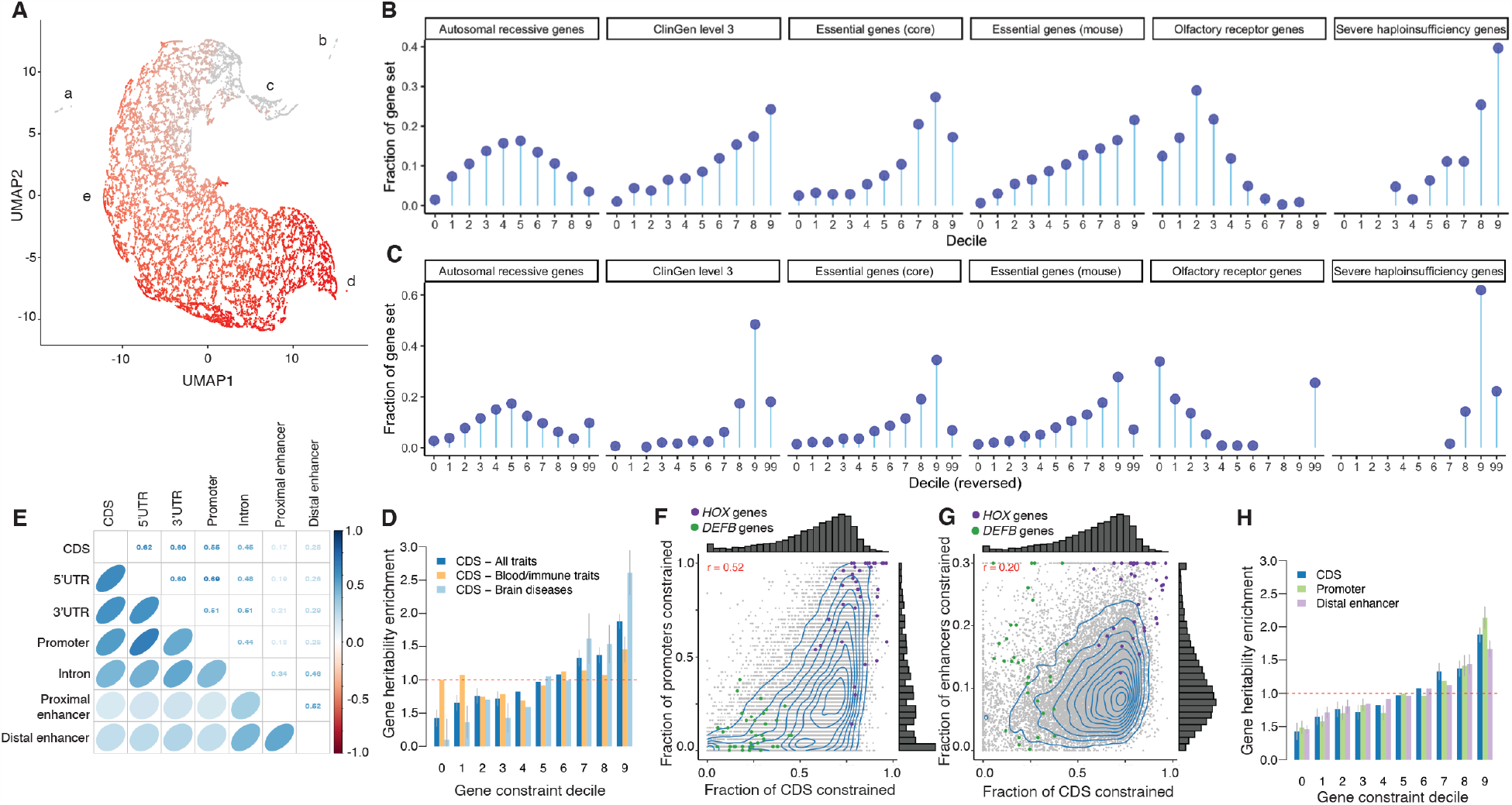
Evolutionary constraint, protein-coding genes, and human disease. (A) Scatterplot of protein-coding (PC) gene clustering (UMAP and DBSCAN). X- and Y-axis are the UMAP coordinates. Each point is a PC gene (N = 19,386). Five clusters are labeled: A = 56 genes whose CDS bases are in complex regions that align poorly; B = 221 genes apparently human-or primate-specific; C=669 genes with good alignment and possible human-specific functions (e.g., five HLA genes and 14 interferon alpha genes); D=15 genes, all highly constrained; and E = all other 18,425 PC genes. Coloring shows fracCdsCons, grey = least and red = most constrained with an anticlockwise gradient in mammalian constraint from upper middle to lower right. (B, C) Gene constraint deciles versus external gene sets as “lollipop plots”. Each panel has 6 subgraphs for autosomal recessive genes, ClinGen level 3 genes, essential genes from Hart, essential genes in mouse, olfactory receptors, and severe haploinsufficiency genes. X-axis = constraint decile (0 = least,, 9=most constrained, 99 = missing). Y-axis = circles are the fraction of the PC genes in a gene set in each decile. (B) Zoonomia fracCdsCons and (C) recapitulates Figure 3 from ref. (3) with LOEUF decile reversed and showing missing data. (D, H) Gene heritability enrichment for SNPs linked to genes of each decile of fracCdsCons (D) and of SNPs linked to genes of decile of constraint in different gene gene features (H). Dashed red lines represent a null enrichment of 1. Error bars are 95% confidence intervals. (E) Spearman correlation of constraint fraction between the parts of PC genes. (F, G) Fraction of CDS constraint (fracCdsCons) vs. fraction or promoter constraint (F) and fraction of distal enhancer constraint (shrinked to values <0.3) (G). Each point is a PC gene, and HOX genes (purple) and defensin beta (DEFB) genes (green) are highlighted.

Given the complexities of human PC genes, it would be surprising if any one gene metric applies to all genes (e.g., LOEUF and pLI are missing for 10.1% of PC genes). We used an empirical approach to identify gene outliers, and identified 277 genes (1.43%) inaccessible to fracCdsCons (clusters A-B, Fig. 4A; *Supplementary Methods, Section 10)*.

We validated fracCdsCons in several ways *(Supplementary Methods, Section 10)*. First, given its widespread use, we compared fracCdsCons to the inverse-scored LOEUF *(3)* and found rho = -0.55. This is notable given the markedly different basis of each measure—constraint over ∼100 million years of mammalian evolution vs statistical modeling of pLoF counts in human WES catalogs *(Supplemental Methods, Section 2)*: empirical confirmation is an important validator for both measures. We next compared fracCdsCons to external gene sets with established patterns of constraint (similar to the LOEUF validation strategy)*(3)* and obtained similar patterns between both scores (Figs. 4B and 4C).

Second, we used an empirical approach to cluster genes based on different constrained metrics (Fig. 4A; *Supplementary Methods, Section 10;* table S14). We identified 277 gene outliers (1.43%) inaccessible to fracCdsCons (clusters A-B), and conducted gene set analyses for 19,109 PC genes (clusters C-E, tables S15 and S16). The 5% most constrained genes (N=955, fracCdsCons 0.811–0.975) were strongly enriched in gene sets: basic embryology (stem cell proliferation/differentiation, tube formation, anterior/posterior patterning, endoderm/mesoderm formation); organ morphogenesis (central/peripheral nervous system, connective tissue, ear, epithelium, eye, gastrointestinal tract, heart, kidney, lung, muscle, myeloid, pancreas, skeleton); cell cycle (phase transition, fate, WNT), cell signaling, positive and negative regulatory processes; and pre-/post-synaptic processes (synapse assembly, postsynaptic density, neurotransmitter regulation, synaptic vesicle cycle, modulation of transsynaptic signaling). The 5% least constrained genes (N=956, fracCdsCons 0–0.150) were strongly enriched in gene sets: microbial defense response (adaptive immunity, bacteria/virus, cell killing, cytokine/interferon); bitter taste and olfaction; and skin development (keratinization, keratinocyte differentiation, epidermal cell differentiation, and epidermis development). The most constrained genes captured processes fundamental to the making of a mammal and the least constrained genes are central to the adaptive evolution of a mammal to its environment—i.e., the specific microbiota, adaptations of smell and taste to detect mates, prey, predators, and poisons, and adaptations of skin for temperature regulation, camouflage, and defense.

Finally, we evaluated the relevance of mammalian gene constraint to human disease. Fig. S10A shows the relationship of fracCdsCons with multiple human disease annotations. For all comparisons, increasing constraint is correlated with increasing relevance for human disease. Fig. S10B depicts the relation with GTEx gene expression, and greater gene constraint is correlated with greater expression in all tissues. “Housekeeping” genes that are uniformly expressed across tissues had greater constraint *(P* < 3×10^−197^) and comprised 3.0% of the least constrained decile and 30.5% of the most constrained decile. Finally, we evaluated the impact of common SNPs linked to PC genes in each fracCdsCons decile by estimating their gene *h*^*2*^ enrichment (defined as *h*^*2*^ enrichment for the decile annotation divided by the mean *h*^*2*^ enrichment over all deciles) using S-LDSC on 63 independent GWAS datasets *(Supplemental Methods, Section 10)*. We observed significantly higher gene *h*^*2*^ enrichment for SNPs linked to genes in the most constrained deciles *(P* = 6.96×10^−59^; Fig. 4D and table S17). Interestingly, we observed stronger gene *h*^*2*^ enrichment patterns in a meta-analysis of nine brain disorders, and gene *h*^*2*^ enrichment patterns nearly independent of gene constraint in a meta-analysis of 11 blood and immune traits (Fig. 4D and table S17).

### Mammalian constraint is correlated between coding and regulatory elements

We extended our approach to measure gene constraint on different regulatory features (including promoters, and ENCODE3 distal enhancers linked to their genes using EpiMap *(34)*), as human diseases and complex traits are predominantly impacted by common regulatory variants. We found substantial correlations of constraint between CDS and the regulatory parts of protein-coding genes, with a higher correlation between CDS and promoter gene constraint *(r* = 0.55) than between CDS and distal enhancer gene constraint *(r* = 0.25) (Figs. 4E-G; gene scores reported in table S18). These correlations are consistent with the idea that if the function of a gene in mammals requires high conservation of protein structure, then its regulatory sequences tend to also be constrained. Interestingly, we observed families of genes with shared constrained patterns (such as *HOX* genes that have constrained exons, promoters and enhancers), and with distinct constrained patterns (such as defensin beta *(DEFB)* genes, which only have constrained enhancers). Finally, we observed that common SNPs linked to genes with constrained promoters and distal enhancers are as enriched in *h*^*2*^ as genes with constrained CDS, suggesting that constraint in regulatory elements can be leveraged in the analyses of human diseases and complex traits (Fig. 4F and table S17).

### Mammalian constraint and copy number variation

Copy number variants (CNVs) are genomic segments that have fewer or more copies compared to a reference genome. CNVs are important drivers of evolution and risk factors for multiple human diseases *(42–44)*. However, CNVs often occur in high repeat/low mappability regions meaning that detecting their presence and significance often carries uncertainty *(45, 46)*. We thus evaluated whether mammalian constraint could help prioritize potentially disease-related CNVs. First, as a qualitative check, we evaluated a pathogenic CNV—a small distal enhancer upstream of *SOX9* with a ClinVar pathogenic annotation as a cause of Pierre Robin sequence—and found that it was highly constrained *(47) (Supplemental Methods, Section 11)*. Second, we evaluated constraint in structural variants (SV) identified in TOPMed *(4)*. We found that singleton (AC=1) SV deletions, inversions, and duplications had similar fractions of constrained bases. However, common (AF ≥ 0.005) SV deletions had far less constraint than SV inversions or duplications. We speculate that singletons are recent mutations relatively unexposed to purifying selection whereas common SV deletions are directly exposed to selection pressures due to the impacts of haploinsufficiency.

Third, these analyses suggest that constrained bases could have utility in CNV prioritization and burden calculations. Given that CNVs are known risk factors for schizophrenia *(48)*, we obtained the CNV call set from the largest published study (21,094 cases, 20,227 controls) *(49)*. After replicating the main analysis, we found that schizophrenia cases had greater CNV constraint burden (the total number of conserved bases impacted by a CNV) compared to controls. The case-control differences were 4-5 logs more significant than two commonly used measures of CNV burden (total number and total bases per person). The improvements were particularly notable for CNV deletions. We suggest that the number of constrained bases impacted by a CNV is a more direct assessment of functional impact—e.g., a large CNV with no constrained bases is less likely to be deleterious than a far smaller CNV that deletes constrained exons, promoters, and/or enhancer elements.

### Cancer driver genes identified with mammalian constraint

Moving from the germline to the somatic genomes, we demonstrated how mammalian constraint in non-coding regions of the genome could be applied to detect candidate cancer driver genes *(Supplementary Methods, Section 12)*. Non-coding constraint mutations (NCCMs, phyloP ≥ 1.2 *(50)*) were identified using whole genome sequencing data (International Cancer Genome Consortium) *(51)* for two types of brain tumors primarily affecting children. Pilocytic astrocytoma is a low-grade tumor *(52)* and medulloblastomas are malignant brain tumors with intertumoral heterogeneity informed by subgroups determined by molecular profiling (i.e., Wingless/Integrated (WNT), Sonic Hedgehog Signaling (SHH), Group 3 and Group 4) *(53)*. We identified NCCMs within introns, 5’and 3’UTRs, and regions within 100kb of each gene *(50)*.

We found drastically different NCCM rates between the two cancers. In pilocytic astrocytoma, known to have coding/translocation mutations primarily in *BRAF*, high NCCM rates were restricted to the *BRAF* locus, in line with the low somatic mutation burden of this tumor type. Strikingly, for medulloblastoma, 114 genes had ≥ 2 NCCMs/100 kb (Fig. 5A) and 525 genes had ≥ 5 NCCMs per gene. These genes were enriched for the GO biological processes “nervous system development” *(P* = 1.32×10^−26^) and “generation of neurons” *(P* = 1.68×10^−22.^). Among the top 114 genes, 15 gene loci were primarily seen in adult cases (≥18 years of age) and 7 loci in pediatric cases (<18 years of age). A subset of these loci is shown in Fig. 5B *(Companion paper #12, Sakthikumar et al)*. An example is *ZFHX4*, previously reported to be differentially expressed in medulloblastoma *(54)*, where NCCMs were predominantly identified in adult patients of the SHH subgroup, and found in high constraint *ZFHX4* intronic regions (Fig. 5C). For the pediatric set of medulloblastoma, potential driver genes included *BMP4* and the *HOXB* locus (containing multiple genes), mostly in patients diagnosed as Group 3 or Group 4. Multiple NCCMs in these two loci were shown to have differential DNA binding capacity in a medulloblastoma cell line *(Companion paper #12, Sakthikumar et al)*. Further, we noted differential gene expression in medulloblastoma compared to cerebellum for multiple NCCM genes, e.g. *HOXB2 (55)*, for which expression levels correlate with patient survival *(56)*.

**Fig. 5.**
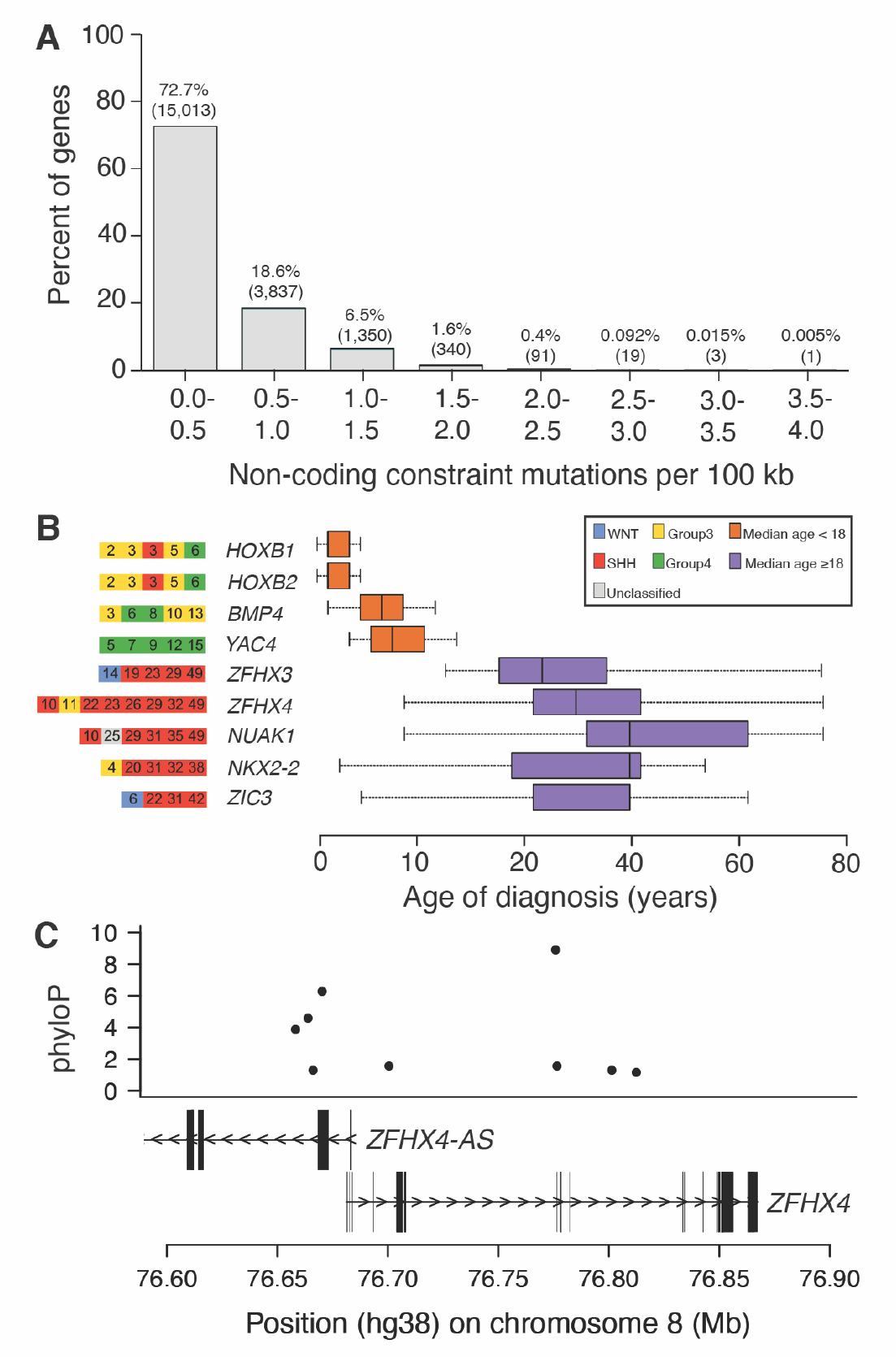
Cancer driver genes identified using NCCM rates. (A) Distribution of the rates of NCCM for medulloblastoma. (B) An example set of the candidate driver genes found either in pediatric (orange) or adult (purple) samples. Age of diagnosis (years) of the patient is indicated together with the tumor subgroup. (C) ZFHX4 locus contains 9 NCCMs drawn from 8 patients.

The addition of evolutionary constraint measures may help advance stratification of medulloblastoma, both with regard to age, and molecular subgroups. More generally, we demonstrate how NCCM analysis can be used as a tool for the identification of novel driver genes in cancer. We suggest that NCCM analysis should be evaluated in more cancer types for its potential to yield a better understanding of disease biology and improved diagnosis and prognosis.

## Discussion

The strength of evolutionary constraint can deepen our understanding of human diseases. The alignment of 240 placental mammals, representing ∼100 million years of evolution, achieved single base resolution that allows detailed evaluation of individual mutations in contrast to previous methodologies of only gene-sized resolution. Evolutionary constraint compares favourably to huge amounts of functional genomics data as functionality in any tissue at any time point will be detected by constraint. We demonstrate that constraint can be used to detect candidate causal mutations in both rare and common disease as well as in cancer, and could be particularly leveraged for brain diseases that are more impacted by constrained genes and biological processes. Finally, we note that primate constraint has a stronger heritability enrichment than when measured across placental mammals in non-coding regions suggesting that sequencing more primates would complement the current efforts to validate function of the multitude of regulatory elements present in the human lineage.

## Acknowledgments

Computations and data handling were enabled by resources in projects, SNIC 2017/7-385, SNIC 2017/7-386, SNIC 2019/3-415, SNIC 2019/30-57, SNIC 2019/8-369, SNIC 2021/2-11, SNIC 2021/5-296, SNIC 2021/6-208, SNIC 2021/5-28, provided by the Swedish National Infrastructure for Computing (SNIC) at UPPMAX, partially funded by the Swedish Research Council through grant agreement no. 2018-05973.

## Diversity & Inclusion

One or more of the authors of this paper self-identifies as a member of the LGBTQ+ community.

## Consortium list

Gregory Andrews^1^, Joel C. Armstrong^2^, Matteo Bianchi^3^, Bruce W. Birren^4^, Kevin R Bredemeyer^5^, Ana M Breit^6^, Matthew J Christmas^3^, Hiram Clawson^2^, Joana Damas^7^, Mark Diekhans^2^, Michael X. Dong^3^, Eduardo Eizirik^8^, Kaili Fan^1^, Cornelia Fanter^9^, Nicole M. Foley^5^, Karin Forsberg-Nilsson^10^, Carlos J. Garcia^11^, John Gatesy^12^, Steven Gazal^13^, Diane P. Genereux^4^, Daniel Goodman^14^, Linda Goodman^15^, Jenna Grimshaw^11^, Michaela K. Halsey^11^, Andrew J Harris^5^, Glenn Hickey^16^, Michael Hiller^17,18,19^, Allyson G. Hindle^9^, Robert M. Hubley^20^, Laura Huckins21, Graham M. Hughes^22^, Jeremy Johnson^4^, David Juan^23^, Irene M. Kaplow^24,25^, Elinor

K. Karlsson^1,4^, Kathleen C. Keough^26,27^, Bogdan Kirilenko^17,18.19^, Klaus-Pieter Koepfli^28,29,30^, Jennifer M. Korstian^11^, Sergey V. Kozyrev^3^, Alyssa J. Lawler^31^, Colleen Lawless^22^, Danielle L. Levesque^6^, Harris A. Lewin ^7,32,33^, Xue Li^1,4^, Yun Li 34, Abigail Lind^26,27^, Kerstin Lindblad-Toh^3,4^, Voichita D. Marinescu^3^, Tomas Marques-Bonet^23,35,36,37^, Victor C Mason^38^, Jennifer R. S. Meadows^3^, Jill E. Moore^1^, Diana D. Moreno-Santillan^11^, Kathleen M. Morrill^1,4^, Gerard Muntané^23^, William J Murphy^5^, Arcadi Navarro^23,39,40,41^, Martin Nweeia^42,43,44,45^, Austin Osmanski^11^, Benedict Paten^2^, Nicole S. Paulat^11^, Eric Pederson^3^, Andreas R. Pfenning^24,25^, BaDoi N. Phan^24^, Katherine S. Pollard^26,27,46^, Kavya Prasad^24^, Henry Pratt^1^, David A. Ray^11^, Jeb Rosen^20^, Irina Ruf ^47^, Louise Ryan^22^, Oliver A. Ryder^48,49^, Daniel Schäffer^24^, Aitor Serres^23^, Beth Shapiro^50,51,^ Arian F. A. Smit^20^, Mark Springer^52^, Chaitanya Srinivasan^24^, Cynthia Steiner^53^, Jessica M. Storer^20^, Patrick F. Sullivan^34,54^, Kevin A. M. Sullivan^10^, Quan Sun^34^, Elisabeth Sundström^3^, Megan A Supple^51^, Ross Swofford^4^, Jin Szatkiewicz^34^, Joy-El Talbot^55^, Emma Teeling^22^, Jason Turner-Maier^4^, Alejandro Valenzuela^23^, Franziska Wagner^47,56^, Ola Wallerman^3^, Chao Wang^3^, Juehan Wang^13^, Jia Wen ^34^, Zhiping Weng^1^, Aryn P. Wilder^48^, Morgan E. Wirthlin^24,25^, Shuyang Yao^54^, Xiaomeng Zhang^24^

^1^Program in Bioinformatics and Integrative Biology; University of Massachusetts Chan Medical School, Worcester, MA 01605, USA.

^2^Genomics Institute, UC Santa Cruz, 1156 High Street, Santa Cruz, CA 95064, USA.

^3^Science for Life Laboratory, Department of Medical Biochemistry and Microbiology; Uppsala University, Uppsala, 75132, Sweden.

^4^Broad Institute of MIT and Harvard, Cambridge MA 02139, USA.

^5^Veterinary Integrative Biosciences; Texas A&M University, College Station, TX 77843, USA.

^6^School of Biology and Ecology; University of Maine, Orono, Maine 04469, USA.

^7^The Genome Center; University of California Davis, Davis, CA 95616, USA.

^8^School of Health and Life Sciences; Pontifical Catholic University of Rio Grande do Sul, Porto Alegre, 90619-900, Brazil.

^9^School of Life Sciences; University of Nevada Las Vegas, Las Vegas, NV 89154, USA.

^10^Department of Immunology, Genetics and Pathology, Science for Life Laboratory; Uppsala University, Uppsala, 751 85, Sweden.

^11^Department of Biological Sciences; Texas Tech University, Lubbock, TX 79409, USA.

^12^Division of Vertebrate Zoology; American Museum of Natural History, New York, NY 10024, USA.

^13^Keck School of Medicine; University of Southern California, Los Angeles, CA 90033, USA.

^14^University of California San Francisco, San Francisco, CA 94143 USA.

^15^Fauna Bio Inc., Emeryville, CA 94608, USA.

^16^Baskin School of Engineering; University of California Santa Cruz, Santa Cruz, CA 95064, USA.

^17^LOEWE Centre for Translational Biodiversity Genomics, 60325 Frankfurt, Germany.

^18^Senckenberg Research Institute, 60325 Frankfurt, Germany.

^19^Faculty of Biosciences; Goethe-University, 60438 Frankfurt, Germany.

^20^Institute for Systems Biology, Seattle, WA 98109, USA.

^21^Department of Psychiatry; Icahn School of Medicine at Mount Sinai, New York, NY, USA.

^22^School of Biology and Environmental Science; University College Dublin, Belfield, Dublin 4, Ireland.

^23^Institute of Evolutionary Biology (UPF-CSIC), Department of Experimental and Health Sciences; Universitat Pompeu Fabra, Barcelona, 08003, Spain.

^24^Department of Computational Biology, School of Computer Science; Carnegie Mellon University, Pittsburgh, PA 15213, USA.

^25^Neuroscience Institute;, Carnegie Mellon University, Pittsburgh, PA 15213, USA.

^26^Gladstone Institutes, San Francisco, CA 94158, USA.

^27^Department of Epidemiology & Biostatistics; University of California, San Francisco, CA 94158, USA.

^28^Smithsonian-Mason School of Conservation; George Mason University, Front Royal, VA 22630, USA.

^29^Smithsonian Conservation Biology Institute, Center for Species Survival, National Zoological Park, Washington, D.C., 20008, USA.

^30^Computer Technologies Laboratory; ITMO University, St. Petersburg 197101, Russia.

^31^Department of Biology; Carnegie Mellon University, Pittsburgh, PA 15213, USA.

^32^Department of Evolution and Ecology, University of California, Davis, CA 95616, USA.

^33^John Muir Institute for the Environment; University of California, Davis, CA 95616, USA.

^34^Department of Genetics; University of North Carolina Medical School, Chapel Hill, NC 27599, USA.

^35^Catalan Institution of Research and Advanced Studies (ICREA), 08010, Barcelona, Spain.

^36^CNAG-CRG, Centre for Genomic Regulation; Barcelona Institute of Science and Technology (BIST), 08036, Barcelona, Spain.

^37^Institut Català de Paleontologia Miquel Crusafont; Universitat Autònoma de Barcelona, 08193, Cerdanyola del Vallès, Barcelona, Spain.

^38^Institute of Cell Biology; University of Bern, 3012 Bern, Switzerland.

^39^Catalan Institution of Research and Advanced Studies (ICREA), 08010, Barcelona, Spain.

^40^CRG, Centre for Genomic Regulation; Barcelona Institute of Science and Technology (BIST), 08036, Barcelona, Spain.

^41^BarcelonaBeta Brain Research Center; Pasqual Maragall Foundation, Barcelona, 08005 Spain.

^42^Narwhal Genome Initiative, Department of Restorative Dentistry and Biomaterials Sciences; Harvard School of Dental Medicine, Boston, MA 02115, USA.

^43^Department of Comprehensive Care, School of Dental Medicine; Case Western Reserve University, Cleveland, OH 44106, USA.

^44^Department of Vertebrate Zoology; Smithsonian Institution, Washington, DC 20002, USA.

^45^Department of Vertebrate Zoology; Canadian Museum of Nature, Ottawa, Ontario K2P 2R1, Canada.

^46^Chan Zuckerberg Biohub, San Francisco, CA 94158, USA.

^47^Division of Messel Research and Mammalogy; Senckenberg Research Institute and Natural History Museum Frankfurt, 60325 Frankfurt am Main, Germany.

^48^Conservation Genetics, San Diego Zoo Wildlife Alliance, Escondido, CA 92027, USA.

^49^Department of Evolution, Behavior and Ecology, Division of Biology; University of California, San Diego, La Jolla, CA 92039 USA.

^50^Howard Hughes Medical Institute; University of California Santa Cruz, Santa Cruz, CA 95064, USA. ^51^Department of Ecology and Evolutionary Biology; University of California Santa Cruz, Santa Cruz, CA 95064, USA.

^52^Department of Evolution, Ecology and Organismal Biology; University of California, Riverside, CA 92521, USA.

^53^Conservation Science Wildlife Health, San Diego Zoo Wildlife Alliance, Escondido CA 92027, USA.

^54^Department of Medical Epidemiology and Biostatistics; Karolinska Institutet, Stockholm, Sweden.

^55^Iris Data Solutions, LLC, Orono, ME 04473, USA.

^56^Museum of Zoology, Senckenberg Natural History Collections Dresden, 01109 Dresden, Germany.

