## Supplementary information for "Leveraging Base Pair Mammalian Constraint to Understand Genetic Variation and Human Disease"

This document contains supplementary information (Methods, Figures, Tables, and References). Note that the *Supplementary Methods sections* contain essays describing the approaches used in this work. These are provided to enable reproducible research, and to document, explain, and justify our analyses.

### Supplementary Methods

This section contains essays that provide detail regarding the approaches taken in this work. These generally have a 1:1 connection to a section of the main manuscript. This information is intended to facilitate reproducible research and to document and justify the decisions made in these analyses. All figures called in the methods have the format *Fig. SX.Y*, where *X* is the section number, and *Y* is the Figure number within section *X*.

#### *Section 1: Brief Summary of Current Genomic Knowledge*

By: Patrick Sullivan

We describe here the latest efforts and available resources to catalog and predict the effect of genetic variation on human phenotypes including: common and rare variant association studies and catalogs of human variation and control of gene expression.

**Genome-wide Association Studies (GWAS).** We downloaded the NHGRI/EMB GWAS catalog (10/2021) (*1*). The *inline table* summarizes the results, and *Fig. S1.1A* shows the publication of GWAS over time and *Fig. S1.1B* shows the cumulative genome-wide significant associations by publication date. As noted below, over 95% of the associated SNPs are non-coding.

| *Feature* | *Number* |
|---|---|
| Studies | 4,593 |
| Traits studied | 3,908 |
| Significant associations ($P \leq 5e\text{-}8$) | 2,763 different traits<br>107,685 unique SNPs<br>156,556 total SNP-trait associations |
| Overlap with coding regions | Of 107,685 unique SNPs, only 4.19% or 4,511 were in a coding region |

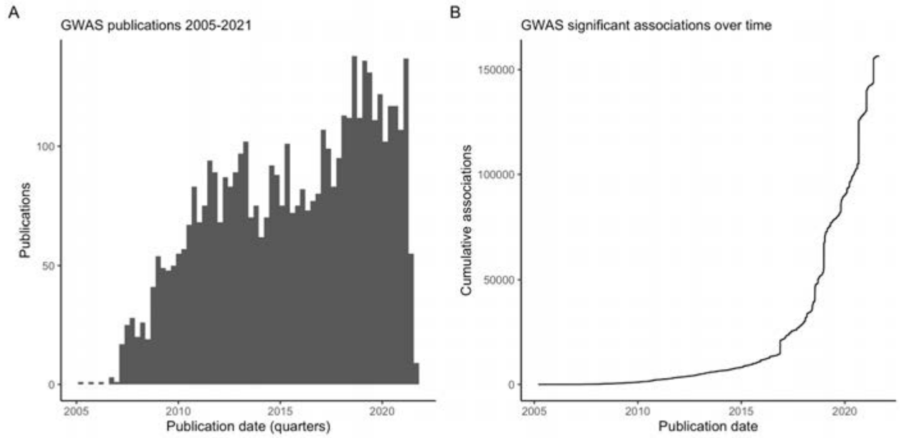

*Fig. S1.1. (A) GWAS publications by publication date by quarter of year. (B) Cumulative number of genome-wide significant SNP-trait associations over time, showing a marked increase in findings since 2017.*

**Genotype-Tissue Expression (GTEx)** has provided a large set of gene expression data in multiple tissues (*2*). Pairing individual gene expression with genomics—in essence, conducting GWAS for SNPs in the vicinity of every transcript—yields a catalog of eQTL-SNP/eQTL-gene pairs. The *inline table* summarizes the results. There are millions of eQTL-SNPs in the genome. Across all tissues, most protein-coding genes have one or more eQTL-SNPs (90.9%, or 18,123/19,937). *Fig1 S1.2* shows the distribution of the number of eQTL-SNPs for each protein coding gene; due to extensive LD in the human genome, most genes have many eQTL-SNPs (*3*).

| *Feature* | *Number* |
|---|---|
| eQTL-SNP/gene pairs | 65,825,763 (all genes, any tissue)<br>41,052,866 (protein-coding genes, any tissue) |
| eQTL SNPs<br>(with ≥1 eQTL-gene) | 4,230,332 (all genes, any tissue)<br>3,552,048 (protein-coding genes, any tissue) |
| eQTL-genes | 34,409 genes with ≥1 eQTL-SNP<br>18,123 protein-coding genes with ≥1 eQTL-SNP |
| eQTL-SNPs per gene | Median 287, IQR 102-550, range 0-10,536 (19,937 genes)<br>(all protein-coding genes, collapsed across tissues, 1,814 genes have none) |
| "Fine-mapped" eQTL-SNPs | Distinct eQTL-SNPs in any GTEx tissue with posterior probabilities ≥ 0.8 in all three credible set algorithms: 0.14% of all eQTL-SNPs |

*GTEx v8, considering all eQTL-SNP/gene pairs with q<0.05. GENCODE gene models. An eQTL ("eQTL-SNP/gene pair") is a significant relationship between a genetic variant ("eQTL-SNP") and the measured expression level of a gene transcript ("eQTL-gene"). Genes with >10,000 eQTL-SNPs were all in the MHC region (NOTCH4, C4A, HLA-DQB1, HLA-DQB2, and HLA-A), due to the exceptionally high LD in this region.*

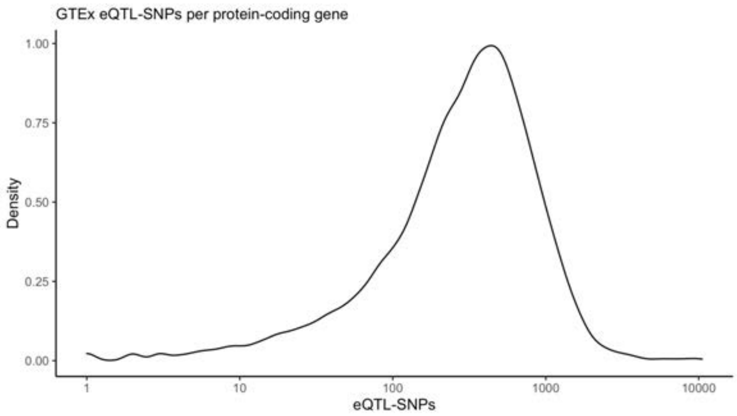

*Fig. S1.2. GTEx v8, density plot of the number of eQTL-SNPs per protein coding gene (18,123 genes). Due to LD, most genes have many eQTL-SNPs.*

In most instances, GWAS and eQTL studies have not directly identified underlying causal variation. Linkage disequilibrium is a double-edged sword. These studies usually genotype a selected set of ~500K biallelic SNPs using a commercial SNP array and then leverage linkage disequilibrium to accurately impute ~8 million variants. However, once a genomic region is identified, linkage disequilibrium usually makes confident identification of causal variation within an associated locus difficult. *Fig. S1.3* depicts a strong association for schizophrenia over a ~1 Mb multigenic region.

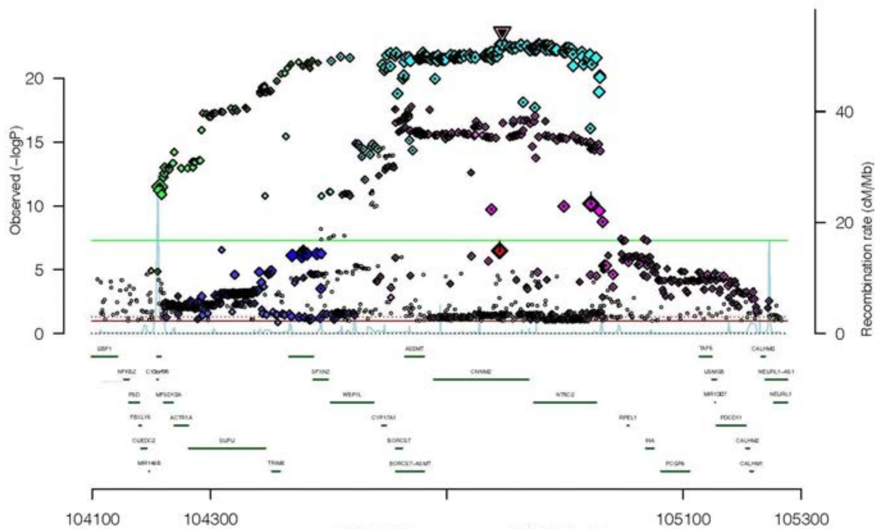

*Fig. S1.3. Locus zoom plot of a genomic region on chr10 strongly associated with schizophrenia. X-axis=hg19 location (in kb). Left Y-axis=association P-value (-$log_{10}$), and the green line is genome-wide significance (P=5e-8). Right Y-axis=local recombination rate. Each point is one SNP. Many SNPs exceed the significance level by 1000x or more. The region is bounded by locally higher recombination rates that are at least partly responsible for high linkage disequilibrium in the region. The associated SNPs are located across 15 genes. This region also contains significant GWAS associations for many other traits, including major depressive disorder, IQ, personality traits, substance use, cardiovascular disease, etc.*

**Single gene diseases.** Online Mendelian Inheritance in Man (OMIM) is a curated source of gene-level findings from human genetics that accumulates human disease gene findings (*4*). The numbers of genes with phenotype-causing mutations was slightly over 1,000 in 2001 and 4,525 in 10/2021. Another source is the variant-level database ClinVar (*5*) which listed 20,493 "pathogenic" variants in 2016 and 85,320 in 10/2021.

**Whole genome sequencing (WGS).** The NHLBI Trans-Omics for Precision Medicine (TOPMed) (*6*) program is conducting WGS on progressively larger samples. *Fig. S1.4A* shows the increase in the number of variants discovered from Freeze 1 in 10/2015 (87 million variants, N=2,643) to Freeze 10b in 4/2021 (1.07 billion variants, N=184,878) as the number of samples and their global diversity increased. *Fig. S1.4B* shows the allele frequency distribution of biallelic SNPs in TOPMed (v8). The rarest variants dominate: 84.8% of biallelic SNPs are in the rarest categories (allele counts ≤ 10, allele frequencies ≤ 3.56e-5), and only 2.6% are common (allele frequency ≥ 0.005).

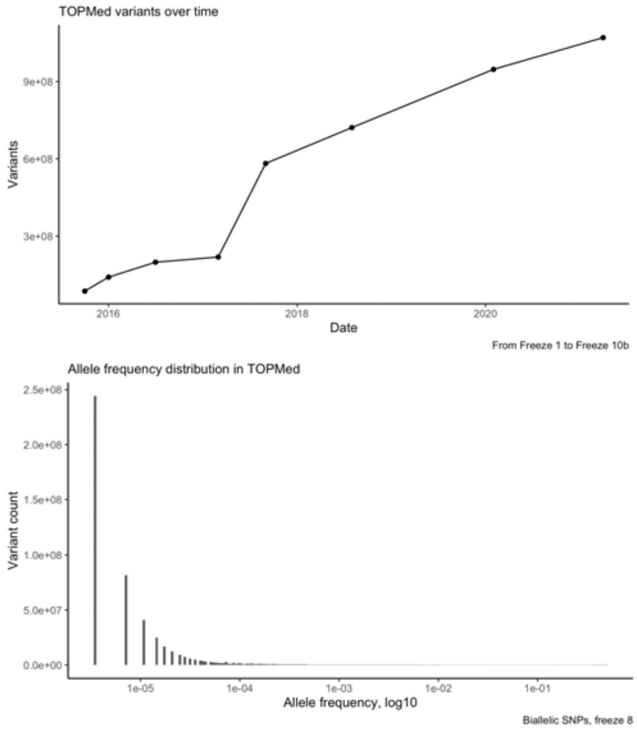

*Fig. S1.4. (A) Marked growth in the TOPMed variant catalog from 2015 to 2021. (B) Allele frequency distribution in TOPMed WGS catalog (V8, N=140,306). The leftmost vertical bars correspond to allele counts of 1, 2, 3, etc. Variants with allele count of 1 account for 46.2% of all variants.*

**Whole exome sequencing (WES).** In 2015, the Exome Aggregation Consortium (ExAC) cataloged 7.4 million variants from WES in 60,706 human subjects along with 179,774 protein-truncating variants (*7*). In 2020, the Genome Aggregation Database (gnomAD) cataloged 14.9 million variants from WES in 125,748 individuals, including 443,769 predicted loss-of-function variants (pLoF) (*8*).

*Fig. S1.5* shows the allele frequency distribution of biallelic SNPs in gnomAD. The rarest variants dominate: 87.9% of biallelic SNPs are in the rarest categories (allele counts ≤ 10, allele frequencies ≤ 3.98e-5), and only 1.2% are common (allele frequency ≥ 0.005). The pLoF subset is defined by mutations that occur at canonical intronic splice sites, frameshift insertion-deletion variants, or introduction of a premature stop codon. However, the numbers of pLoF variants are far less than missense and synonymous variants. Reasonable prediction of the fraction of missense variants that alter protein function is an unsolved problem.

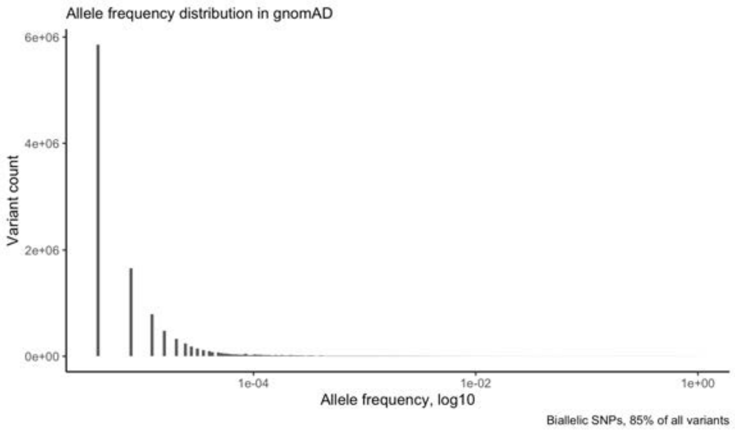

*Fig. S1.5. Allele frequency distribution in the gnomAD exon-focused variant catalog. The leftmost vertical bars correspond to allele counts of 1, 2, 3, etc. Variants with allele count of 1 account for 52% of all variants, and common variants (AF ≥ 0.005) are only 1.2% of the total.*

**Summary.** These data sets document the marked increase in genomic knowledge in the recent past. For the approximately 200,000 genomic positions associated with human rare and common variant studies (i.e., combining the GWAS catalog SNPs and ClinVar "pathogenic" variants), the functional importance or impact of most is unclear. Most GWAS findings are non-coding, and a major hypothesis is that they directly or indirectly point at regulatory variants. The ~200K cataloged associations in the human genome from WES and WGS have greatly exceeded knowledge of their potential functional consequences (particularly in non-coding regions). Even for strong eQTL-SNP to eQTL-gene expression associations—the RNA phenotype "closest" to DNA variation—linkage disequilibrium generates many associations, and there is a clear need for ways to prioritize the observed associations to understand which are more likely to be causal. Taken together, these "fine-mapping" and functional interpretation of non-coding variation problems have motivated the NIH/NHGRI Impact of Genomic Variation on Function (IGVF) consortium ($185 million, 25 awards across 30 sites).

*Section 2: Comparison of Zoonomia Constraint with Human Surveys*

By: Patrick Sullivan, Steven Gazal, Jennifer R. S. Meadows

The *inline table* below compares high level features of mammalian constraint to constraint measures derived from large catalogs of human genetic variation. As noted in *Supplemental Methods, Section 1*, the most comprehensive human catalogs to date are gnomAD (based on WES) and TOPMed (based on WGS) (*6*, *8*). These approaches share the intention of inferring the impact of purifying selection via observed constraint. Beyond this similarity, the methods have sets of strengths and weaknesses.

| *Feature* | *Zoonomia constraint* | *Catalogs of variation in extant humans* [A] |
|---|---|---|
| Basis | Intolerance to variation over ~100 million years of mammalian evolution | Intolerance to variation within a sample of extant humans |
| Resolution | Base pair | Currently, gene- and possibly coding exon-level (*9*). Base pair resolution requires large expansion in sample sizes |
| Evaluable regions | Genomic positions with sufficient numbers of mammalian species aligning. Will miss complex regions and genes with human- or primate-specific functions (e.g., *ZNF588*) (*10*) [B] | In principle, a strength is the capacity to evaluate most genes (including primate- or human-specific) and genomic regions (including complex variants) inaccessible to species comparisons. Exonic frameshifts via insertion-deletion variants can be detected but not generally evaluable via cross-species comparisons § |
| Exposure to purifying selection | Relatively extensive: multiple individual organisms, species, genomic backgrounds, environmental contexts, and sexes. Reproductive fitness is a strength. | Relatively limited: ultra-rare variants exposed for tens of person-years in few individuals, genomic backgrounds, sets of environmental exposures, and sexes. Effects on reproductive fitness and late-onset diseases often unknown. |
| Inference of function | Agnostic to functional mechanism | Requires algorithmic prioritization to identify impactful variation. Current expression quantitative trait loci (eQTL) analyses and epigenomic surveys are incomplete in samples of cell types over developmental time. eQTL analyses are complicated by cell mixtures, gene co-expression, and linkage disequilibrium to highlight causal variant-gene pairs (*3*). |
| Common variation | Identify constrained common variants that are nonetheless variable in humans † | Limited capacity |
| Future expansion | Relatively inexpensive to add 100s to 1000s of additional species | Relatively expensive to add WES or WGS for >100,000 of additional humans |
| Ascertainment | Sensitive to choice of species. DNA may be unavailable (extinct, endangered, or impractical to obtain) | Sensitive to choice of ancestry. Cohorts may directly or indirectly exclude humans with marked physical/cognitive disabilities and early mortality ‡ |

*(A) The phrase "extant humans" is used to distinguish surveys of living/recently deceased humans from genomic studies of ancient humans, and to highlight ascertainment features of the cohorts that comprise studies like gnomAD or TOPMed. (B) Interestingly, although ZNF588 appears to have human-specific functions in brain development, it is not constrained in gnomAD (pLI=0) or, as expected, in Zoonomia.*

† Of 552 million SNPs in TOPMed (v8, chr1-22, chrX), 14,091,740 are common (allele frequency ≥ 0.005). Of the common SNPs, 1.8% (252,080) are constrained. Of the constrained common SNPs, only 9.9%

(25,035) are in CDS; of the constrained, common, CDS SNPs, only 10.5% (2,621) occur at a synonymous site. Constrained common SNPs are a particularly interesting subset given that they are under evolutionary constraint in mammals but are commonly variable in extant humans. They may play roles in important but variable human phenotypes, and may be a priority SNP set for gene-environment interactions (see *Supplementary Methods, section 4*).

‡ For most cohorts in human variation catalogs, subjects had to survive to adulthood, provide informed consent, and meet inclusion/exclusion criteria. An unknown number of individuals with strongly deleterious mutations would not be in these studies due to censoring—e.g., early mortality at some point between fertilization and eligibility for inclusion in a human sequencing catalog. Additional individuals would not be included due to some physical disability or illness (i.e., not "healthy" controls) or because of cognitive impairment (that might complicate or make impossible provision of informed consent for participation). Some of the information in these unobserved variants can be captured in gene-based measures of constraint (i.e., fewer observed deleterious variants in a gene compared to a model expectation), but this will generally occur in genes with high levels of constraint and may be missed if the variant occurs in a non-constrained segment of a constrained gene or in a constrained segment of a non-constrained gene. Direct measurement and association of these rare variants with disease is far more informative.

§ We consider here the annotation of observed genetic variation in exons, the "Achilles heel" of WES studies. Zoonomia is a base pair annotation, and its major strength is its potential to stratify important single nucleotide variants at missense and synonymous positions (the most numerous types of variation). It has lesser informativeness for predicted loss-of-function (pLoF) variants that are a key focus for many WES studies. pLoF are defined by their occurrence in structurally-defined parts of protein-coding genes: intronic splice sites; stop-gain, missense mutation to stop or insertion of a Ter codon; frameshift (insertion-deletion of bases that are not a multiple of 3); and start- or stop-loss. Although there are nuances and complexities, pLoF variants are generally identified by their location in relation to a gene model and the genetic code (CDS exons and codon-position). As we show in *Supplementary Methods, section 8*, many of the sites at which pLoF can occur have high levels of evolutionary constraint—e.g., start codons, stop codons, intronic splice sites, and positions that can mutate to generate a stop codon). The specific base-pair constraint of a given CDS base may add little new information for an observed variant in one of these locations. In particular, evolutionary constraint adds little to interpretation of ultra-rare and often individual-specific frameshift variants as these are not directly observed in cross-species alignments (there are 3 possible changes to a single base but many possible insertions or deletions of 1, 2, 4, 5, or more bases). Moreover, impactful frameshifts can occur in non-constrained regions. For example, consider three codons: AAA-GTG-AAA (Lys-Val-Lys). A frameshift deletion of the "G" in the fourth position (REF:ALT AG:A) yields a pLoF of AAA-TGA (Lys-Ter, peptide termination). Under a random model of indel generation, this pLoF could occur in a critical, highly constrained CDS region or in an unconstrained, neutrally-evolving CDS region—base-pair constraint may add no new information. The reality may be more complex if local features make indel generation non-random.

***Section 3: Deriving Mammalian and Primate Constraint Measures for the Human Genome***

By: Michael Dong, Ola Wallerman, Xue Li, Matthew J Christmas, Voichita D. Marinescu, Jennifer R. S. Meadows

In this section, we describe the generation of the key constraint metrics. *PhyloP scores* were computed for each human position to reflect constraint over mammalian evolution (240 mammal species), and *PhastCons scores* reflect constraint over the evolution of 43 primate species.
*Notes.* Full details of the rationale behind Zoonomia, justification of the species selected, the specific species, the sources of whole genome sequences, the builds used, and the reference-free alignment approach are given in a prior paper (*11*). As these data are extensive, they are not duplicated here. The Zoonomia alignment has 241 genome sequences from 240 mammalian species (*Canis lupus familiaris* was included twice, and handled as described below). In this section, "MAF" refers to Multiple Alignment Format. Human gene models were based on *GENCODE v36 (11/2020)* (*12*), and the genome build was *hg38/GRCh38* (unless otherwise noted).

**Neutral Model.** Three human neutral models (autosomes, chrX, and chrY) were generated using the HAL-format (HAL Tools v2.1) (*13*) of the Zoonomia alignment of 241 mammalian sequences (*11*). Ancestral repeats were identified by applying RepeatMasker Open-4.0. 2013-2015 (http://www.repeatmasker.org) to the ancestral sequence of the mammal alignment. Sequence from the penultimate ancestral branch (fullTreeAnc238) was used instead of the most ancestral sequence (fullTreeAnc239), as interspersed repeats on fullTreeAnc238 had better reconstruction of the eutherian ancestral form than fullTreeAnc239 (A. Smit, personal communication, 2 Feb 2021). Repeat coordinates were converted to fullTreeAnc239 using halLiftOver (*13*). Ancestral sequence repeat sets were filtered to exclude: non-mammalian repeats shared with birds and reptiles; repeat sequences annotated as structural RNA copies, satellites, tandem repeats, and low complexity annotations; and regions where synteny was not present across the four major branches of the mammalian tree (Xenarthran, Afrotherian, Laurasiatherian and Euarchontoglires). A representative random set of ancestral repeat positions (i.e., 100Kb) from the above list was the input for the neutral evolution model calculation. An ancestor-referenced multiple alignment format alignment was extracted (hal2maf). PhyloFit (with default parameters, --subst-mod REV --EM) was applied to the corrected tree to estimate branch lengths, fixing the tree topology. The resulting alignment was processed (mafDuplicateFilter) to remove sequences that aligned multiple times to the same region in order to avoid biases due to species over-alignment. These steps was repeated for the sex-specific models but using repeat positions from chrX or chrY of the human-referenced alignment (i.e., 100Kb). Primate neutral models were constructed in the same fashion, with the ancestral branch reconstruction based on the 43 primates present in the alignment. Synteny with five of the reconstructed branches in the primates tree were checked.

**PhyloP and phastCons constraint score calculation.** The alignment was preprocessed with mafDuplicateFilter (https://github.com/dentearl/mafTools), in order to keep one sequence per species (i.e., the best match) in each alignment block compared to the sequence to the consensus for that block (*14*). For the primate-specific data, non-primate species were filtered out from the alignment using the mafSpeciesSubset command from mafTools. Alignment depth, ACGT ratios, and lists of species across the alignment were collected using the Bio.Align package from BioPython (https://biopython.org/wiki/Multiple_Alignment_Format) in a custom script. Total branch lengths for each alignment block in the multiple alignment format files were calculated using the tree_doctor command with the branch length (--branchlen) option from the PHAST software package. Two constraint scores, phyloP and phastCons, were calculated using the PHAST package, (https://github.com/CshlSiepelLab/phast). PhyloP (v1.5) was used to calculate per-base constraint and acceleration p-values (*15*). Scores were presented as –log p-values under null hypothesis of neutral evolution, where computation involved performing a likelihood ratio test at each alignment column (--method LRT) and with the output of constraint and acceleration scores (--mode CONACC). PhyloP scores calculated on the human-referenced, MAF-formatted, duplicate-filtered alignment range from -20 to 9.28. Scores from the human-referenced, primates-only alignment (43-way) range from -20 to 1.26. Higher scores reflect greater deviation from the neutral models and hence greater constraint.

Scores calculated on the primates have lower ranges, as the total branch lengths of clade-specific trees are relatively lower compared to the entire mammalian tree. PhastCons (Phast v1.5), implemented a phylogenetic hidden Markov model (phylo-HMM) to identify evolutionarily constrained elements (*16*, *17*). In contrast to phyloP single base constraint, phastCons metrics incorporate the columns of flanking bases. Model parameters were selected to match those used to generate the phastCons output for the UCSC 100 Species Vertebrate Multiz Alignment (expected-length=45, target-coverage=0.3, and rho=0.31). The phastCons, a per-base constraint score, was outputted.

**Constraint score thresholds.** We applied two metrics of constraint for 2.85 billion bases in the human genome, phyloP base scores across the evolutionary history of mammals (240 species, range -20 to 9.28, higher scores indicating greater constraint) and windowed phastCons base scores across the evolutionary history of primates (43 species, range 0 to 1, higher scores indicating greater constraint). Both metrics compared the observed phylogeny to a neutral model. The phastCons metric focuses on bases which may have diverged over mammalian history but remained under constraint over the shorter primate branch lengths.

A mammalian phyloP threshold was determined by converting absolute phyloP scores into p-values and then to q-values using a false discovery rate (FDR) correction (*18*) (R function *qvalue*). Outputs were classified as constrained or accelerated based on the sign of the score, any column with a resulting q-value ≤ 0.05 was deemed significantly constrained or accelerated (5% FDR 240 mammal phyloP constraint score ≥ 2.27, 3.53% of the human genome). We took a different approach for primate constraint given the fewer species (43) and lesser branch lengths available; a phyloP score across primates species would be underpowered to discriminate highly constrained bases from the background. We identified the phastCons threshold that yielded a similar fraction of the genome under constraint as for all mammals (i.e., phastCons base score ≥ 0.96, 3.54% of the human genome). Given that the 43 primates are a subset of 240 mammals, the locations of significantly constrained bases in mammals and primates overlap.

We note that we validated the mammalian phyloP threshold, the use of different scores to measure constraint in mammals and primates, and the base pair resolution of mammalian phyloP scores by using heritability analyses of human diseases and complex traits.

**Fraction constrained of human genome.** We estimated the lower bound for the fraction of sites under purifying selection across the genome ($\pi_{hat}$) by comparing the empirical cumulative distribution function

(ECDF) of phyloP scores across the genome to the ECDF of ancestral repeats, following the same method detailed in (*19*) where:

$\pi_{hat}$ = 1 - mins F(s)/G(s)

Here, F is the ECDF for all sites across the genome (a function of scores s) and G is the ECDF for ancestral repeats (i.e., neutrally evolving sites). We extracted coordinates of ancestral repeats from the UCSC repeatmasker track and filtered these to retain a set of ancestral repeats present in four clades of the mammalian tree (Xenarthran, Afrotherian, Laurasiatherian, and Euarchontoglires). The ECDFs of phyloP values for all positions across the genome, as well as those only in the set of ancestral repeats, were calculated using the ecdf function in R v.4.0.4. PhyloP values below -1.5 were excluded so that the extreme left tail of the ECDFs did not heavily influence estimates of constraint. We estimated $\pi_{hat}$ for human- and primate-centered phyloP sets.

***Section 4: Genomic Properties of Constraint Scores***

By: Michael Dong, Matteo Bianchi, Ola Wallerman, Chao Wang, Jennifer R. S. Meadows, Patrick Sullivan

We describe here the properties of mammalian constraint: genome-wide and in genes, gene parts, and regulatory features.

#### Annotations and basic definitions

The *inline table* below defines and references the data sources for these analyses.

| *Group* | *Type* | *Description* |
|---|---|---|
| Transcripts | GENCODE v36 (11/2020) (*12*) | All transcripts, dropping those with an annotation "issue". Selected all protein-coding (PC) transcripts, kept non-PC transcripts with different gene IDs and no PC transcript overlap. For PC genes, we required consistency in the cDNA, CDS, and peptide lengths (e.g., that the coding length was 3x the peptide length and start codon methionine). Most genes have many transcripts; we created a transcript hierarchy to pick one transcript per gene via order of selection (pickOne): MANE (*20*), gnomAD canonical transcript (*8*), BUSCO (*21*), PC, lncRNA, pseudogene, and other. We also picked a random transcript per gene (pickRandom) and lists of all transcripts (pickAll). |
| | GENCODE, cDNA | cDNAs are transcribed exons from a gene. For a protein-coding gene, cDNA consists of UTRs and CDS. Non-coding genes lack CDS. |
| | GENCODE, parts | For each PC transcript, we identified its structural parts: proximal promoter (500 and 50 bp upstream of TSS), TSS, 5'UTR exons, start codon, coding exons, canonical intronic splice sites, introns, stop codon, and 3'UTR exons. Some PC genes do not have two UTRs and some are intronless. |
| | GENCODE biotypes | PC, lncRNA, miRNA, pseudogene, and 30 others (some rare). As noted above, PC genes were given priority over any overlapping non-PC gene type. |
| Regulatory features (ENCODE3) (*21*, *22*)(*23–26*) | cCRE | cCRE: candidate cis-regulatory elements. PLS: canonical promoter-like signatures (± 200 bp of GENCODE TSS, high DHS and H3K4me3 signals). pELS: proximal enhancer-like signatures (± 2 kb of GENCODE TSS, high DHS and H3K27ac signals, low H3K4me3 signal). dELS: distal enhancer-like signatures (> 2 kb from GENCODE TSS, high DHS and H3K27ac signals). |
| | DHS | DNase hypersensitive sites in 243 cell lines and tissues |
| | ChIA-PET anchors | Anchors from chromatin loops in 24 cell types using ChIA-PET (cohesin RAD21). |
| | TF binding sites | TF (transcription factor) binding site footprints (within DHSs) |
| Regulatory features in brain | Brain OC | Open chromatin (OC) using ATAC-seq in human brain (36 datasets from fetal cortex, adult cortex, and multiple other brain regions) (*27–30*). |
| | Brain enhancers | PsychENCODE brain enhancers (*31*). |

#### Properties of constrained bases in the human genome

The *inline table* below shows the genome-wide proportions of constrained bases for three FDR thresholds. As noted in *Supplementary Methods, Section 2*, we principally consider bases as constrained in mammals for phyloP ≥ 2.27. For 43 primates, the phastCons filter analogous to phyloP FDR 0.05 in all mammals was phastCons base score ≥ 0.961. Given the higher resolution afforded by 240 mammalian species, we focus principally on phyloP in this section.

| *FDR* | *phyloP* | *Constrained bases* | *Fraction of genome †* | *Elements ‡* |
|---|---|---|---|---|
| 0.001 | ≥ 4.54 | 35,942,399 | 0.0126 | 18,557,879 |
| 0.01 | ≥ 3.27 | 57,866,048 | 0.0203 | 30,177,826 |

| | | | | |
|---|---|---|---|---|
| 0.05 | ≥ 2.27 | 100,651,377 | 0.0353 | 56,711,267 |

*† total bases with a phyloP score: 2,852,623,265 (chr1-22, X Y). ‡ After merging adjacent constrained bases.*

We compared the overlap of constraint in mammals to that in primates: mammals=100,651,377 bases, primates=101,134,907 bases, intersection=46,739,217, union=155,047,067, and Jaccard index of 0.301.

The *inline table* below shows GC content and nucleotide proportions in different genomic partitions (three phyloP FDR thresholds and genome-wide). GC content is strongly related to constraint, from 40.9% genome-wide to 55.5% in the most constrained bases. For comparison, CDS bases had GC content of 52.3% (pickOne PC transcript set).

| *Partition* | *Fraction of genome* | *GC* | *A* | *C* | *G* | *T* |
|---|---|---|---|---|---|---|
| fdr 0.001 | 0.0126 | 0.555 | 0.221 | 0.280 | 0.276 | 0.224 |
| fdr 0.01 | 0.0203 | 0.530 | 0.234 | 0.268 | 0.261 | 0.236 |
| fdr 0.05 | 0.0353 | 0.487 | 0.254 | 0.247 | 0.240 | 0.259 |
| genome | 1 | 0.409 | 0.295 | 0.204 | 0.205 | 0.296 |

As shown in *Fig. S4.1*, bases constrained in mammals showed a marked clustering tendency with the genome-wide distance between constrained bases of median 2 bp (IQR 1-8) vs a random median of 24 bp (IQR 10-47).

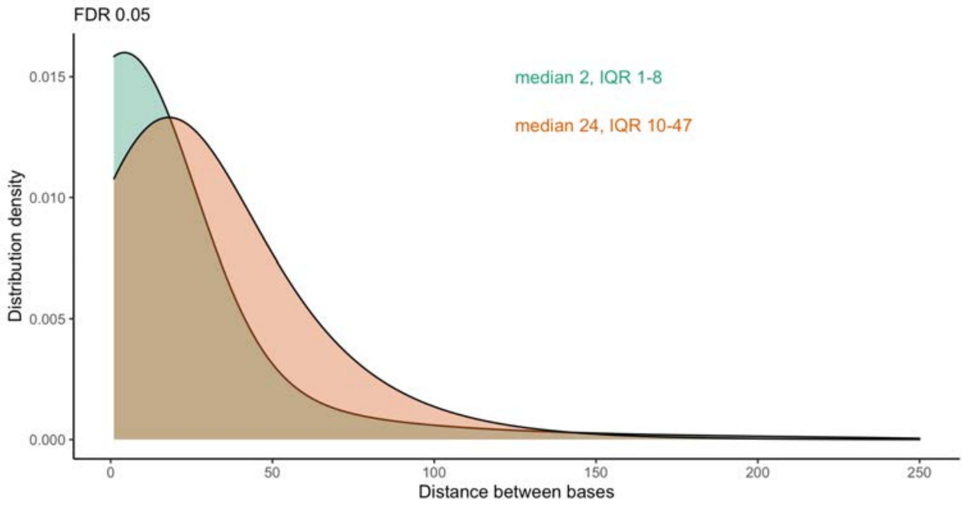

*Fig. S4.1. Clustering of constrained bases across the genome. Distance in bases between significantly constrained bases. Green is the phyloP mammalian alignment and red is a random draw with similar structure (i.e., same number of bases within the same set of genomic intervals containing phyloP measurements). FDR=false discovery rate, IQR=interquartile range.*

**Constrained genes, PC gene parts, and regulatory elements**

We next evaluated the degree of constraint within multiple important genome annotations (definitions and sources noted above). In *Fig. S4.2*, we show the relationship between the enrichment in a genome partition and the fraction of the genome occupied by that partition. The enrichment compares the fraction of the partition that is constrained in mammals divided by the fraction of the genome that is constrained. For example, introns are the largest annotation but with no notable enrichment. We note four enrichment groupings:

- Small annotations that are highly enriched for constrained bases including start codons, stop codons, exon splice sites, and TSS;
- PC transcripts are enriched for constrained bases but particularly for CDS and 5'UTRs more than 3'UTRs;
- Proximal promoters (50 bp upstream of TSS) are notably enriched for constrained bases and more than for a broader definition of proximal promoter (500 bp upstream of TSS) and for a commonly used promoter region definition (TSS ± 2kb);
- Multiple regulatory annotations showed enrichment including ENCODE promoter-like elements, brain enhancers, open chromatin in brain, and ENCODE enhancer-like elements.

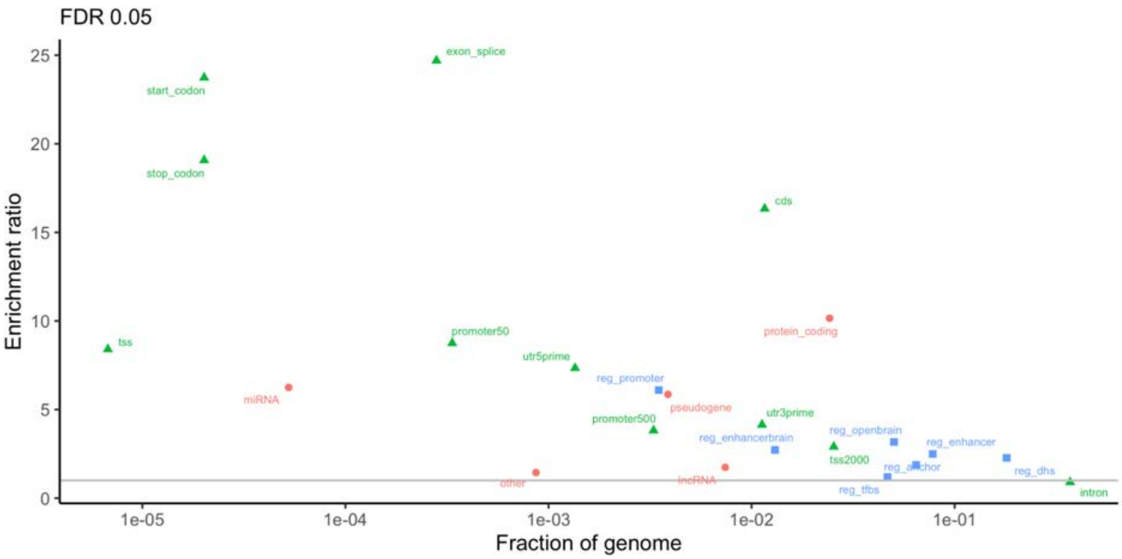

*Fig. S4.2. Enrichment ratios for genomic features. Enrichment ratios are the fraction of a feature that is constrained (5% FDR mammalian phyloP), divided by the fraction of the genome that is constrained. For example, 0.577 of CDS bases are significantly constrained (=19,069,108/33,073,903) and 0.0352 of the genome is significantly constrained (=100,651,377/2,852,623,265) for an enrichment ratio of 16.3 (=0.577/0.352). The horizontal grey line has an enrichment ratio of 1. Genomic features are defined above. Orange=cDNA exons; green=parts of PC genes (pickOne set); and blue=regulatory annotations.*

Enrichment of constrained bases in start/stop codons, splice sites, and TSS was expected as was the CDS enrichment, and these are examined more closely in other parts of this manuscript.

### Constraint in codons

As shown in *Fig. S4.2*, CDS of PC genes show high levels of constraint with 57.7% of CDS bases constrained across mammalian evolution. However, codons (not bases in isolation) are the major functional units in CDS. Thus, using pickOne PC transcripts, we computed the median phyloP score for all 192 combinations of codon and position (=64 x 3). The number of codons per amino acid is variable: 1 codon for Met and Trp; 2 codons for Asn, Asp, Cys, Gln, Glu, His, Lys, Phe, and Tyr; 3 codons for Ile and Ter; 4

codons for Ala, Gly, Pro, Thr, and Val; and 6 codons for Arg, Leu, and Ser. For 32 codons, positions 1 and 2 determine the amino acid as position 3 can be any base (Ala, Arg, Gly, Leu, Pro, Ser, Thr, and Val).

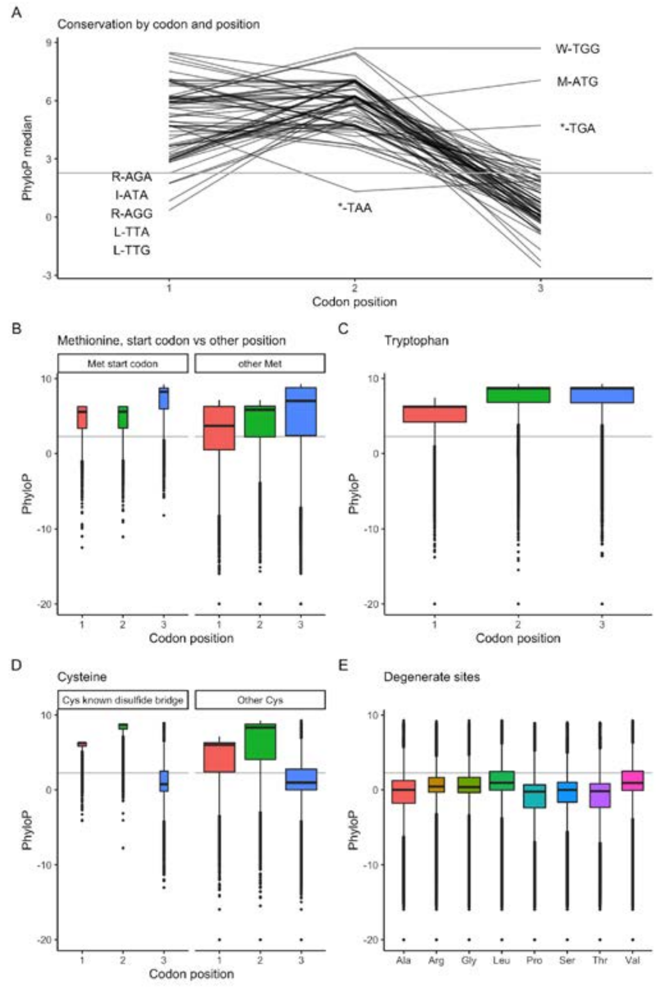

*Fig. S4.3. (A) Parallel plot of median phyloP scores for all PC CDS by codon and codon position (pickOne subset, 32,127,234 bases and 10,709,078 codons). X-axis=codon position. Y-axis=median phyloP score (grey line at 2.27 is the FDR 0.05 genome-wide cutoff). The plot has 64 lines, one per codon. The median phyloP scores for each position were 4.93, 5.98, and 0.68. However, there were exceptions to these general trends as labeled on the graph. For example, position 3 for tryptophan (W-TGG, median 8.70) and methionine (M-ATG, median 7.06) are highly constrained–these are the two amino acids with a single codon and any third position change is non-synonymous. I=Ile, L=Leu, M=Met, R=Arg, W=Trp, *=Ter. (B) Median phyloP values for Met in the start position (8.7%) versus elsewhere in the CDS (91.3%). As Met has one codon (ATG), any mutation is non-synonymous and is an start-lost if in the first codon. Met is highly constrained but particularly in the start position (median phyloP per position of 5.6, 5.6, and 8.3) compared to Met elsewhere in a protein (median phyloP 3.7, 5.8, and 7.0). The IQR is far smaller in start codon Met (IQR around 2.8) compared to Met in other positions (IQR of 4.1 or greater). (C) Tryptophan (Trp) is the other single codon amino acid. All positions are highly constrained but with variability. Of all transcripts, 93.1% contain a Trp and, for most of these transcripts, all positions in all Trp are highly constrained; for ~25% of transcripts containing ≥ 10 Trp, not all Trp positions were constrained (e.g., 96/98 Trp in MUC16 and 87/95 Trp in DMBT1 were not constrained at all positions). (D) Evaluating cysteine (Cys) residues that are or are not part of a known intra-peptide disulfide bridge (6.85% of 756,942 Cys). Non-synonymous mutations in a Cys can ablate a disulfide bridge and lead to loss-of-function due to disruption in peptide folding. Cys in known disulfide bridges had higher median phyloP values and narrower IQR. (E) Half of all codons are degenerate as positions 1 and 2 determine the amino acid and position 3 can be any base. The box plots show the distribution of phyloP scores in the degenerate third positions of Ala, Arg, Gly, Leu, Pro, Ser, Thr, and Val. The medians show low levels of constraint (Pro -0.22 to Val 0.93). However, there is variability with 18.6% of 5,263,653 degenerate sites being highly constrained.*

*Fig. S4.3* shows an analysis of constraint in 32.13 million CDS bases. *Fig. S4.3A* shows the median phyloP scores per codon and position. Over all CDS codons, position 1 was highly constrained (phyloP median 4.93), position 2 yet more constrained (phyloP median 5.98), and position 3 far less constrained (phyloP median 0.68). Several codons differed from this general trend. Position 3 in the single codon amino acids Met and Try are highly constrained as any mutation is missense. For the stop codons (TAA, TAG, and TGA), position 2 of TAA is less constrained as 1 of the 3 possible mutations is synonymous. TGA position 3 is highly constrained as all mutations are missense (stop-lost) whereas, for TAA and TAG, 1 of 3 mutations are synonymous. As shown in *Fig. S4.3B-E*, key aspects of this CDS constraint overview are that phyloP scores were sensitive to multiple features of the protein code.

### Constraint and human genetic variation

We next evaluated the relationship between mammalian constraint and allele frequency. At the time of this writing in 10/2021, the most extensive catalog of human variation is TOPMed (*6*). TOPMed (v8) is the largest and most comprehensive dataset currently available, and is based on WGS of 140,306 extant humans from diverse ancestries. The numbers of base positions for the key variant types are: 552,420,373 biallelic single nucleotide variants (84.5%, SNV), 27,946,337 biallelic deletions (4.3%, biDEL), 9,402,236 biallelic insertions (1.4%, biINS), and 64,184,482 multiallelic variants (9.8%, 3-15 alleles, multiple combinations of SNVs and indels). A SNV is defined by its position in the reference sequence, the reference base, the observed alternative allele, and the allele count (AC) and allele frequency (AF) in a sample. As Zoonomia is a mammalian species alignment, we use TOPMed AC and AF over all samples (to consider humans as a species). TOPMed variants were annotated by adding phyloP scores and the mutational consequences for CDS SNVs.

*Fig. S4.4* shows the genome-wide relationship between constraint and variation in TOPMed. There is a small negative correlation between constraint and AC for SNVs (*Fig. S4.4A*). As biallelic AC=1 is the most common SNV class (*Supplemental Methods Section 1*), most SNVs occur at non-constrained sites. The small negative correlation of constraint with AC suggests an impact of purifying selection in a subset of SNVs. For the other variant types, there is minimal relation of constraint with frequency; because Zoonomia is a base pair annotation, its salience is highest for SNVs (discussed in *Supplemental Methods Section 2*).

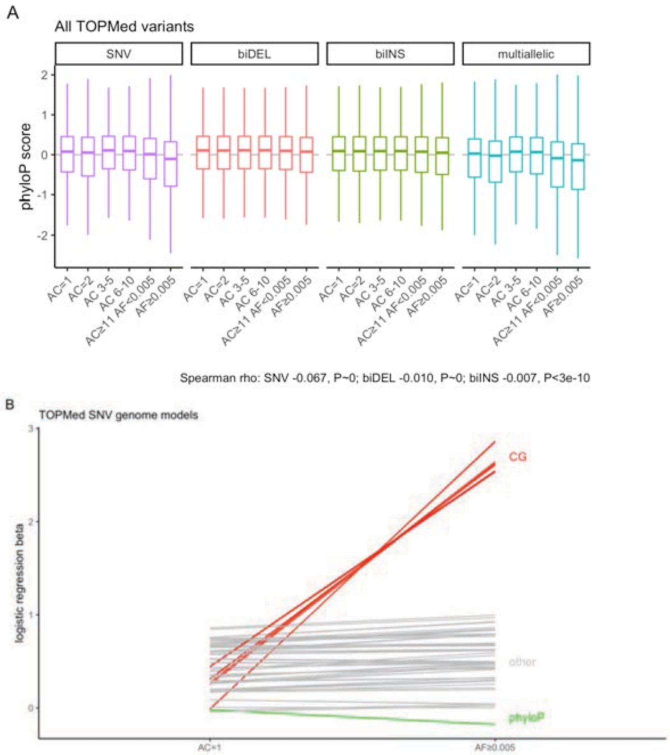

*Fig. S4.4. Plots of the relationship between AC and phyloP score for all TOPMed variants by frequency and type. Variants were classified in 5 AC bins of decreasing rarity plus common variation (AF ≥ 0.005). We included all ancestries as appropriate for mammalian comparisons. (A) Description of genome-wide constraint, variant type, and variant frequency. The boxplot thick line is the median, the rectangle shows 25th–75th percentiles, and whiskers extend 1.5 times the IQR above and below the box. Violin plots were impractical due to the number of variants. AC=allele count, AF=allele frequency. There is a small negative correlation between AC-phyloP for SNVs and near zero correlations for non-SNV types of variation. (B) Parallel plot of modeling results for*

*singleton (AC=1) and common SNVs (AF≥0.005). Because nearly all P-values were smaller than the operating system's minimum numeric value (P < 2.2e-308), the Y-axis shows patterns of beta coefficients between the two logistic regression models. Each model had 64 terms (phyloP score plus 63 base sequences AAC to TTT compared to reference AAA). Green line shows betas for phyloP constraint. Most of the 3 base context terms were similar (grey lines) with the exception of the 8 terms containing the dinucleotide CG (red, GCN or NGC). The modelling was done in this way (i.e., singleton SNVs vs 10x random local positions and common SVS vs 10x random local positions) due to the great imbalance in the numbers of variants: of all 552 million SNVs, 45.6% AC=1 and 2.6% AF ≥ 0.005.*

To evaluate the AC-SNV relationship more formally, we contrasted the extreme cases from the leftmost facet of *Fig. S4.4A* (i.e., singletons with AC=1) to the common variants with AF ≥ 0.005). In the first model, the dependent variable was the genome-wide occurrence of a singleton biallelic SNV (AC=1) compared to 10 randomly selected positions for each SNV ± 10kb. Model predictors were phyloP score and the three base context centered on the SNV. The second model was the same as the first but the dependent variable was the presence of a common biallelic SNV (AF ≥ 0.005). Because nearly all terms had P~0, we focused on the patterns of logistic regression beta values (*Fig. S4.4B*). Controlling for base context and in comparison to random nearby positions, the beta for phyloP constraint was more negative for common SNVs (beta -0.178, se 2.38e-4) than singleton SNVs (beta -0.020, se 5.36e-5), consistent with the Spearman correlation. The beta terms for most of the three base context terms in the models were similar; the notable exceptions were eight base context terms containing the dinucleotide CG where the betas were considerably larger compared to random nearby positions.

We next focused on the constraint of SNVs occurring in CDS. *Fig. S4.5* summarizes the 6,140,669 TOPMed SNVs that occur in 31,809,819 unique CDS bases. Most SNVs were rare (AC=1 44.8%) and common SNVs were the least frequent grouping (AF≥0.005 1.4%). Most were missense mutations (missense 65.4%, synonymous 32.8%, and mutation to stop 1.8%). SNV mutation types show different relations with AC and constraint. Synonymous SNVs occur at non-constrained sites with a small inverse relation with AC. Missense SNVs and rare terminal mutation SNVs occur at highly constrained sites and the median constraint levels are very high except for common SNV.

B

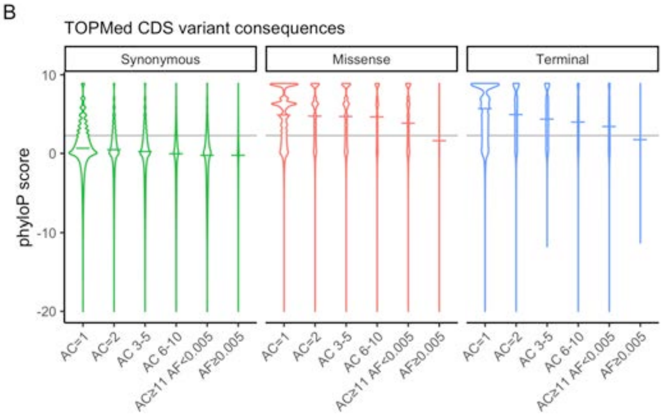

| Consequence | AC group | SNVs | median | p25 | p75 |
|---|---|---|---|---|---|
| Synonymous | AC=1 | 833,832 | 0.68 | -0.37 | 2.39 |
| | AC=2 | 306,718 | 0.51 | -0.66 | 2.26 |
| | AC 3-5 | 341,124 | 0.26 | -1.40 | 1.93 |
| | AC 6-10 | 183,752 | -0.03 | -2.67 | 1.52 |
| | AC≥11 AF<0.005 | 302,509 | -0.20 | -3.28 | 1.29 |
| | AF≥0.005 | 46,009 | -0.21 | -3.22 | 1.18 |
| Missense | AC=1 | 1,859,513 | 4.91 | 1.72 | 6.98 |
| | AC=2 | 640,927 | 4.79 | 1.58 | 6.99 |
| | AC 3-5 | 674,911 | 4.74 | 1.49 | 7.01 |
| | AC 6-10 | 335,357 | 4.68 | 1.31 | 7.00 |
| | AC≥11 AF<0.005 | 466,002 | 3.88 | 0.91 | 6.57 |
| | AF≥0.005 | 40,706 | 1.63 | 0.05 | 4.60 |
| Terminal | AC=1 | 59,398 | 5.73 | 2.32 | 8.66 |
| | AC=2 | 18,025 | 4.94 | 2.08 | 8.32 |
| | AC 3-5 | 16,741 | 4.38 | 1.87 | 7.03 |
| | AC 6-10 | 7,301 | 4.03 | 1.57 | 6.87 |
| | AC≥11 AF<0.005 | 7,560 | 3.42 | 1.09 | 6.28 |
| | AF≥0.005 | 284 | 1.78 | 0.26 | 4.10 |

*Fig. S4.5. (A) Relation of AC and constraint for all TOPMed CDS variants by the consequences of the variant (synonymous, missense, or terminal/mutation to stop). These violin plots show phyloP score densities; within each facet, the area within the violin is proportional to the number of occurrences. The horizontal colored lines are medians. The grey horizontal line is at phyloP=2.27. Most SNVs were rare (AC=1 44.8%, AC=2 15.7%, AC 3-5 16.8%, AC 6-10 8.6%, AC≥11 to AF<0.005 12.6%, and AF≥0.005 1.4%). Most were missense mutations (missense 65.4%, synonymous 32.8%, and mutation to stop 1.8%). (B) Summary data used to create Fig. S4.5A, noting the differences between AC=1 (median phyloP 0.68, 4.91, and 5.73) versus common SNVs (-0.21, 1.63, and 1.78).*

Due to the importance of CDS, we fit two multivariable logistic regression models. The data frame is fully defined in the next section. For the first model, the dependent variable was the presence of a singleton biallelic SNV with AC=1 versus all other CDS sites containing no variation. For the second model, the dependent variable was the presence of a common biallelic SNV (AF ≥ 0.005) versus all other CDS sites containing no variation. The predictors were phyloP score, CDS base (A, C, G, or T), codon position (1, 2, or 3), the number of possible synonymous mutations at that site (0-3), if the position could mutate to stop, the number of CDS transcripts in which a base is used, whether the base was in multiple reading frames, and local constraint (mean phyloP score in a rolling window of ±9 bases excluding the current base).

The results are summarized in the *inline table* below. For biallelic singleton SNVs (AC=1), the major effects were: greater odds of occurring at a C or G base, in codon positions 1 and 2, lesser odds of occurring at a base that could mutate to stop, inverse association with constraint, and greater odds of occurring in regions of higher constraint. The catalog of singleton SNVs is incomplete and this class of variant is of recent origin (*de novo* in the subject or an immediate ancestor) with lesser exposure to purifying selection. In comparison, common biallelic SNVs have a stronger inverse association with constraint, greater odds of occurring at a C or G base, and tend not to occur in important positions (codon position 1 or 2, or at bases that could mutate to stop). The major difference in the two analyses is in the effect of local constraint where singleton SNVs are more likely to occur in constrained regions and common SNVs are less likely.

| | *AC=1 vs other* | | | *AF≥0.005 vs other* | | |
|---|---|---|---|---|---|---|
| *Variable* | *Beta (SE)* | *Statistic* | *P* | *Beta (SE)* | *Statistic* | *P* |
| phyloP score | -0.043 (0.00025) | -171.63 | < 2.2e-308 | -0.20 (0.00085) | -238.83 | < 2.2e-308 |
| CDS base C | 0.55 (0.0018) | 301.85 | < 2.2e-308 | 1.00 (0.01) | 100.31 | < 2.2e-308 |
| CDS base G | 0.49 (0.0019) | 259.86 | < 2.2e-308 | 0.93 (0.01) | 91.23 | < 2.2e-308 |
| CDS base T | -0.19 (0.0021) | -92.13 | < 2.2e-308 | -0.11 (0.012) | -9.63 | 5.97e-22 |
| Codon pos 1 | 0.12 (0.0025) | 48.32 | < 2.2e-308 | -0.038 (0.012) | -3.07 | 0.00214 |
| Codon pos 2 | 0.10 (0.0026) | 39.03 | < 2.2e-308 | -0.015 (0.013) | -1.09 | 0.277 |
| Mut to self | 0.0047 (0.001) | 4.46 | 8.23e-6 | 0.01 (0.0048) | 2.11 | 0.0347 |
| Mut to stop | -0.20 (0.0022) | -87.98 | < 2.2e-308 | -0.23 (0.014) | -16.36 | 3.94e-60 |
| Base usage | -0.014 (0.00046) | -29.61 | 1.01e-192 | -0.016 (0.0025) | -6.3 | 3.06e-10 |
| Base in mult frames | 0.068 (0.0065) | 10.46 | 1.34e-25 | -0.077 (0.038) | -2.06 | 0.0398 |
| Local cons (±9 bp) | 0.0089 (0.00044) | 20.53 | 1.15e-93 | -0.027 (0.002) | -13.7 | 1.00e-42 |

**Constraint and common SNVs**

Because common biallelic SNVs are the main markers used in GWAS, we conducted additional analyses focusing on this class of variants. These SNPs had AF ≥ 0.005 over all TOPMed samples. As TOPMed is composed of ancestral groups of different sizes, it is possible that a common variant in a smaller ancestry group might not be common in the full group so we also included SNPs in AFR, ASI, EUR, or SAS that were common in any one population. There are 15,777,878 SNPs (i.e., common biallelic SNVs) in TOPMed and 1.849% (N=291,669) are constrained, far less than the fraction of the fraction of the genome that is constrained (3.53%). For each common SNP, we added annotations for location in CDS, ENCODE3 regulatory elements (promoter, DHS/open chromatin, enhancer, or transcription factor binding site), or in the "fine-mapped" set of distinct eQTL-SNPs in any GTEx tissue (posterior probabilities ≥ 0.8 in 3 credible set algorithms). We fit a logistic regression model over these data with common SNP constraint as the dependent variable and six SNP annotation predictors.

The *inline table* shows the results. All terms were significant. SNP location in CDS, promoter, GTEx fine-mapped eQTL-SNP, open chromatin, and enhancer regions were positively associated and location in TF binding site was negatively associated with whether a common SNP was under evolutionary constraint.

| *term* | *fraction †* | *estimate* | *Odds ratio* | *se* | *statistic* | *p* |
|---|---|---|---|---|---|---|
| cds | 0.007 | 3.09 | 21.977 | 7.30E-03 | 423.15 | < 2.2e-308 |
| reg_promoter | 0.002 | 1.15 | 3.158 | 2.09E-02 | 55.18 | < 2.2e-308 |
| finemap eQTL SNV | 0.0003 | 0.89 | 2.435 | 5.51E-02 | 16.23 | 3.12E-59 |
| reg_dhs | 0.103 | 0.89 | 2.435 | 5.50E-03 | 160.99 | < 2.2e-308 |
| reg_enhancer | 0.043 | 0.41 | 1.507 | 7.43E-03 | 55.05 | < 2.2e-308 |
| reg_tfbs | 0.0025 | -0.12 | 0.887 | 1.16E-02 | -10.53 | 6.56E-26 |

*† the number of SNPs in each annotation category divided by the total number of common SNPs. The "fine-mapped" eSNPs are a tiny subset (0.14%) of the full set of eSNPs. We also ran a model with all GTEx eQTL-SNPs and got rather different results: fraction=0.271, estimate=-9.143, se=0.00470, statistic=-30.4, and p=1.14e-202.*

In the above analysis, common CDS SNPs had 22-fold ($e^{3.09}$) greater odds of being constrained: 28% of common SNPs in CDS were constrained compared to only 1.7% of common SNPs not in CDS. This was unexpected given the results in *Fig. S4.5,* but could be explained if selection effects were variable across genes—common CDS SNPs may be permitted in some genes but not others. An important role for some genes is to generate variability in protein structure (e.g., olfactory receptors) whereas many genes are highly intolerant of any CDS variation.

We determined the number of common, constrained CDS SNPs for each PC gene. Irrespective of amino acid consequence, we found that 37.8% of genes contained no common SNPs in constrained CDS whereas other genes had up to 10% of CDS bases composed of common constrained SNPs. (We found similar findings for any common SNP regardless of constraint, 10.7% of genes had none and some genes had common SNPs in up to 26% of CDS).
Gene set analysis (hypergeometric test, background of all PC genes, Bonferroni correction to P < 0.05) of 980 PC genes in the top 5% for fraction of CDS containing common, constrained SNPs were significantly enriched for:

- G-protein coupled receptors (GPCR)
- "Druggable" genes (*32*)
- Olfactory receptors, taste
- Skin development
- Cell defense response to bacterium, cell killing, lymphocyte chemotaxis, CCR chemokine receptor binding

These genes contain human variable alleles in many highly constrained positions. The biological processes suggested by these gene sets function at the interface of a mammal and its environment in the sense of allowing adaptation of a mammal to its environmental niche: olfaction and taste are central to identifying nutrients and avoiding predators and phytotoxins; skin development for thermoregulation, protection, and coloration; and immune processes central to adaptation to local microbiota.

GPCRs are a large and diverse gene family (~4% of human genes) of cell surface receptors that detect extracellular molecules. Some 25% of approved drugs target GCPRs (*33*). GPCRs are involved in multiple sensory modalities (vision, taste, smell), key neurotransmitter systems relating to behavior and mood (e.g., dopamine and serotonin), and immune sensing (chemokine receptors) among others. Many of the gene sets are due to different types of GCPR (e.g., there are 116 olfactory receptors along with chemokine receptors). There are also 178 genes in this list that are "druggable" (*32*) but are not GPCRs. Other notable groups of genes include: hepatic metabolic genes—alcohol and aldehyde dehydrogenases (including *ADH1B*), cytochrome P450 genes (including the clinically important *CYP2D6* and *CYP2A6*)—and genes involved in blood groups (*ABO*) and hemoglobin subunits (*HBE1*, *HBG2*, and *HBQ1*); and steroid receptors (including *ESR1*). Notably the list does not include key CNS neurotransmitter receptors like *DRD2*.

Moreover, this is not a dismissable set of genes. Many of these genes have disease or medical relevance as 131 have an OMIM entry for diverse conditions: albinism, alcohol dependence, amyloidosis, asthma, familial candidiasis, warfarin resistance, deafness, glycogen storage diseases, hypogonadotropic hypogonadism, hypotrichosis, and retinitis pigmentosa. The list also includes neurological disorders (developmental delay, epilepsy, and spinocerebellar ataxias).

#### *Section 5: Benchmarking ClinVar and CADD against phyloP constraint*

By: Matteo Bianchi, Graham Hughes, Patrick Sullivan and Jennifer R. S. Meadows

**ClinVar class annotated for mammalian phyloP score.** The ClinVar (*34*) summary file was downloaded (7 May 2021), and SNVs extracted and annotated with mammalian phyloP scores. Summary statistics were calculated for benign, pathogenic, and "likely classes" for these groups (*inline table below*). ClinVar pathogenic variants had markedly higher phyloP scores (Wilcoxon Rank-Sum $P < 2.2 \times 10^{-16}$).

| *Group* | *Subgroup* | *N* | *phyloP mean ± sd* |
|---|---|---|---|
| Benign | benign | 87,881 | 0.09 ± 3.71 |
| | likely benign | 143,761 | 0.07± 3.88 |
| | total | 231,642 | 0.08 ± 3.82 |
| Pathogenic | pathogenic | 46,248 | 6.05 ± 3.02 |
| | likely pathogenic | 27,637 | 6.48 ± 2.70 |
| | total | 73,885 | 6.21 ± 2.91 |

In addition, for variants in an early version of ClinVar (01/2016), we compared the phyloP scores for the classification of the same variants in 2021. For variants whose status was unchanged (benign/likely benign or pathogenic/likely pathogenic), phyloP scores were highly discriminating (median 0.27 vs 6.36). Similarly, for variants whose status changed from uncertain to benign or to pathogenic, phyloP scores were also highly discriminating (median 0.76 vs 6.34). Variants whose status was downgraded had a median under that of all CDS bases (3.37 vs 3.6) (*inline table below*). This analysis suggests the prospective utility of phyloP scores in clinical variant classification.

| *Status* | *ClinVar 2016* | *ClinVar 2021* | *N* | *p50* | *p25* | *p75* |
|---|---|---|---|---|---|---|
| Unchanged | benign | benign | 16,099 | 0.27 | -0.97 | 2.13 |
| Unchanged | pathogenic | pathogenic | 19,706 | 6.36 | 5.19 | 8.75 |
| Classified | uncertain | benign | 3,263 | 0.76 | -0.46 | 3.75 |
| Classified | uncertain | pathogenic | 1,473 | 6.34 | 4.74 | 8.72 |
| Downgraded | pathogenic | benign | 327 | 3.37 | 0.39 | 6.08 |

**CADD compared with mammalian phyloP.** Combined Annotation Dependent Depletion (CADD) (*35*) integrates multiple annotations, including functional and pathogenicity predictions, experimentally measured regulatory effects and modifications, information on genetic context, as well as constraint metrics based on existing mammalian and vertebrate scores (i.e. GERP++, phastCons, phyloP). While using a single universal cutoff for CADD to discretise variants into pathogenic and benign is inherently flawed (due to the nature of the score and the unique nature of any analysis framework), the recommendation is to further evaluate variants with a scaled CADD score ranging from 10 to 20 or above (*35*). A scaled score ≥10 indicates that the variants are predicted to be the 10% most deleterious substitutions identifiable in the human genome, while a score ≥20 indicates the 1% most deleterious.

We downloaded gnomAD release 3.0 (*8*) SNV positions with CADD v1.6 (*36*) directly from the CADD browser (https://cadd.gs.washington.edu/download) and annotated these with mammalian phyloP scores. This resulted in 540,568,449 variant positions for consideration, and this set was used to generate all classes of variants described below. For multiallelic variants, we calculated the average CADD score of all alternate alleles and found that phyloP and CADD metrics were moderately correlated (Spearman rho = 0.49, $P < 2.2\times10^{-16}$).

To explore the predictive potential of mammalian phyloP scores in comparison to scaled CADD scores, we evaluated the extent to which these metrics were able to capture genetic variation overlapping with different gene and regulatory annotations with increasing analysis thresholds. The gene and regulatory annotations were obtained from i) GENCODE (v36) using one gene transcript (pickOne, see Table page 12), ii) ENCODE3 (DHS, enhancer, promoter, TFBS) and iii) UNICORNs (*Companion paper #1, FS1* and *Supplementary Methods, Section 8*). The phyloP threshold was kept constant at ≥ 2.27 and CADD ranged from ≥10 to ≥20 in single score steps (*Fig. S5.1A*). A convergence between the fraction constrained by both phyloP and CADD was observed at several annotations at approximately CADD ≥15, suggesting that phyloP scores could potentially be employed for prioritizing variants (*Fig. S5.1B*). At this general cutoff certain gene and regulatory features (e.g. DHS, enhancer, TFBS) showed a greater enrichment for phyloP, indicating that this metric might perform better for these classes of annotations (*Fig. S5.1B*). Considering that one of the current objectives of the CADD project is to improve and advance prediction in regions of the genome that are not protein-coding (e.g. cis-regulatory elements and non-coding RNA species) (*36*), the implementation of Zoonomia mammalian phyloP within the CADD framework may greatly boost the power to identify functional relevant genetic variation in these regions.

To assess the independent and shared value of phyloP and CADD scores as contributors to the same above-described gene and regulatory elements, we again calculated the enrichment as above, but for three different categories of variant positions, those annotated with i) only scaled CADD, ii) only phyloP, and iii) with both scaled CADD and phyloP scores (*Fig. S5.1C*). Of note, the category containing variant positions with both scaled CADD (≥10 step and above) and phyloP constraint scores (≥ 2.27) showed the highest enrichment for the great majority of the annotations, especially at average levels of deleteriousness (i.e. scaled CADD ≥ 15). This indicates the utility of mammalian phyloP scores in the context of functional prediction. For higher steps of deleteriousness, it seems that the great majority of the annotations considered are enriched for variants with only constrained phyloP (i.e. ≥ 2.27), suggesting that this metric can capture the residual functional signals (*Fig. S5.1D*).

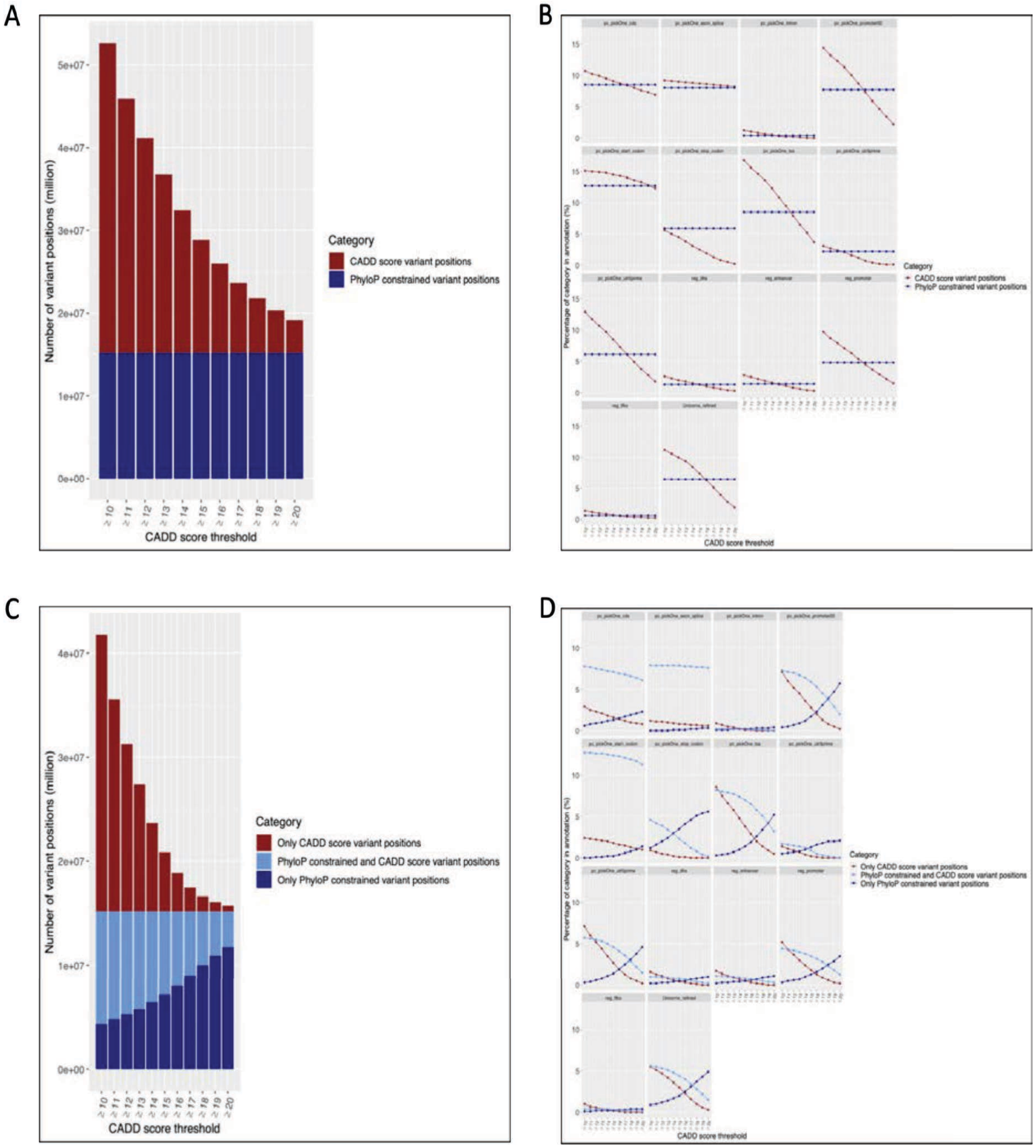

*Fig. S5.1. (A) Total number of variable positions contrasted at each scaled step of CADD. The phyloP (score ≥ 2.27) and number of analysed positions is invariant in each step (N=15,165,814). (B) Enrichment of scores in each genomic class partitioned across CADD scores. (C) Number of variable positions analysed at each scaled step of CADD. Each category is variable as the intersection of CADD and phyloP is also considered. (D) Enrichment of phyloP only, and intersected CADD and phyloP scores are contrasted against the scaled CADD score only category for each genomic class.*

**Case example of variation at constrained base pair positions implicated in human health.** A recent WES study using varination from 645,626 individuals, associated body mass index (BMI) with rare coding variants in *GPR75* (*37*). We annotated the 1,620 coding bases of *GPR75*, using the genomic definitions and coordinates reported by the authors, and showed that the fraction of constrained bases was even across this space (*Fig. S5.2A*). This proportion was increased when only the 46 pLOF sites driving the BMI results were considered.

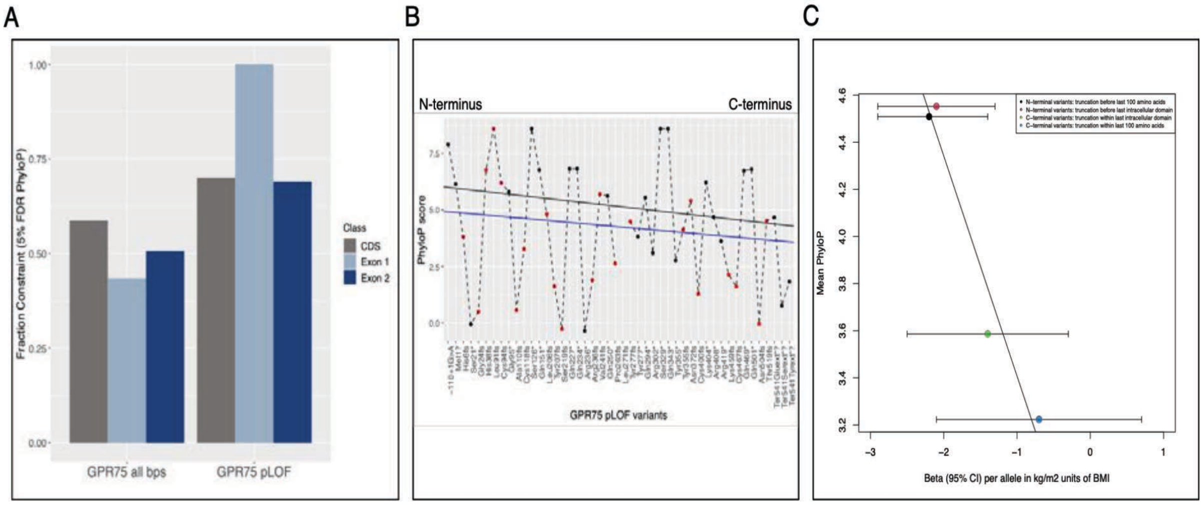

*Fig. S5.2. GPR75 annotated with mammalian phyloP. (A) Fraction of constraint in GPR75 scored for all defined CDS bases, and for bases identified as leading to pLoF. Note, GPR75 pLoF for exon 1 contains a single constrained variant. (B) PhyloP scores for each pLoF variant. SNV (N=24) and non-SNV (N=21) variants are indicated (black and red dots respectively). Regression lines were generated for all variants (blue) or only SNVs (black). (C) All variants used to illustrate the relationship observed between phenotypic effect size and mean phyloP score for the considered gene fragments (regression coefficient = -0.93, P = 0.02).*

For all 46 pLoF variants we extracted the corresponding phyloP score. Polymorphic features were considered as follows: for SNVs ($N$=24), we assigned the phyloP score of the corresponding base, for deletions ($N$=16) and duplications ($N$=5) we averaged the phyloP score of the deleted or duplicated base(s), for insertions ($N$=1) we set the phyloP score as missing. The last three variant types were classed as non-SNV. Nevertheless, we acknowledge that, given that a phyloP score is a single nucleotide-specific metric, extrapolating a composite score for non-SNV variants may be difficult to directly interpret. We explored the relationship between individual phyloP score and effect on BMI. Although not significant, a clear negative trend towards lower constraint scores at the C-terminus was noted *Fig. S5.2B*. This was irrespective of which variant set was used (All variants: regression coefficient = -0.03, $P$ = 0.32; only SNVs: regression coefficient = -0.04, $P$ = 0.36). The authors noted that rare genetic variation at the N-terminus was more deleterious for the functionality of the protein, causing truncations with greater effect and the individual phyloP burden score reflected this. The authors further quantified the beta effect truncating variants had on BMI using four regions, N- and C-terminal truncation spaces, where the outcome was within a 100 amino acid space, or where it occurred within an intracellular domain. We investigated the relationship between the mean phyloP score of protein-truncating variants within the four spaces and beta effect, and found a very strong correlation (regression coefficient = -0.93, $P$ = 0.02), although beta measures had overlapping confidence intervals (*Fig. S5.2C*). The mean phyloP score of each of these four spaces exceeds the phyloP base pair threshold for constraint ($\geq$ 2.27), however we note that the stronger beta effect is borne by higher phyloP scores. This indicates that there are likely functional nuances among pLOF variants that phyloP highlights in the context of this phenotype and this gene.

**Quantification of variation at constrained amino acids implicated in human health.** To investigate the frequency of constrained amino acid sites involved in known human disease states, all sites showing constrained residues across protein alignments were extracted from the set generated using the Tool to infer

Orthologs from Genome Alignments (*Companion paper #1, FS1; Companion paper #10, Kirilenko et al*). The TOGA alignment dataset consisted of 37,552 alignments representing 19,300 genes, with different alignments existing for genes with multiple human transcripts. Sites where a species had a deletion (represented by a gap in the alignment) or an indeterminate amino acid (represented by an 'X'), were ignored and did not impact the 'constrained' status of a specific site. TOGA alignments were selected as they contained both Zoonomia and previously sequenced taxa, including non-placental mammals (See *Companion paper #1, FS1*). Constrained sites across alignments were identified using a Perl script. ClinVar annotations were used to label sites, and entries available for each gene were collated, with mutations considered 'pathogenic' and 'likely pathogenic' extracted. As a comparison, entries considered 'benign' and 'likely benign' were also collated to determine whether constrained sites had more pathogenic than benign instances. As the human reference sequence used in ClinVar was not necessarily the same reference sequence present in the protein alignment, the relative positions between both were determined by inferring a pairwise alignment using MAFFT (*38*). Gaps in either sequence were accounted for when present, such that the relative positions between human sequences could be identified and explored in ClinVar. If the residue at a site in the TOGA human sequence differed to the residue at the same relative site in the ClinVar reference sequence, that site was not investigated further. Relative sites between human sequences were determined using several Perl scripts (https://github.com/GMHughes/ZoonomiaScripts). For each gene, the transcripts available and the list of constrained sites, with ClinVar references, are reported in *table S19*.

The TOGA dataset was used to assess if constraint scores could be used to predict deleterious mutations at constrained sites using the case example of ADRB2. Previously, a deep mutational screen of the sole human exon had been performed, and primary signalling of the protein assessed (*39*). The top 15 sites in ADRB2 reported by the authors (*39*) as intolerant to mutation in (P211, G315, Y326, W286, N51, F289, D113, S319, W158, G83, C190, C184, W99, C191, D79) were constrained across all mammals in our data, as were a number of other sites predicted as being crucial to functionality (G102, C106, A181, Y185, S203, V222, A271; 161/483 sites constrained). Taken together, these results suggest that the single metric of phyloP could help predict and direct the genetic basis of novel disease-causing substitution mutations.

**Naturally occurring adaptive 'pathogenic' variants in non-human mammals to aid the study of human health.** While certain substitutions may result in a human disease state, identification of these residues occurring naturally in non-human mammals has the potential to act as biomarkers or sources of novel therapies. Sites showing pathogenic amino acid substitutions in ClinVar were used to identify instances of pathogenic residues occurring naturally as a wild type in non-human mammals. A total of 697 such mutations were found across 330 genes (TOGA alignment from above). One example of a human disease mutation naturally occurring in other mammals was observed in SOD1, the gene responsible for amyotrophic lateral sclerosis (ALS) in humans. Pathogenic mutations were found in several mammal species at five sites: E22, E23, D97, E101 and N140. The E22K mutation, which results in the loss of a *TaqI* restriction site (*40*), naturally occurs in three xenarthran species (*Tamandua tetradactyla*, *Myrmecophaga tridactyla*, *Choloepus hoffmanni*), one bat species (*Carollia perspicillata*) and in all five rhinocerotid species present (*Rhinoceros unicornis*, *Dicerorhinus sumatrensis sumatrensis*, *Diceros bicornis*, *Ceratotherium simum*, *Ceratotherium simum cottoni*), possibly representing the ancestral rhinocerotid state. Another mutation, E101K was observed in equids (*Equus przewalskii*, *Equus caballus*, *Equus burchellii boehmi*, *Equus asinus*) and three rhinocerotid species. These data augment a previous analysis of SOD1where an E40K mutation otherwise pathogenic in humans was observed as naturally occurring in *E. caballus* (*41*). Our data show that this mutation is present in all perissodactylid species for which we have sequences, suggesting not only that multiple mutations in SOD1 that are pathogenic to humans were present in the Perissodactyla ancestor, but that these residues may offer some selective advantage. Additional mutations for the other three sites were found across a range of mammals (E23L, D97V, N140K; *table S20*). Naturally occurring adaptive 'pathogenic' variants expressed without a disease phenotype in non-human mammals warrant further investigation, representing a source of information that may lead to the development of new therapies, alleviating disease symptoms. Additionally, given that several species analysed are 'critically endangered', this work also highlights the important role that wildlife conservation will play in the future of genomics-based medicine.

***Section 6: Constraint Variation and GWAS of Human Disease***
By: Steven Gazal

We applied stratified LD score regression (S-LDSC), a polygenic approach for partitioning the heritability ($h^2$) causally explained by common human SNPs across genomic annotations (*42–44*). All analyses were conditioned to the baseline-LD model (version 2.2, 97 annotations including functional annotations and multiple constrained annotations) and to six ENCODE3 annotations (PLS, pELS and dELS annotations and corresponding flanking regions defined in *Supplementary Methods, Section 2*) for ~10M reference SNPs with a minor allele count ≥ 5 in the 1000 Genomes Project European reference panel (*45*). All analyses were performed on hg19 (after liftOver of constraint scores from hg38).

We analyzed a set of 63 GWAS summary statistics with independent association data (labeled as independent traits) by excluding genetically correlated traits in overlapping samples by measuring the intercept of cross-trait LD score regression (*46*), as previously described (*42*). For traits with summary statistics from the UK Biobank, we also excluded traits with a squared genetic correlation (*47*) greater than 0.1 (similar to the squared phenotypic correlation threshold used in (*48*)). The 63 datasets included six traits that were duplicated in two different datasets (genetic correlation of at least 0.9). After excluding these duplicates, we analyzed 57 independent diseases and complex traits. Traits were prioritized using the z-score for nonzero SNP-heritability computed using S-LDSC with the baseline-LD model (minimum of 6, as in (*43*)). We also performed a meta-analysis of 11 autoimmune diseases and blood cell traits (5 included in the 63 datasets and 6 additional traits, increasing the total number of GWAS datasets analyzed to 69; *table S1*) and 9 brain disorders (all included in the 63 datasets); note that Alzheimer disease was considered as a brain disorder only. Results were meta-analyzed across traits using random effects.

To evaluate phyloP and phastCons constraint scores, we created 15 annotations based on genome-wide percentile scores for each score: phyloP in mammals, phyloP in primates, phastCons in mammals, and phastCons in primates. We validated the use of different scores to measure constraint in mammals and primates by the fact that phyloP scores were unable to detect single-base constraint in primates due to lack of power (43 primates, lesser branch lengths) to detect significant $h^2$ enrichment. In mammals, both phyloP and phastCons scores performed very similarly in heritability analyses, and we defined phyloP as superior for having single-base resolution (*fig. S1*). In addition, by performing S-LDSC conditional analyses of different constrained scores, we observed that a constraint annotation derived from phastCons scores in mammals was systematically less informative than annotations derived from phyloP scores in mammals and phastCons scores in primates (see *inline table*).

| | | | Model 1 | | | Model 2 | | | Main Model | | |
|---|---|---|---|---|---|---|---|---|---|---|---|
| | | **Prop._SNPs** | **Enrichment** | **Tau*** | **Tau* P** | **Enrichment** | **Tau*** | **Tau* P** | **Enrichment** | **Tau*** | **Tau* P** |
| **Mammals phyloP** | - | **1.52%** | 7.70 (0.35) | 0.20 (0.05) | 2.24E-05 | - | - | - | 7.84 (0.37) | 0.29 (0.05) | 1.17E-10 |
| **Mammals PhastCons** | - | **1.64%** | 8.96 (0.44) | 0.25 (0.07) | 2.25E-04 | 9.01 (0.44) | 0.35 (0.07) | 4.79E-07 | - | - | - |
| **Primates PhastCons** | - | **1.57%** | 11.03 (0.39) | 0.71 (0.05) | 6.13E-48 | 11.06 (0.39) | 0.71 (0.05) | 6.22E-49 | 11.10 (0.40) | 0.74 (0.05) | 1.19E-53 |

*S-LDSC conditional analyses to justify constraint annotations used in main analyses. We investigated the informativeness of three constraint annotations derived from phyloP scores in mammals, phastCons scores in mammals, and phastCons scores in primates. For consistency, each annotation represents 3.54% of the genome bases with the highest score (consistent with the FDR 5% threshold measured in phyloP scores in mammals). Using S-LDSC on 63 independent GWAS, we analyzed three models: Model 1 including all annotations; Model 2 including annotations derived from phastCons scores; and the Main Model (i.e. used in our main analyses) including annotations derived from phyloP scores in mammals and phastCons scores in primates. All analyses are conditioned to the baseline-LD model and six ENCODE3 annotations. We report the SNP-$h^2$ enrichments of the three constraint annotations, their normalized conditional effect sizes (Tau*), and their significance (Tau* P). We observed that the constraint annotation derived from phastCons scores in mammals was systematically less informative than annotations derived from phyloP scores in mammals and phastCons scores in primates.*

For further analyses, we created a "constrained in mammals" annotation by using phyloP scores at the FDR 5% threshold (i.e., phyloP scores ≥ 2.27), which represents 3.54% of the genome bases with the highest phyloP scores (*Supplementary Methods, Section 3*). We also created an "accelerated in mammals" annotation by considering SNPs with a phyloP score in mammals ≤ -2.270; we note that this annotation has a non-significant depletion of $h^2$ in (0.9±0.1 fold). To create a "constrained in primate" annotation, we considered SNPs with a phastCons score in primates ≥ 0.961, as it also covers 3.54% of the genome bases with the highest phastCons score in primates (*Supplementary Methods, Section 3*). We note that the phastCons Viterbi outputs defined 6.2% of the genome bases and 3.9% of common SNPs as constrained in primates; the corresponding annotation had an enrichment of 7.1±0.2x and was not considered in main analyses.

The final conditional analysis contained 106 total annotations: 97 annotations from the baseline-LD model, six ENCODE3 annotations, and three annotations from Zoonomia ("constrained in mammal", "accelerated in mammal" and "constrained in primate").

We partitioned constrained annotations using functional annotations from the baseline-LD model and from ENCODE3. We defined SNPs as *Exonic* by using the Coding annotation of the baseline-LD model. We defined SNPs as *Promoter* by using the union of two promoter annotations of the baseline-LD model and ENCODE3 promoters (PLS), and by excluding SNPs previously defined as *Exonic*. We defined SNPs as *Enhancer* by using the union of four enhancer annotations of the baseline-LD model and ENCODE3 pELS and dELS, and by excluding SNPs previously defined as *Exonic* or *Promoter*. We defined SNPs as *non-functional* by excluding SNPs previously defined as *Exonic*, *Promoter*, and *Enhancer*, but also by excluding SNPs with a posterior probability of being a QTL-SNP > 0.05 in the four fine-mapped QTL annotations of the baseline-LD model (*49*). Note that this functional annotation is independent of the UNICORNs defined in *Supplementary Methods, Section 8*.

We applied S-LDXR to the same 31 independent traits as in (*50*). We jointly analyzed the recommended baseline-LD-X model (62 total annotations, 28 main binary annotations), the constrained in mammals annotation, the constrained in primates annotation, and their intersection.

We partitioned SNP-heritability of low-frequency variants using S-LDSC with the baseline-LF model, as described in (*48*). Unlike previous analyses, we used 27 independent UK Biobank traits that have well-imputed low-frequency variants and that are powerful enough to partition low-frequency variant architectures (*48*). To compute mean standardized squared effects as a function of allele frequencies, we applied the following steps:

1. divided the functional annotation of interest into three low-frequency variant bins and three common variant bins,
2. ran S-LDSC with the baseline-LF model for 27 traits,
3. computed per-SNP heritabilities for 27 traits,
4. computed per-SNP squared effects by dividing per-SNP heritabilities by $2p(1- p)$, where $p$ is the allele frequency in UK10K (used here as a reference panel),
5. computed standardized squared effects of each bin by summing the standardized per-SNP squared effects with the bin, and computed corresponding standard errors using a Jackknife procedure, and
6. meta-analyzed standardized squared effects across the 27 traits.

#### ***Section 7: Functionally-informed fine-mapping***

By: BaDoi Phan, Steven Gazal

We applied functionally-informed fine-mapping (PolyFun), an approach for prioritizing candidate-causal variants from GWAS by accounting for LD and per-SNP heritability, on 24 well-powered UK Biobank diseases and complex traits (*51*, *52*) (mean $N$ = 440K; *table S12*). All analyses were performed using SuSIE as the underlying fine-mapper and on EUR ancestry UKBB imputed SNPs and LD matrices (*53*). We performed fine mapping using three progressive annotation sets to assess the informativeness of Zoonomia mammalian phyloP and primate phastCons to fine-map more confident candidate causal SNPs: 1) non-functional, fine-mapping using LD information on the SNPs alone, 2) baseline-LF, 187 annotations as previously published using the baseline-LF version 2.2.UKB set (*52*) , and 3) baseline-LF+Zoonomia, an updated set of 209 annotations comprising the baseline annotations, the Zoonomia annotations, six ENCODE3 annotations, and excluding 12 older primate and mammalian constrained annotations.

#### *Section 8: Constraint highlights functionally important SNP in unannotated and functionally validated regions*

By: Matthew Christmas, Jennifer R. S. Meadows, Xue Li

**UNannotated Intergenic COnstraint RegioNs (UNICORNs)** define binned regions of the genome with high constraint but which lack functional annotations (*Companion paper #1, FS1* for more detail). Briefly, The unannotated regions of the genome were defined subtractively (BEDTools v.2.29.2), removing GENCODE v37 (*12*) exons (UTRs and exons for all protein-coding genes) and promoters (TSS±1 kb), ENCODE3 (*22*) cCREs, DHS (including TF binding sites), ChIA-PET anchors, three promoter annotations (Promoter_UCSC, Human_Promoter_Villar, PLS_hg38), and six enhancer annotations (Enhancer_Andersson, Enhancer_Hoffman, Human_Enhancer_Villar, SuperEnhancer_Hnisz, dELS_hg38, pELS_hg38. Positions were lifted to hg38 where appropriate. For the remaining unannotated regions, bins of constraint were identified by first grouping any two phyloP constraint positions within 5 bp of each other, retaining clusters greater than 10 bp, giving a set of 424,179 UNICORNs. A set of non-constraint clusters was also generated to enable comparisons with all unannotated intergenic regions greater than 10 bp that did not contain constraint positions (phyloP < 2.27). This set generated over 6 million clusters, which were randomly down-sampled to a matched set of 424,179 elements.

We tested for enrichment of fine-mapped SNPs (*Supplemental Methods, section 7*) within UNICORNs, non-constrained unannotated regions (Unannotated), and the rest of the annotated genome (Other), by overlapping these regions with the available SNPs. The number of SNPs present in each region was 833, 5,895, and 305,599 respectively with mean PIP of 0.15, 0.05, and 0.05 respectively. The list of fine-mapped SNPs, their PIP score and UNICORN regions are listed in *table S13*. SNP PIP scores for each region were compared using linear regression in R v.4.0.1. This was performed independently for all traits within the fine mapped SNPs dataset, with FDR correction for multiple testing (*inline table*).

| Trait | Comparison | Estimate | Std Error | Statistic | P | # SNP | FDR |
|---|---|---|---|---|---|---|---|
| Forced Vital Capacity (FVC) | UNICORN vs. Other | -0.174 | 0.013 | -13.547 | 0.00E+00 | 15057 | 0.00E+00 |
| BMI | UNICORN vs. Other | -0.092 | 0.008 | -11.794 | 0.00E+00 | 26081 | 0.00E+00 |
| BMI | UNICORN vs. Unannotated | -0.099 | 0.009 | -11.317 | 0.00E+00 | 26081 | 0.00E+00 |
| Forced Vital Capacity (FVC) | UNICORN vs. Unannotated | -0.163 | 0.014 | -11.282 | 0.00E+00 | 15057 | 0.00E+00 |
| Years of Education | UNICORN vs. Other | -0.110 | 0.012 | -9.317 | 0.00E+00 | 9756 | 0.00E+00 |
| Height | UNICORN vs. Other | -0.117 | 0.013 | -9.272 | 0.00E+00 | 44096 | 0.00E+00 |
| Height | UNICORN vs. Unannotated | -0.126 | 0.014 | -9.206 | 0.00E+00 | 44096 | 0.00E+00 |
| Years of Education | UNICORN vs. Unannotated | -0.114 | 0.014 | -8.389 | 0.00E+00 | 9756 | 0.00E+00 |
| Bone mineral density (Heel T Score) | UNICORN vs. Other | -0.112 | 0.014 | -8.102 | 0.00E+00 | 28532 | 0.00E+00 |
| Smoking Status | UNICORN vs. Other | -0.100 | 0.013 | -7.438 | 0.00E+00 | 5331 | 0.00E+00 |
| Systolic Blood Pressure | UNICORN vs. Unannotated | -0.119 | 0.017 | -6.992 | 0.00E+00 | 17992 | 0.00E+00 |
| Resp. & ENT Diseases | UNICORN vs. Unannotated | -0.598 | 0.086 | -6.955 | 0.00E+00 | 2007 | 0.00E+00 |
| Resp. & ENT Diseases | UNICORN vs. Other | -0.577 | 0.084 | -6.893 | 0.00E+00 | 2007 | 0.00E+00 |
| Smoking Status | UNICORN vs. Unannotated | -0.100 | 0.015 | -6.686 | 0.00E+00 | 5331 | 0.00E+00 |
| Waist-hip Ratio | UNICORN vs. Other | -0.111 | 0.017 | -6.638 | 0.00E+00 | 14542 | 1.05E-10 |
| Bone mineral density (Heel T Score) | UNICORN vs. Unannotated | -0.095 | 0.015 | -6.347 | 2.23E-10 | 28532 | 6.68E-10 |

| | | | | | | | |
|---|---|---|---|---|---|---|---|
| Systolic Blood Pressure | UNICORN vs. Other | -0.100 | 0.016 | -6.135 | 8.69E-10 | 17992 | 2.45E-09 |
| Asthma | UNICORN vs. Other | -0.534 | 0.090 | -5.945 | 3.24E-09 | 2106 | 8.63E-09 |
| Asthma | UNICORN vs. Unannotated | -0.530 | 0.093 | -5.706 | 1.32E-08 | 2106 | 3.34E-08 |
| Eosinophil Count | UNICORN vs. Other | -0.196 | 0.036 | -5.457 | 4.90E-08 | 15498 | 1.18E-07 |
| FEV1-FVC Ratio | UNICORN vs. Other | -0.089 | 0.018 | -5.073 | 3.95E-07 | 19426 | 9.04E-07 |
| Eczema | UNICORN vs. Other | -0.358 | 0.072 | -5.002 | 5.93E-07 | 3932 | 1.29E-06 |
| Eosinophil Count | UNICORN vs. Unannotated | -0.181 | 0.037 | -4.875 | 1.10E-06 | 15498 | 2.29E-06 |
| Waist-hip Ratio | UNICORN vs. Unannotated | -0.086 | 0.019 | -4.559 | 5.18E-06 | 14542 | 1.04E-05 |
| Eczema | UNICORN vs. Unannotated | -0.328 | 0.075 | -4.367 | 1.29E-05 | 3932 | 2.49E-05 |
| FEV1-FVC Ratio | UNICORN vs. Unannotated | -0.071 | 0.019 | -3.731 | 1.91E-04 | 19426 | 3.53E-04 |
| Red Blood Cell Count | UNICORN vs. Other | -0.120 | 0.033 | -3.677 | 2.36E-04 | 19974 | 4.20E-04 |
| Platelet Count | UNICORN vs. Other | -0.099 | 0.027 | -3.622 | 2.92E-04 | 24639 | 5.01E-04 |
| Platelet Count | UNICORN vs. Unannotated | -0.091 | 0.028 | -3.232 | 1.23E-03 | 24639 | 2.04E-03 |
| Neuroticism | UNICORN vs. Other | -0.036 | 0.013 | -2.801 | 5.11E-03 | 5524 | 8.18E-03 |
| Morning Person | UNICORN vs. Other | -0.048 | 0.018 | -2.745 | 6.07E-03 | 4398 | 9.40E-03 |
| Red Blood Cell Count | UNICORN vs. Unannotated | -0.085 | 0.034 | -2.544 | 1.10E-02 | 19974 | 1.65E-02 |
| Hair Color | UNICORN vs. Other | -0.052 | 0.028 | -1.878 | 6.04E-02 | 7321 | 8.79E-02 |
| Neuroticism | UNICORN vs. Unannotated | -0.025 | 0.014 | -1.799 | 7.20E-02 | 5524 | 1.02E-01 |
| Hair Color | UNICORN vs. Unannotated | -0.051 | 0.030 | -1.712 | 8.70E-02 | 7321 | 1.19E-01 |
| Red Blood Cell Distribution Width | UNICORN vs. Other | -0.071 | 0.057 | -1.237 | 2.16E-01 | 14631 | 2.88E-01 |
| Morning Person | UNICORN vs. Unannotated | -0.021 | 0.021 | -1.020 | 3.08E-01 | 4398 | 3.99E-01 |
| Cardiovascular Diseases | UNICORN vs. Unannotated | -0.027 | 0.031 | -0.884 | 3.77E-01 | 7139 | 4.76E-01 |
| Type 2 Diabetes | UNICORN vs. Unannotated | -0.064 | 0.089 | -0.721 | 4.71E-01 | 1299 | 5.52E-01 |
| Type 2 Diabetes | UNICORN vs. Other | -0.062 | 0.083 | -0.739 | 4.60E-01 | 1299 | 5.52E-01 |
| White Blood Cell Count | UNICORN vs. Other | -0.025 | 0.033 | -0.738 | 4.61E-01 | 15873 | 5.52E-01 |
| Tanning | UNICORN vs. Unannotated | 0.032 | 0.056 | 0.580 | 5.62E-01 | 4619 | 6.42E-01 |
| Red Blood Cell Distribution Width | UNICORN vs. Unannotated | -0.031 | 0.058 | -0.523 | 6.01E-01 | 14631 | 6.70E-01 |
| White Blood Cell Count | UNICORN vs. Unannotated | -0.017 | 0.035 | -0.504 | 6.14E-01 | 15873 | 6.70E-01 |
| Hypothyroidism | UNICORN vs. Unannotated | -0.028 | 0.085 | -0.334 | 7.38E-01 | 2671 | 7.87E-01 |
| Tanning | UNICORN vs. Other | 0.011 | 0.053 | 0.211 | 8.33E-01 | 4619 | 8.69E-01 |
| Cardiovascular Diseases | UNICORN vs. Other | -0.004 | 0.030 | -0.142 | 8.87E-01 | 7139 | 9.06E-01 |
| Hypothyroidism | UNICORN vs. Other | -0.002 | 0.083 | -0.021 | 9.83E-01 | 2671 | 9.83E-01 |

Most fine-mapped SNP within UNICORNs were limited to a single trait (738/833), but others showed association to multiple traits (*inline table*), indicating that these variants lacking annotation could be used to further the biological understating of these traits. One variant, rs72782676, implicated in four traits was placed into genomic context with predicted TFBS (FS), and ENCODE3 regulatory annotations (cCREs, DHS). Further, allele frequencies for all SNPs in the region from TOPMed data freeze 8 were included, as well as mammalian phyloP and primate phastCons scores. Plots were generated using ggplot2 package in R v.4.0.1.

| Trait Number | Fine-mapped SNPs |
|---|---|
| 1 | 738 |
| 2 | 78 |
| 3 | 13 |
| 4 | 3 |
| 5 | 1 |
| Total | 833 |

**Massively parallel reporter assays (MPRAs)** are designed to interrogate the regulatory potential of hundreds to thousands of DNA sequences in a single high-throughput experiment. We tested the utility of mammalian phyloP to predict the strongest effect in MPRA reads. If successful, phyloP would first and foremost be benmarked as a tool to suggest causative function at base pair resolution, and could further be used to prioritise variants for more limited MPRA experiments. We accessed MPRA data targeted to four areas of genome regulation: eQTLs (*54*), 3'UTRs (*55*) and promoters and enhancers (*56*). Each data set represented different levels of experimental complexity in terms of construct size, completion of mutagenesis within the construct and number of cell lines assayed.

For the eQTL MPRA set (*54*), the coordinates for the 150 bp or genomic sequence for the constructs and the 29,108 unique variant positions were downloaded. These represent a combination of GWAS and eQTL hits drawn from lymphoblastoid cell lines (LCLs), with results drawn from populations of European and West African ancestry. For the 3'UTR MPRA, the construct contained a maximum of 101bp of genomic sequence, with 12,134 variants representing the signals of recent human evolutionary adaptation, or drawn from the results of GWAS analyses, some with linked eQTL effect. The MPRAs were performed in six cell lines, HEK293FT, HEPG2, HMEC, K562, LCL, and SKNSH. Genomic positions were lifted from hg19 to hg38, where appropriate, and annotated with mammalian phyloP scores. eQTL MPRA oligos were categorized into three groups based on activity: (1) Neutral, oligo with no regulatory activity for either allele; (2) Active, oligo with either allele showing regulatory activity; and (3) Skew, differential expression is observed between the two target alleles within an oligo. For the 3'UTR analysis (*55*), activity was extended to include any of the six cell lines. A t-test was performed to assess differences between the groups (eQTL, Fig. S8.1; 3'UTR, Fig. S8.2).

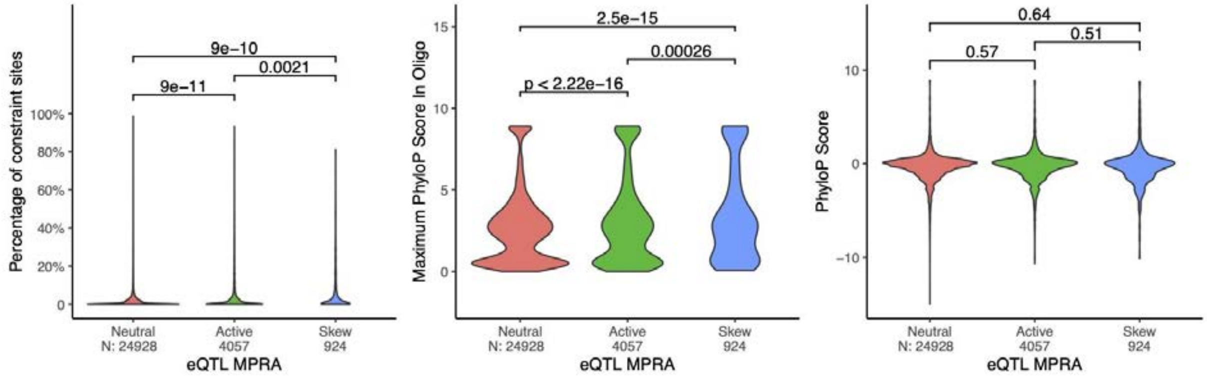

*Fig. S8.1. For the eQTL data, the relationship between constraint and regulatory potential (Neutral, Active, Skew) as (A) the percentage of bases, (B) the maximum phyloP score within an oligo, and (C) phyloP score at selectively tested base.*

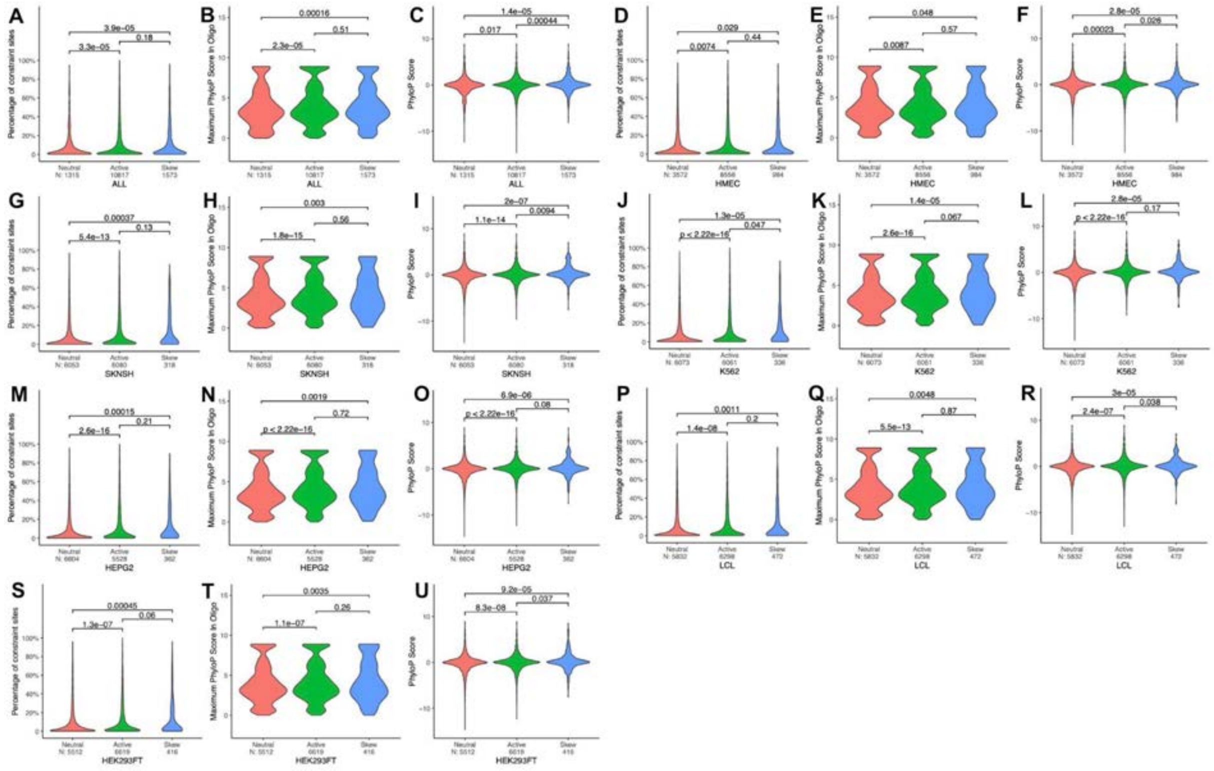

*Fig. S8.2. For the 3'UTR data, the relationship between constraint and regulatory potential (Neutral, Active, Skew) as the percentage of bases, the maximum phyloP score within an oligo, and phyloP score at selectively tested base is plotted for (A-C) All cell lines combined, (D-F) HMEC, (G-I) SKNSH, (J-L) K562, (M-O) HEPG2, (P-R) LCL and (S-U) HEK293FT cell lines independently.*

For both the eQTL and 3'UTR MPRAs, we calculated the enrichment of true expression-modulating variants at eight FDR cutoffs (FDR%/phyloP: 50/0.12, 30/0.90, 10/1.78, 5/2.27, 4/2.42, 3/2.60, 2/2.85, 1/3.27), by comparing these values to those of "all" variants considered in the specific MPRA set. The available number of variant positions varied per data set. For the eQTL MPRA set, it ranged from 29,108 for "all" group, to 420 for the "FDR 1%" group, while for 3'UTR MPRA, it ranged from 12,134 variants for the "all" group, to 471 for the "FDR 1%" group. To make the number of variants in each group comparable, we randomly selected 350 variants (eQTL MPRA) or 450 variants (3'UTR MPRA) in each group, calculated the percentage of expression-modulating variants and repeated this 1,000 times. We also calculated the standard deviation of the percentage of expression-modulating variants for each group. The fold enrichment was calculated as the percentage of true expression-modulating variants at FDR 5% divided by the percentage derived from the "all" group. This was 1.3 fold for 3'UTR MPRA, and 1.8 fold for eQTL MPRA (*Fig. S8.3*).

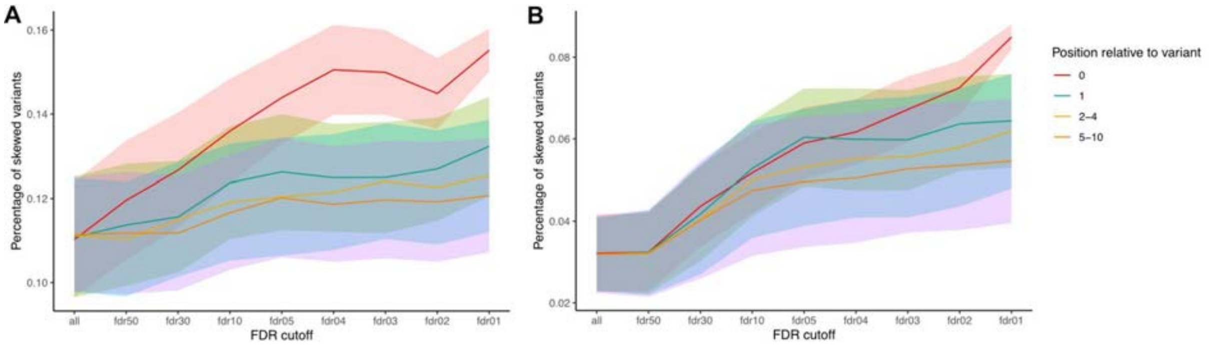

*Fig. S8.3. Enrichment and standard deviation of true expression modulating skewed variants captured at increasing FDR steps for (A) 3'UTR and (B) eQTL MPRA data sets.*

For the final regulatory space, the MPRA data from the error-prone PCR saturation mutagenesis of 9 enhancer and 11 promoter disease-associated regions, size ranging from 186 to 600 bp, was downloaded (*56*). That data set contained the functional measurements for over 30,000 single nucleotide substitutions and deletions, as each position from the region was mutated to the other three nucleotides or a deletion, and an MPRA effect read out reported. We calculated the average of the MPRA effects for each position to represent the relative ability of this position. These positions were subsequently annotated with mammalian phyloP scores, and the Spearman correlation between phyloP score and MPRA effect for each promoter region computed (*inline table*). The correlation ranged from negligible in the *FOXE1* locus (0.00) to highest in the *LDLR* locus(0.51), and so we also annotated the regions using ClinVar pathogenic variants and plotted the results for visual inspection (MPRA effect >2 *Fig. S8.4,* MPRA effect <2 *Fig. S8.5*). We note that high constraint is not indicative of correlated effect across the oligo and that cell line specific effects will alter the correlation (e.g. correlation ranges from 0.04 to 0.10 for the *TERT* locus in HEK293T and SF7996 cell lines).

| Regulatory Space | Name | Cell line | Spearman rho |
|---|---|---|---|
| Enhancer | *SORT1* | HepG2 | -0.09 |
| | *UCE UC88* | Neuro-2a | -0.02 |
| | *MYC* | LNCaP + 100nM DHT | -0.04 |
| | *MYC* | HEK293T | -0.01 |
| | *BCL11A* | HEL 92.1.7 | 0.06 |
| | *TCF7L2* | MIN6 | 0.10 |
| | *ZRS* | NIH/3T3 (with HOXD13/ HOXD13+H AND2) | 0.13 |
| | *RET* | Neuro-2a | 0.15 |
| | *ZFAND3* | MIN6 | 0.31 |
| | *IRF6* | HaCaT | 0.23 |
| | *IRF4* | SK-MEL-28 | 0.35 |
| Promoter | *FOXE1* | HeLa | 0.00 |
| | *Factor IX (F9)* | HepG2 | 0.02 |
| | *TERT* | HEK293T | 0.04 |
| | *MSMB* | K562 | 0.04 |
| | *HNF4A (P2)* | HEK293T | 0.13 |

| | | |
|---|---|---|
| *TERT* | SF7996 | 0.10 |
| *GP1BB* | HEL 92.1.7 | 0.05 |
| *HBB* | HEL 92.1.7 | 0.19 |
| *HBG1* | HEL 92.1.7 | 0.19 |
| *PKLR* | K562 | 0.21 |
| *LDLR* | HepG2 | 0.51 |

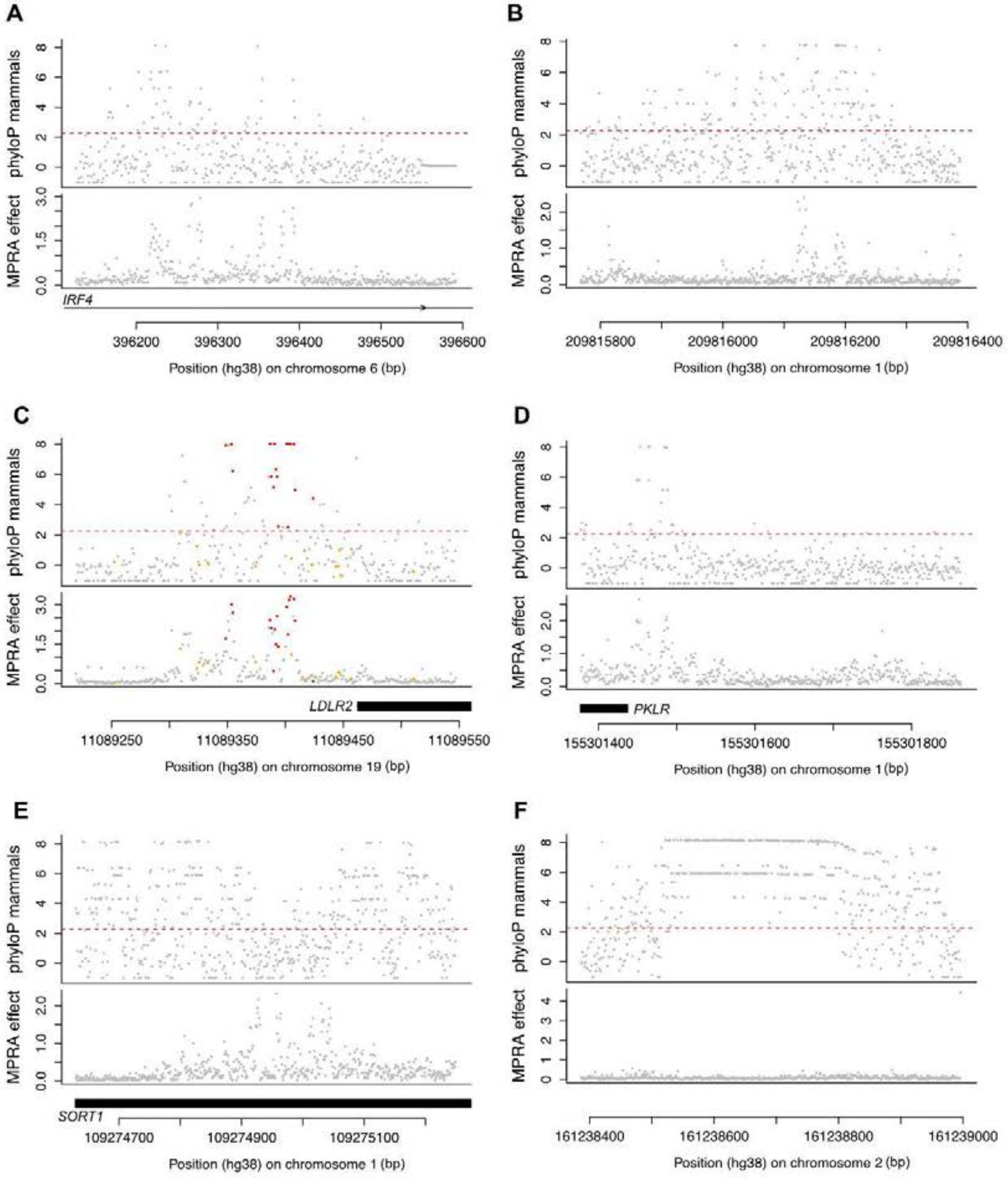

*Fig. S8.4. PhyloP score plotted in comparison to MPRA effect. Constrained (red), and unconstrained (orange) ClinVar pathogenic variants are plotted to highlight known deleterious positions where available. Loci in this comparison have at least one variant with MPRA effect greater than 2. A) IRF4, B) IRF6, C) LDLR2, D) PKLR, E) SORT1, F) UC88.*

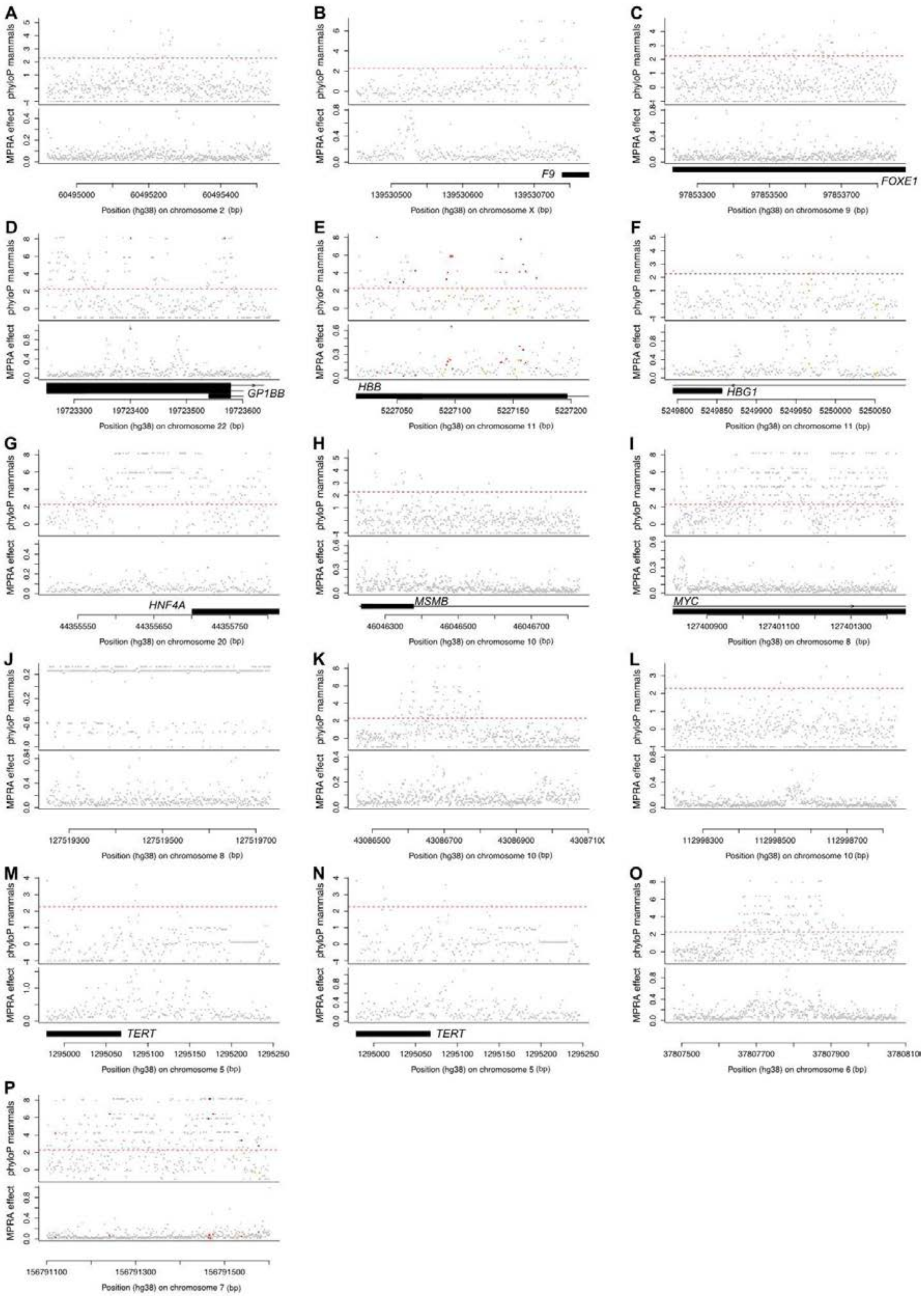

*Fig. S8.5. PhyloP score plotted in comparison to MPRA effect. Constrained (red), and unconstrained (orange) ClinVar pathogenic variants are plotted to highlight known deleterious positions where available. Loci in this comparison have an MPRA effect less than 2. A) BCL11A, B) F9, C) FOXE1, D) GP1BB, E) HBB, F) HBG1, G) HNF4A, H) MSMB, I) MYC focused on rs6983267, J) MYC focused on rs11986220, K) RET, L) TCFL2, M) TERT in cell line GBM, N) TERT in cell line HEK, O) ZFAND3, P) ZRSL.*

***Section 9: Creation of a Gene-Based Measure of Constraint***
By: Patrick Sullivan, Quan Sun, Yun Li

Measures of the degree to which a gene is constrained across mammalian evolution are often used to help understand the impact of genetic variation on human disease. Measures of gene-based constraint include pLI (*7*), LOEUF (*8*), and RVIS (*57*). Developing a measure of gene constraint is complicated by the extraordinary diversity of human genes. In this section, we describe and validate measures of gene constraint based on the Zoonomia alignment.

Human gene models were based on GENCODE v36 (11/2020) (*12*), and the genome build was hg38/GRCh38. *Supplementary Methods, Section 4* shows data sources and definitions. We conducted extensive checks for each protein-coding (PC) transcript, and removed those flagged as incomplete or containing an error like a mismatch between the number of codons and amino acids. The final list contained 196,073 transcripts.

The most common transcript biotype was PC (29.7%), and there were 58,272 PC transcripts from 19,397 PC genes (chr1-22, chrX, and chrY). There was a median of two PC transcripts per gene (IQR 1-4, range 1-86). Non-independence is a significant issue as 64.4% of CDS bases are in multiple transcripts. Although our main interest was PC genes, we determined the fraction of cDNA exons for transcripts of all biotypes that were constrained in 240 mammals (bases with phyloP ≥ 2.270) and in 43 primates (bases with phastCons ≥ 0.961).

We compared multiple measures of evolutionary constraint in PC genes (pickOne subset, 19,397 PC genes/transcripts, *Supplementary Methods, Section 4*). Beginning with the simpler methods, *(1)* was the fraction of CDS bases constrained in mammals (phyloP ≥ 2.270), and *(2)* was the fraction of CDS bases constrained in primates (phastCons ≥ 0.961).

These simple measures may not capture many important features of PC genes given that codons, not individual bases, are the main informational unit in a PC gene. As illustrated in *Fig. S9.1*, codons vary considerably in usage over all CDS. The next two measures of gene constraint were relative and compared every CDS base (in its reading frame defined by codon and position) to the same codon-positions in all other PC codons. For all 192 codon-positions (=64 codons x 3 positions each), we computed phyloP summary statistics (mean, sd, median, and IQR) over 32.13 million CDS bases and 10.71 million codons, For example, for the 173,162 uses of the Ala codon GCA, positions 1-3 had mean (sd): 4.88 (3.64), 5.38 (3.45), and -1.85 (3.623). For each PC gene, we computed *(3)* the mean normalized phyloP score at each CDS base by subtracting the mean for that codon-position and dividing by the standard deviation, and then *(4)* the median robust normalized phyloP score at each CDS base (subtract median, divide by IQR).

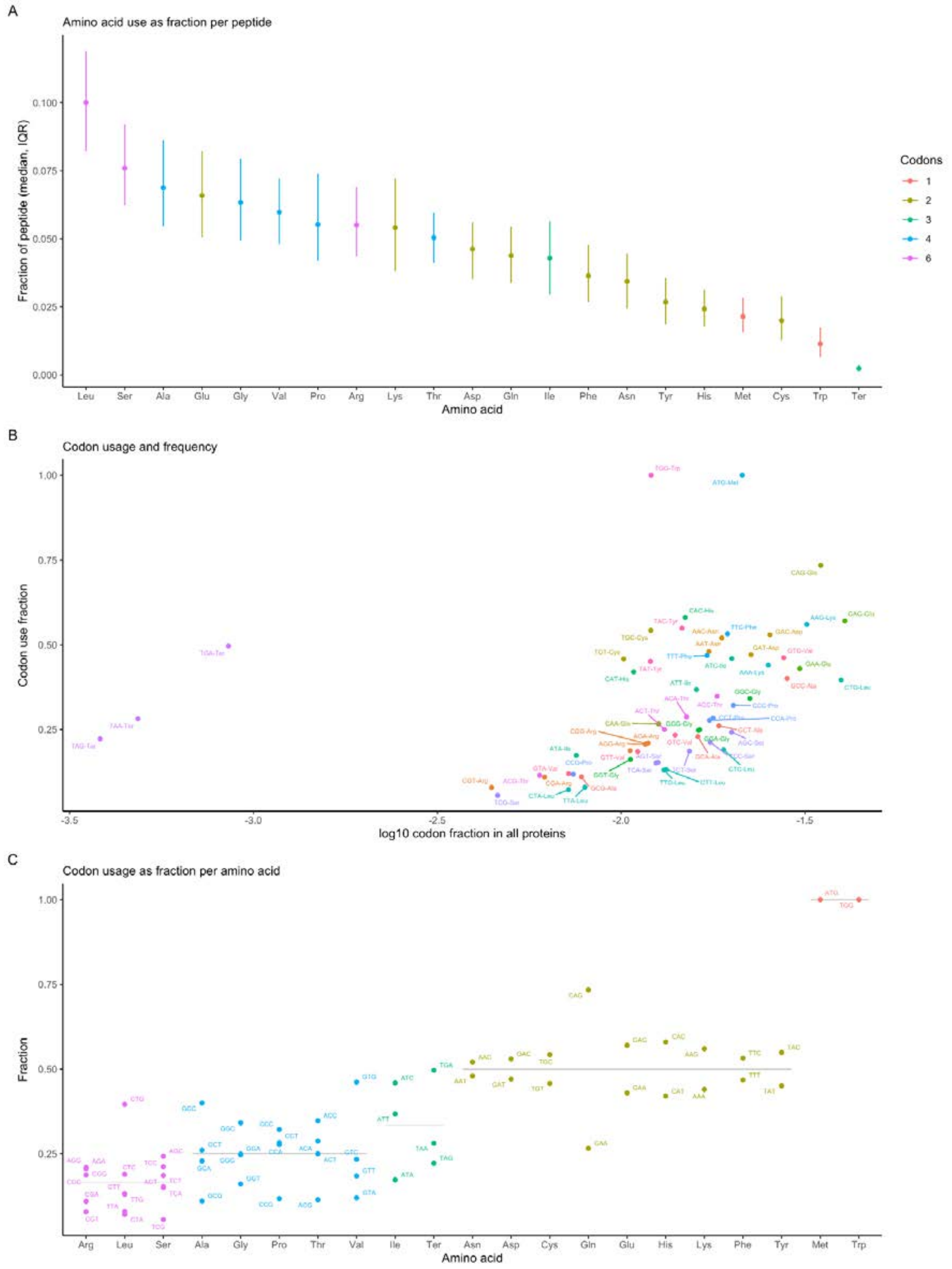

*Fig. S9.1. (A) Distribution of amino acids per protein (pickOne transcript set). X-axis=20 amino acids (3 standard 3 letter code) plus stop codon. Y-axis=fraction of a peptide that is each amino acid. Points are medians and whiskers are IQR. The coloring shows the number of codons per amino acid (range 1-6). Peptide amino acid content ranges from a median of 1.27% (Trp) to 10.10% (Leu). (B) Codon usage. X-axis=log$_{10}$ fraction over all peptides and the Y-axis is codon usage. The frequencies of codon usage are markedly variable. (C) Codon usage bias. X-axis=amino acids (sorted and colored by number of codons, 6, 4, 3, 2, and 1). Y-axis=fractional usage of codons per amino acid. Points show codons per amino acid. Grey lines are expectations under equal*

*codon usage (i.e., ⅙, ¼, ⅓, ½, and 1). Except for Met and Trp (one codon each), all codons deviate from equal usage, most notably for Leu-CTG, Val-GTG, Ter-TGA, and Gln-CAG.*

We next applied a model-based approach to adjust phyloP scores for a range of coding features with an impact on the CDS base constraint. The dependent variable was the phyloP score for each CDS base (median 3.6 over the pickOne set of 32.13 million CDS bases). The *inline table* below defines 12 covariates and gives the median phyloP values associated with the covariate levels. These included: codon data (start codon, stop codon, position in codon, the codon string, and the specific base), mutational consequences (of the 3 possible mutations at a base, the number of synonymous mutations and whether a mutation to stop could occur), positional features (coding base located at an inner edge of an exon, the number of PC transcripts including a given base, whether a base appeared in multiple reading frames), whether the base had a notably small number of species aligning, and, to capture local constraint, the rolling mean phyloP score (±9 bases excluding the current base).

| *Name* | *Definition* | *phyloP descriptive statistics* |
|---|---|---|
| phyloP.am | Dependent variable, phyloP score per coding base | Median: 3.6 |
| startCodon | Initial codon (Met, ATG), TRUE/FALSE | Median: TRUE=6.2 |
| stopCodon | Terminal codon (TAA, TAG, TGA) , TRUE/FALSE | Median: TRUE=4.4 |
| exonEdge | Coding base is at 5’ or 3’ exon edge, TRUE/FALSE | Median: TRUE=6.3 |
| coding_base | Coding base (A, C, G, T) | Median: A=2.9, C=3.6, G=4.7, T=2.9 |
| codonPos | Base position in codon (1, 2, 3) | Median: 1=5.3, 2=6.2, 3=0.7 |
| NspeciesLow | Coding base with ≤ 110 species aligning (5% of all CDS), TRUE/FALSE. For reference, median species per CDS base 237, IQR 231-240 | Median: TRUE=0.3 |
| mut_to_self | For the 3 possible mutations of a base, number of synonymous mutations (0-3). Four-fold degenerate sites have value 3 | Median: 0=6.1, 1=1.4, 2=0.9, 3=0.3 |
| mut_to_stop | For 3 possible mutations of a base, any mutation to stop, TRUE/FALSE | Median: TRUE=6.1 |
| cdsUse | Number of PC transcripts including a CDS base (range 1-85), coded 0-4, ≥5 | Median: 1=3.0, 2=3.5, 3=3.7, 4=3.9, ≥5=4.7 |
| cdsFrame | CDS base used in >1 reading frame (0.88%), TRUE/FALSE | Median: TRUE=3.7 |
| phyloPrm19ex | Excluding current base, rolling mean phyloP in window 19 bases (± 9 bases) | – |
| codonString | Codon in which a base lies (64 levels, AAA to TTT) | Median range: Thr-ACA 1.3 to Trp-TGG 6.9 |

A multivariable linear model was fit with $R^2$ of 0.547. Of the 76 terms in the model, only 2 were non-significant, 74 had P-values < 1e-20, and 56 had |Z-score| > 40 and a P-value smaller than the operating system’s minimum numeric value of 2.2e-308). The most impactful terms in the model (|Z-score| > 300) are in the *inline table* below. There were notable independent effects for codon position 1 and 2 (vs position 3) and if a coding base was C or G (vs A). The local context (±9 bases) was also important as were the mutational consequences. *(5)* The gene constraint metric was the mean adjusted phyloP score over all CDS bases per PC gene.

| *term* | *estimate* | *se* | *statistic* | *P* | *Comment* |
|---|---|---|---|---|---|
| codonPos = 2 | 2.58 | 2.16e-3 | 1195 | <2e-308 | Codon position (vs 3), consistent with highest constraint overall |
| codonPos = 1 | 2.03 | 2.08e-3 | 979 | <2e-308 | Consistent with second highest constraint overall |
| coding_base = G | 2.00 | 1.58e-3 | 1260 | <2e-308 | Coding base (versus A), consistent with GC enrichment in CDS and in constrained bases |
| coding_base = C | 1.84 | 1.65e-3 | 1112 | <2e-308 | As above |
| exon_edge = TRUE | 1.29 | 4.15e-3 | 310 | <2e-308 | Exon bases at the sides of an exon (adjacent to intronic splice sites) |
| phyloPrm19ex | 0.89 | 2.57e-4 | 3475 | <2e-308 | Local constraint, rolling mean phyloP score ±9 bases excluding central base ("19" is 9 + 9 + 1) |
| mut_to_stop = TRUE | 0.58 | 1.82e-3 | 321 | <2e-308 | Any mutation in this position yields stop codon |
| mut_to_self (0-3) | -0.89 | 9.23e-4 | -967 | <2e-308 | Number of synonymous mutations at this site |

The informational units in PC genes are codons not individual base scores, and the above approaches may not completely capture the complexities of codon constraint—in many instances, evolutionary selection will operate on amino acids and not necessarily individual bases. As a complementary approach, we thus evaluated the extent to which each human amino acid was constrained across species. The TOGA multiple species alignment provides peptide comparisons for 442 vertebrate species (including all 240 Zoonomia species). All species are aligned to the human reference for each transcript and each amino acid. We used the R package *Bio3D (58)* to compute the normalized Shannon entropy score for each amino acid (ranging from 0=high entropy/not constrained to 1=low entropy/highly constrained). This measure was based on the standard amino acid alphabet. We also evaluated collapsing amino acids into 10 groups by physiochemical properties (e.g., hydrophobic/aliphatic, aromatic, polar, etc), and found highly correlated results. *(6)* The gene constraint metric based on the TOGA multiple species protein alignment was the mean normalized Shannon entropy score over all CDS bases.

*(7)* From *Supplementary Methods, Section 4*, we included the fraction of common, constrained CDS SNPs for each PC gene given the results above suggesting its salience for identifying constraint.
In summary, we compared 9 gene constraint metrics in the pickOne transcript set. For each PC gene, we calculated:

1. *fracCdsConsFdr05*, number of CDS bases with mammalian phyloP ≥ 2.27 divided by the number of CDS bases
2. *fracCdsConsPri*, number of CDS bases with primate phastCons ≥ 0.96 divided by the number of CDS bases
3. *phylopNormMean*, mean of normalized phyloP score in CDS bases (vs all codon-position phyloP scores)

4. *phylopNormMeanRobustP50*, median of robust normalized phyloP scores in CDS bases (vs all codon-position phyloP scores)
5. *phylopFittedMean*, mean of phyloP scores in CDS bases after adjustment for 12 covariates in a linear model
6. *HnormMean*, mean normalized Shannon entropy score from the TOGA multiple species protein alignment
7. *fracCdsConsCommon*, number of common, constrained CDS SNPs divided by the number of CDS bases
8. For comparison, we included the gnomAD *LOEUF* and *pLI* scores for the same PC transcripts (*8*) derived from large exome sequencing studies, and use different methods to contrast the observed number of predicted loss-of-function variants to a model-based expectation per transcript

The *inline table* below shows summary statistics for these 9 measures along with the number of CDS bases (N=19,387 genes, pickOne transcript set). Greater values indicate higher constraint except for LOEUF and fracCdsConsCommon which are inverted. The completion rates are generally high, but HnormMean, LOEUF, and pLI are missing for >10% of genes. The univariate histograms for most measures show reasonable spread except for fracCdsConsCommon (point mass at zero) and pLI (essentially bimodal).

| *measure* | *complete* | *median* | *p25* | *p75* | *histogram* |
|---|---|---|---|---|---|
| cdsBases | 1.000 | 1284 | 807 | 2073 | ▇▁▁▁▁ |
| fracCdsConsFdr05 | 1.000 | 0.620 | 0.462 | 0.724 | ▁▁▃▇▁ |
| fracCdsConsPri | 1.000 | 0.653 | 0.378 | 0.840 | ▃▃▅▆▇ |
| phylopNormMean | 0.951 | 0.037 | -0.343 | 0.337 | ▁▁▃▇▁ |
| phylopNormRobustp50 | 0.951 | -0.007 | -0.336 | 0.091 | ▁▁▂▇▇ |
| phylopFittedMean | 0.951 | 3.642 | 2.668 | 4.402 | ▁▁▃▇▁ |
| HnormMean | 0.889 | 0.498 | 0.349 | 0.629 | ▂▅▇▅▁ |
| fracCdsConsCommon | 0.995 | 0.001 | 0.000 | 0.001 | ▇▁▁▁▁ |
| LOEUF | 0.899 | 0.883 | 0.483 | 1.306 | ▇▇▇▆▆ |
| pLI | 0.899 | 0.002 | 0.000 | 0.472 | ▇▁▁▁▂ |

*Fig. S9.2A* shows the Spearman pairwise correlations among these measures. There were small correlations with the number of CDS bases with the notable exception of LOEUF which was strongly correlated with the number of coding bases (rho = -0.51). This suggests that LOEUF is impacted, and perhaps biased, toward informativeness in larger CDS transcripts. The main gene constraint measures derived from Zoonomia (fracCdsConsFdr05, phylopNormMean, phylopNormRobustp50, and phylopFittedMean) were highly intercorrelated (Spearman rho ≥ 0.95), and there were high correlations with fracCdsConsPri (rho ≥ 0.87) but far lower correlations for HnormMean (rho 0.45-0.52). fracCdsConsCommon was not informative due to lack of variation.

As shown in *Fig. S9.2B*, the relation of fracCdsConsFdr05 with the normalized and adjusted measures of gene constraint were reasonably linear across the X-axis range. A subset of genes with low fracCdsConsFdr05 values (<0.2) had alternative gene metrics that were shifted lower (i.e., lesser constraint). Because of these strong relationships and given its simple calculation, we selected *fracCdsConsFdr05* (or *fracCdsCons*) as the gene constraint metric.

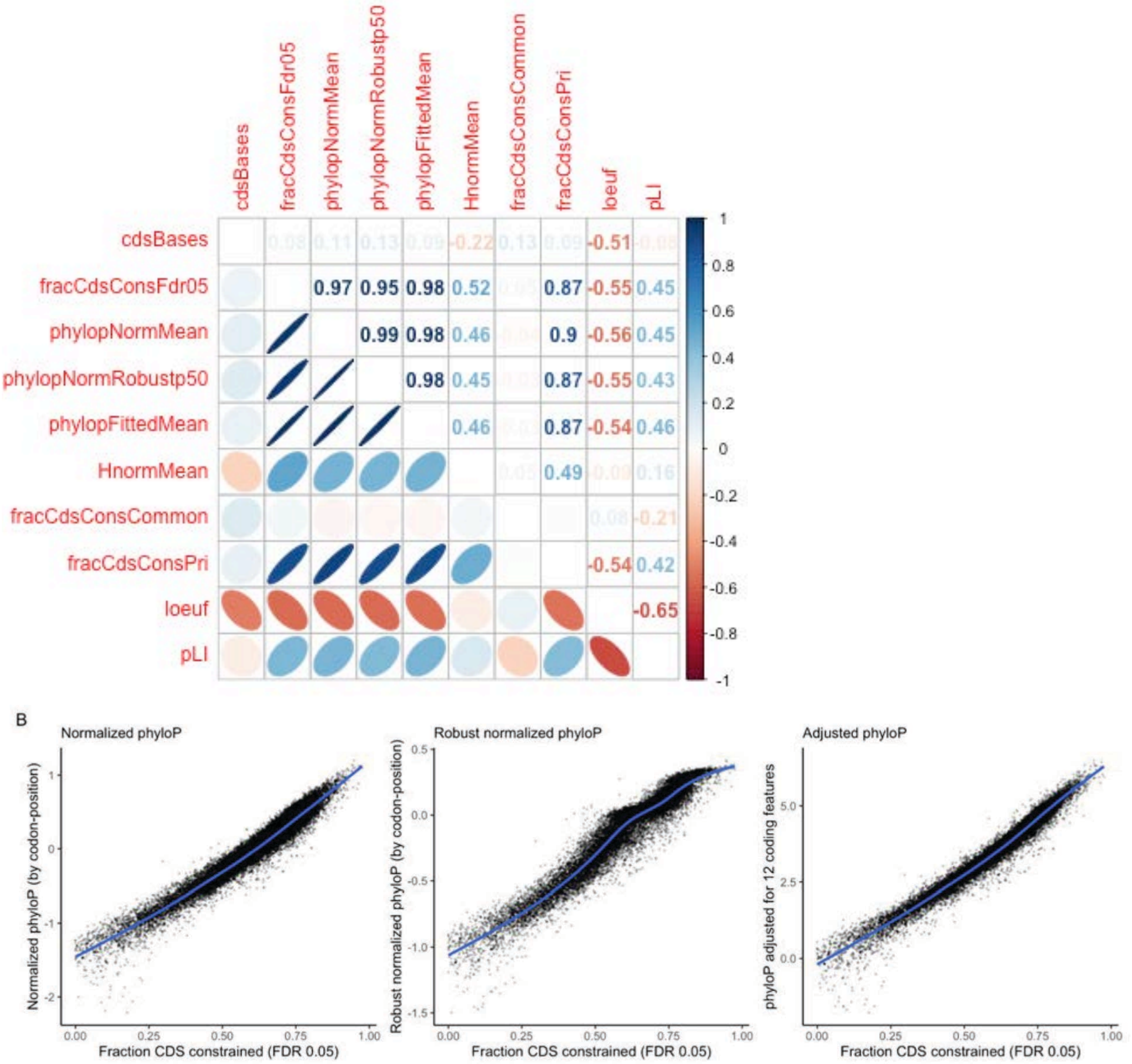

*Fig. S9.2. (A) Spearman correlations of the gene constraint metrics and CDS size. (B) Scatterplots of the data underlying Spearman correlations. X-axis=fracCdsConsFdr05, Y-axis=three other evolutionary constraint measures. Each point is a PC gene. The blue line is a lowess smoother. We chose to use fracCdsConsFdr05 as it contains essentially the same information as the other phyloP-based measures and for its ease of calculation.*

**Exon constraint.** The above analyses apply to CDS (coding bases), exons whose bases can be translated into peptides. Many other genes are composed of non-coding exons that can be transcribed into cDNA. We also created a gene-based metric for cDNA which can be calculated for any biotype. *fracExonCons* is the fraction of cDNA bases that are constrained in mammals. For PC genes, the Spearman rho of *fracCdsCons* (CDS) with *fracExonCons* (CDS and UTRs) was 0.64. *Fig. S9.3* shows box plots for five biotypes. Most non-coding genes have low median levels of constraint but a fraction have high levels of constraint. Constraint can be used to identify non-coding genes with high levels of mammalian constraint as one index of potential importance.

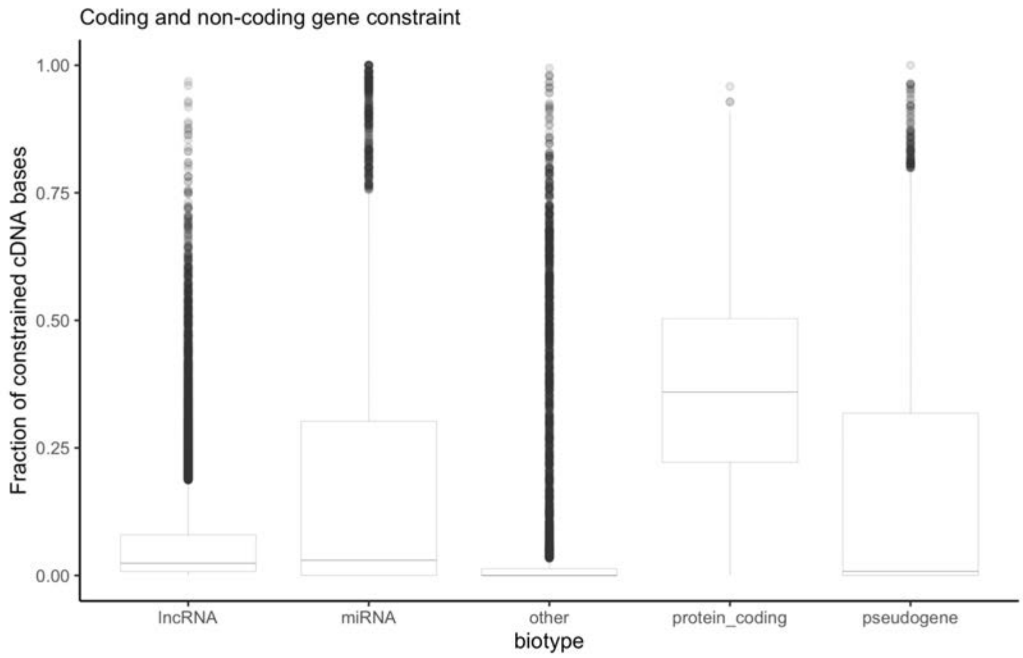

*Fig. S9.3. Box plots of the distributions of fracExonCons for five biotypes.*

*Section 10: Evaluation of a Gene-Based Measure of Constraint*

By: Patrick Sullivan, Steven Gazal

**Identifying genes whose constraint is measurable:** Human PC genes are highly diverse. No single measure of gene constraint will be applicable to all genes. We sought an empirical approach to identify PC genes for which an evolutionary gene constraint metric is applicable. As detailed in *Supplementary Methods, Section 9*, the gene constraint metric is *fracCdsCons* (number of CDS bases with phyloP scores ≥ 2.270 divided by the total number of CDS bases). For each PC gene (pickOne transcripts, N=19,386), we created four annotations: fracCdsCons, fracConsPr (the fraction of CDS bases constrained in 43 primates, phastCons ≥ 0.961), the fraction of CDS bases without a phyloP score, and the fraction of CDS bases with a low number of species aligning (≤110). The latter two measures capture difficult to align coding bases across 240 mammalian species (i.e., due to error, complexity, and sequence specific to primates or humans). Adapting scRNA-seq methods, we applied UMAP (*59*) as a dimensional reduction algorithm and then used DBSCAN to identify clusters in the results (*60*) (*Fig. S10.1*).

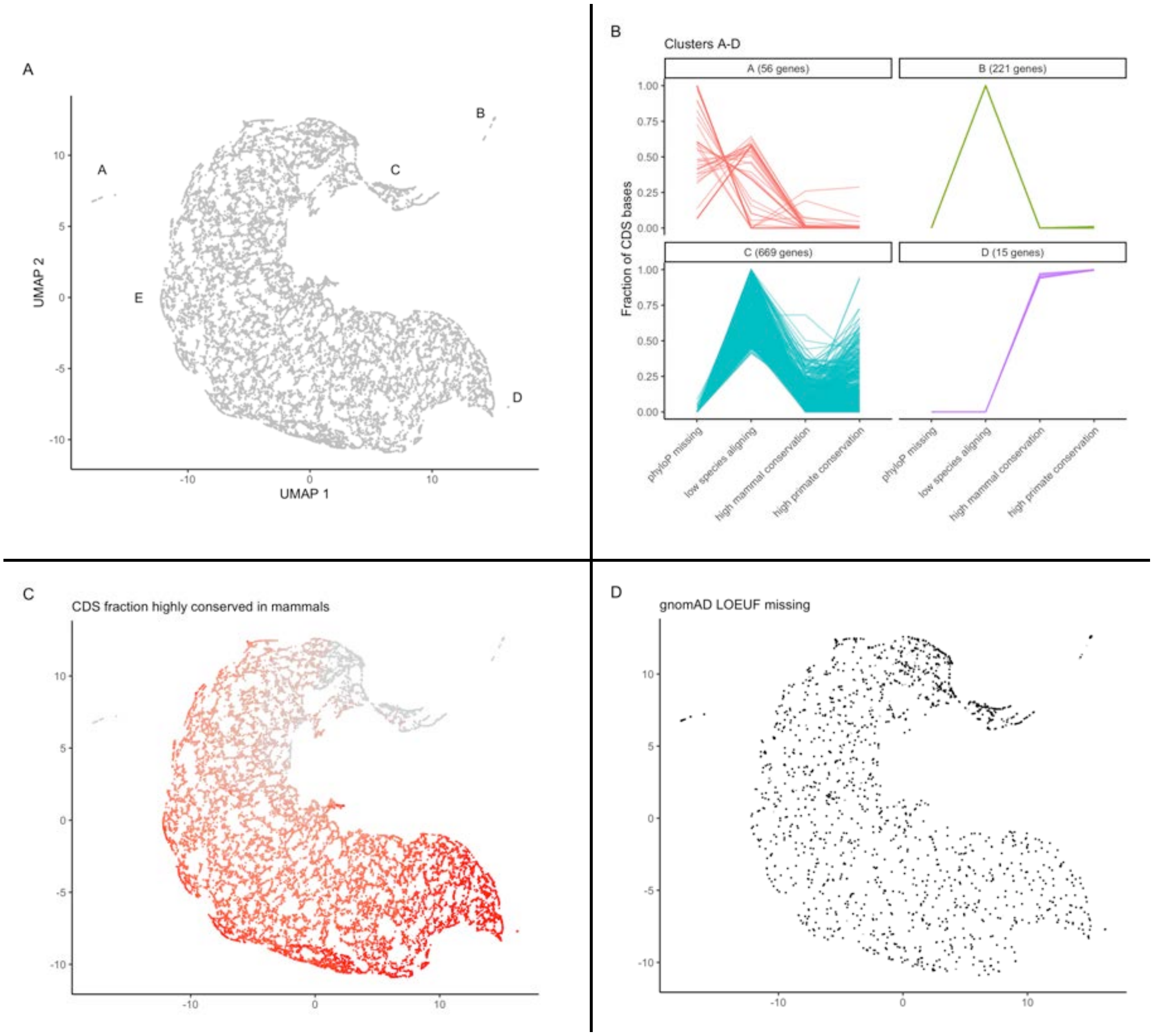

*Fig. S10.1. (A) Results from UMAP dimensional reduction and five clusters from DBSCAN (labeled A-E). (B) Parallel plot of outlier clusters A-D. (C) Mammalian constraint (fracCdsCons, grey=least, red=most). (D) Genes with missing gnomAD LOEUF scores.*

These analyses yielded an interpretable high-level classification of human PC genes with respect to mammalian evolutionary features. The *inline table* describes the clusters. We evaluated human ortholog data (ENSEMBL, 1:1 high-confidence human orthologs from 73 species including 23 primates) (*61*).

| *Cluster* | *PC genes* | *fracCdsCons p50* | *phyloP missing* | *Species aligning low* | *constrained CDS* | *Human/primate only* |
|---|---|---|---|---|---|---|
| A | 56 | 0 | 49/56 >10% | 30/56 >10% | 2/56 >10% | High (91%, 51/56) |
| B | 221 | 0 | None | 221/221 = 100% | 221/221 = 0% | High (90%, 199/221) |
| C | 669 | 0.04 | 0/669 >10% | 669/669 > 10% | 178/669 >10% | High (76%, 508/669) |
| D | 15 | 0.95 | None | None | 15/15 >94% | Low (7%, 1/15) |
| E | 18,425 | 0.63 | 5/18,425 >10% | 714/18,425 >10%) | Variable (0-93%) | Low (9%, 1605/18425) |

*Cluster E* contained most genes (18,425) and the other 4 clusters were different types of outliers. The 56 genes in *Cluster A* did not map well and were poorly constrained (e.g., *GAGE**, *NBPF*, USP17L** and several olfactory receptors). The 221 genes in *Cluster B* mapped well but were not constrained, and 90% had no ortholog or an ortholog only in primates, a pattern suggestive of a human- or primate-specific gene. *Cluster C* (669 genes) was notable for low numbers of species with CDS alignment (e.g., five *HLA* genes and 14 interferon alpha genes). The 15 genes in *Cluster D* mapped well and were very highly constrained across mammals (especially in primates, fraction CDS constrained > 0.993, multiple *HOX** genes).

We conducted hypergeometric gene set analyses for outlier Clusters B and C (outlier clusters with >200 genes). Genes in Cluster B showed significant enrichment for gene sets involved in: (a) olfaction, (b) epidermis development, and (c) male gamete generation. Genes in Cluster C showed significant enrichment for: (a) adaptive immunity, (b) olfaction, and (c) epidermis development. A common theme in Clusters B and C is the position of these genes at the interface between a mammal and its external milieu, and these may evolve for a mammal to adapt to specific environments. *Fig. S10.1C* shades each gene from light grey (fracCdsCons = 0) to red (fracCdsCons = 1). There is a gradient in mammalian constraint from the top middle anticlockwise to the lower right.

Interpretation: Cluster A contains genes whose CDS bases are in complex genomic regions that align poorly; Clusters B and C contain genes that evolved during mammalian evolution in response to environment; Cluster D contains the most highly constrained genes; and Cluster E are all other genes. We retained Clusters C, D and E as sufficiently comparable genes and viewed Clusters A and B as not having interpretable phyloP-based gene constraint.

**Gene set analysis of most and least constrained genes:** We conducted gene set analysis for 19,109 PC genes (Clusters C, D, and E) in the top or bottom 5% (ventile) of the fraction of mammalian gene constraint (fracCdsCons, overall median 0.623, IQR 0.471-0.725, range 0-0.975). Gene set analyses were conducted for genes in the top 5% (955 genes, fracCdsCons 0.811–0.975) and the bottom 5% (956 genes, fracCdsCons 0–0.150). Gene set analysis (hypergeometric test, background all PC genes, Bonferroni-correction to $P < 0.005$) were conducted for Gene Ontology (GO) pathways, for genes with high expression specificity across over ~1,200 human cell types (via single-cell RNA-seq, Human Cell Landscape, HCL), and the expert-curated set of synaptic genes (SynGO) (*62–64*). The results are summarized in the *inline table* below.

| *Genes* | *Gene sets* | *Summary of significant pathways* ($P_{bonf} < 0.005$) |
|---|---|---|
| Bottom 5% | GO | defense response (adaptive immunity, bacteria/virus, cell killing, cytokine/interferon) |
| | | taste (bitter), smell |
| | | skin development, keratinization |
| | SynGo | None |
| | HCL | None |
| Top 5% | GO | basic embryology (stem cell proliferation/differentiation, tube formation, anterior/posterior patterning, endoderm/mesoderm formation) |
| | | organ morphogenesis (central nervous system, peripheral nervous system, connective tissue, ear, epithelium, eye, gastrointestinal tract, heart, kidney, lung, muscle, myeloid, pancreas, skeleton) |
| | | cell cycle (phase transition, fate, WNT), cell signaling, positive and negative regulatory processes |
| | SynGO | Presynapse and postsynaptic processes (synapse assembly, postsynaptic density, neurotransmitter regulation, synaptic vesicle cycle, modulation of transsynaptic signaling) |
| | HCL | Fetal only. Neurons (intestine, kidney, lung, pancreas, spinal cord, stomach); retinal progenitor; male leydig cells; fibroblasts |

Gene set analyses of the *bottom ventile* (lowest mammalian constraint) were dominated by genes at the interface of a mammal with its environment: microbiota defense, taste and smell, and multiple skin processes. These gene sets are central to the adaptive evolution of a mammal to its environment—i.e., the specific microbiota, adaptations of smell and taste to detect prey, predators, and poisons, and adaptations of skin for temperature regulation, camouflage, and defense.

Gene sets in the *top ventile* (highest mammalian constraint) were notably fundamental to mammalian development. Significant gene sets were involved in basic embryology, morphogenesis of most organ systems, cell cycle, and basic synaptic processes.

**Mammalian constraint and external validators:** Following part of the validation strategy for gnomAD LOEUF (*8*), we evaluated the pattern of constraint with respect to external gene sets with established patterns of constraint. We repeated the analyses previously reported for LOEUF in comparison to fracCdsCons (coding both as deciles, 0=least and 1=most constrained).

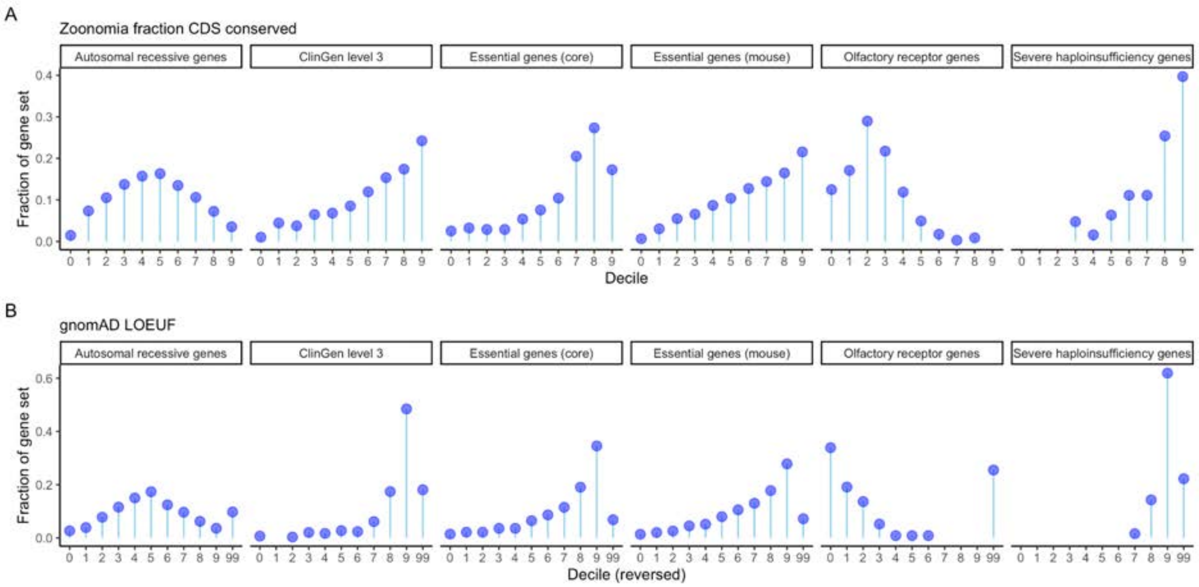

*Fig. S10.2. Gene constraint deciles versus external gene sets as "lollipop plots". Each panel has 6 subgraphs for autosomal recessive genes, ClinGen level 3 genes, essential genes from Hart, essential genes in mouse, olfactory receptors, and severe haploinsufficiency genes. X-axis-=constraint decile (0=least,, 9=most constrained, 99=missing). Y-axis=circles are the fraction of the PC genes in a gene set in each decile (A) Zoonomia, fraction CDS constrained. (B) Recapitulates Figure 3 from the gnomAD paper with LOEUF decile reversed and showing missing data.*

The results are generally similar between *Figs. S10.2A* and *S10.2B*. Autosomal recessive genes peak in the mid-range of constraint. The fraction of genes with strong evidence of pathogenicity, essential genes, and severe haploinsufficiency increased with degree of constraint. Olfactory genes were concentrated in the least constrained groupings. On the other hand, often sizable fractions of genes in these categories had no LOEUF score (the "99" category). The patterns were perhaps slightly cleaner for fracCdsCons (especially autosomal and essential genes in mouse).

**Constraint and human disease and gene expression:** We next evaluated the relevance of mammalian gene constraint scores to human disease. *Fig. S10.3A* shows the relationship of fracCdsCons multiple human disease annotations. For all comparisons, increasing constraint is correlated with increasing relevance for human disease. *Fig. S10.3B* depicts the relation with GTEx gene expression, and greater gene constraint is correlated with greater expression in all tissues. "Housekeeping" genes that are uniformly expressed across tissues had greater constraint ($P < 3x10^{-197}$, 3.0% of the least constrained decile and 30.5% of the most constrained decile).

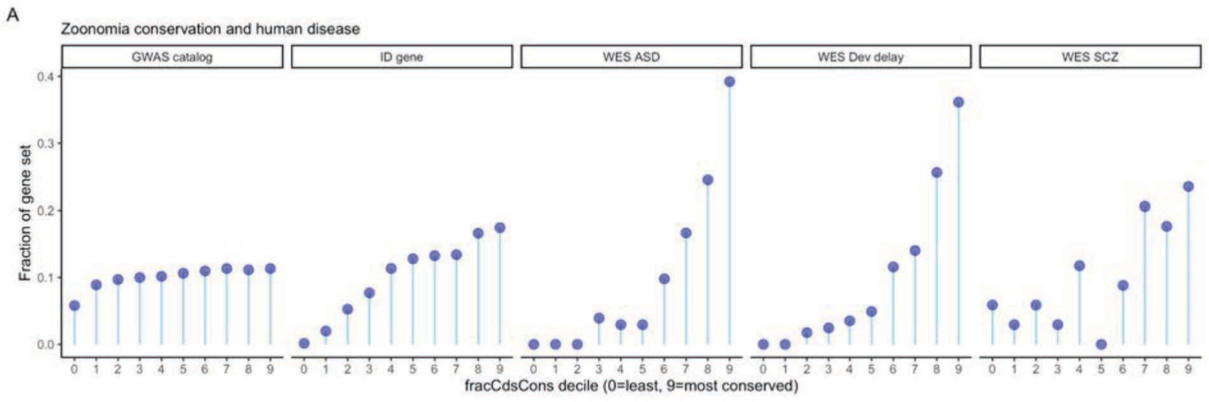

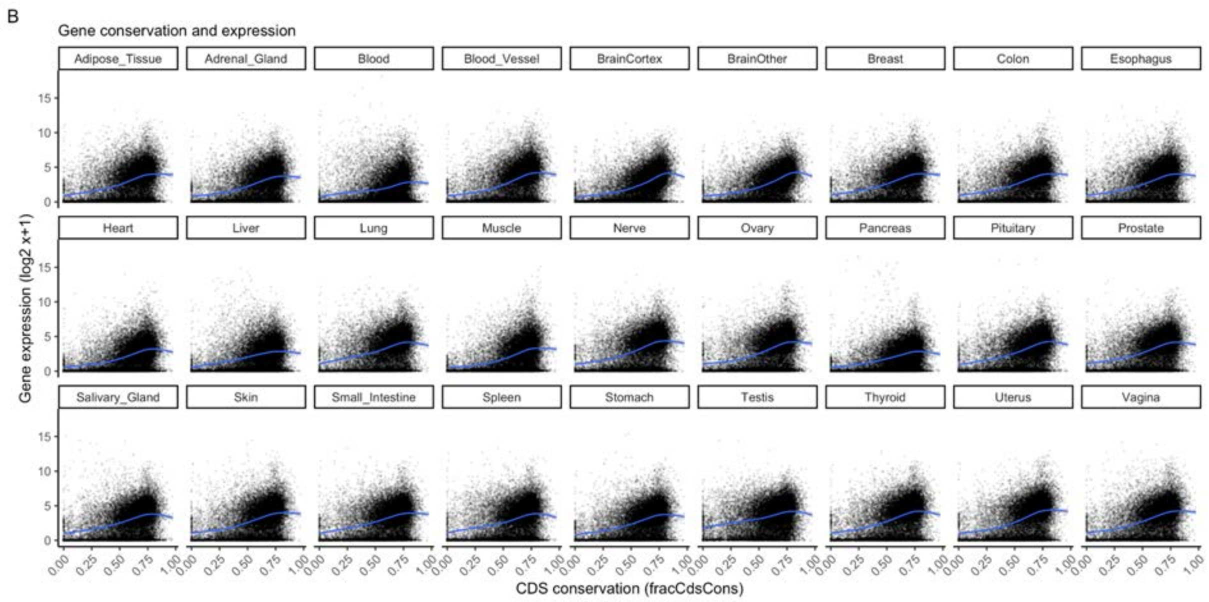

*Fig. S10.3. (A) fracCdsCons deciles versus external gene sets: genes with a genome-wide significant association in the EBI/NHGRI GWAS catalog, genes implicated in intellectual disability (ID), and genes implicated by whole exome association (WES) studies of autism spectrum disorder (ASD), developmental delay (dev deley), and schizophrenia (SCZ). Although there is some variation, genes with greater mammalian constraint tend to have greater human disease relevance. (B) 27 scatterplots, one for each GTEx tissue, where X-axis=fracCdsCons (0=least and 1=most constrained) and Y-axis=mean gene expression with -$log_2$ transformation. Each point is a gene. The blue line is a loess smoother. Although the Y-axis-intercepts and slopes of the blue lines vary, greater constraint is positively correlated with gene expression in all tissues.*

We evaluated the impact of common SNPs linked to PC genes in each fracCdsCons decile by using S-LDSC on 63 independent GWAS datasets (*Supplemental Methods, Section 6*). In the main analyses, SNPs were linked to genes using a functional approach (i.e., considering SNPs in exons, promoters, linked enhancers, or are fine-mapped cis-eQTLs using the cS2G approach (*65*)). We also performed secondary analyses by using a naive approach linking SNPs falling in a 100kb window around a gene. Analyses included the baseline-LD v.2.2 model, one annotation for all genes, 10 annotations for the deciles of six gene scores (fracCdsCons, pLI, LOEUF, and fraction of promoter, proximal enhancer and distal enhancer that were linked to the genes, see below). For each decile annotation, we meta-analyzed its heritability enrichment (i.e. fraction of heritability linked to the genes in the annotation divided by the fraction of common SNPs linked to the genes in the annotation) across independent GWAS datasets. We also computed its gene heritability enrichment, defined as the meta-analyzed heritability enrichment of the annotation divided by the mean of the meta-analyzed heritability enrichments of the 10 decile annotations (considered as a constant to compute standard errors). Significance of gene heritability enrichments were computed using z scores. We observed significantly higher gene $h^2$ enrichment for SNPs linked to genes in the most constrained deciles (*Fig. S10.4* and *table S17*). Interestingly, while SNPs linked to genes had a higher heritability enrichment in a meta-analysis of 11 blood and immune traits vs 9 brain disorders, we observed stronger gene $h^2$ enrichment patterns for brain disorders compared to blood and immune traits (*Fig. S10.4*), suggesting that fracCdsCons can be used to prioritize genes in brain disorders. We obtained similar conclusions using both SNP-gene linking strategies (*Fig. S10.4*). We also observed similar patterns of gene $h^2$ enrichment between fracCdsCons, pLI and LOEUF (*Fig. S10.5*).

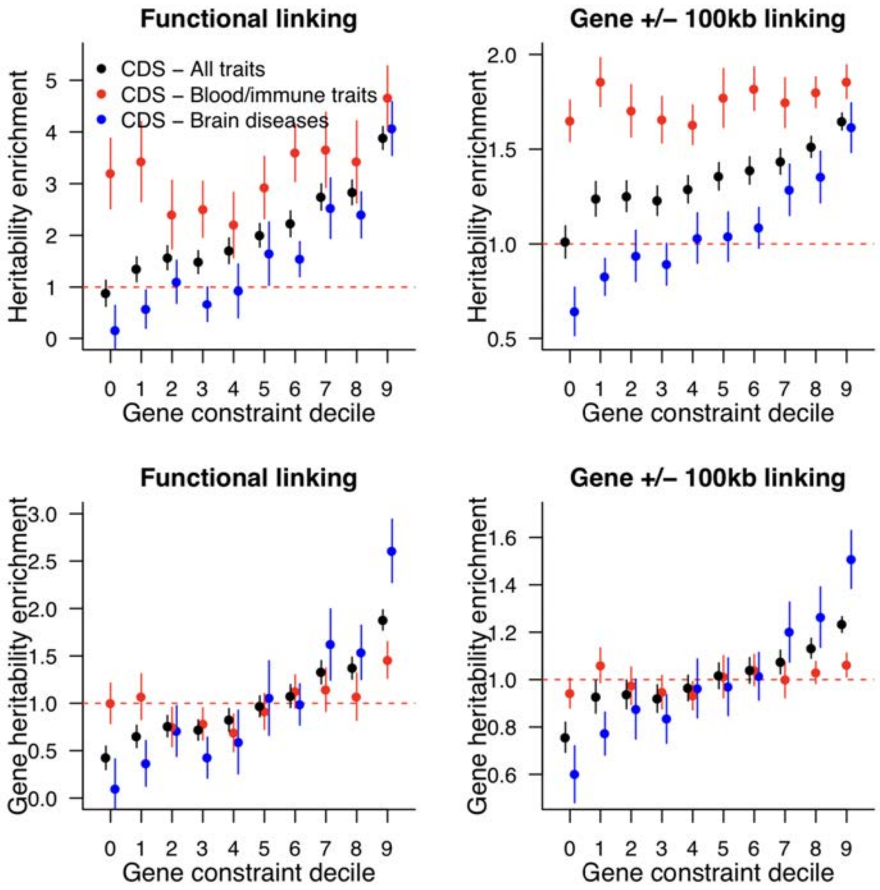

*Fig. S10.4. Heritability results SNPs linked to genes of each fracCdsCons deciles. We report heritability enrichment and gene heritability enrichment by using two approaches to link SNPs to genes. In each scenario we meta-analyzed values on 63 independent GWAS (black), 11 blood and immune traits, and 9 brain disorders. Dashed red lines represent a null enrichment of 1. Error bars are 95% confidence intervals.*

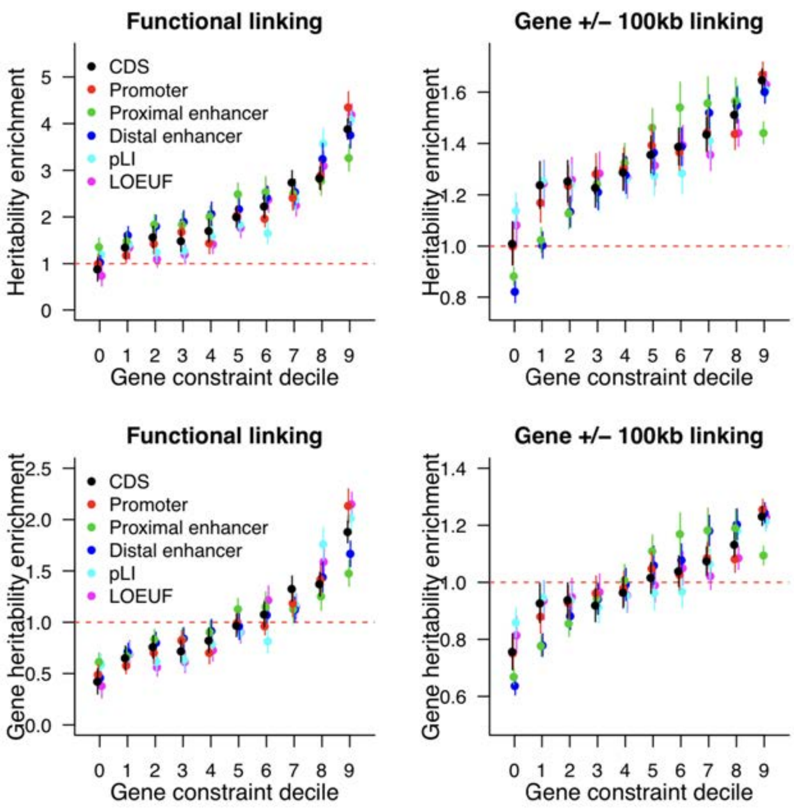

*Fig. S10.5. Heritability results for SNPs linked to genes of each decile of 6 gene constraint metrics. We report heritability enrichment and gene heritability enrichment by using two approaches to link SNPs to genes. In each scenario we meta-analyzed values on 63 independent GWAS (black). Dashed red lines represent a null enrichment of 1. Error bars are 95% confidence intervals.*

**Mammalian constraint compared to LOEUF:** We specifically compared fracCdsCons to the gnomAD LOEUF measure (*8*) given its widespread use. The Spearman correlation of fracCdsCons with LOEUF was -0.55, and LOEUF had the undesirable property of a strong residual correlation with the number of coding bases (*Supplemental Methods, Section 9*).

*Fig. S10.1D* shows the dispersed location of genes with missing gnomAD LOEUF scores (gene constraint based on human exonic variation) (*8*). Many PC genes do not have LOEUF scores (N=1,962 genes, 10.1%). *Fig. S10.6* shows an upset plot to contrast fracCdsCons clusters with LOEUF scores. In general, fracCdsCons appears to provide measures of constraint to more PC genes than LOEUF (98.6% vs 89.1%) and is thus applicable to a greater number of PC genes.

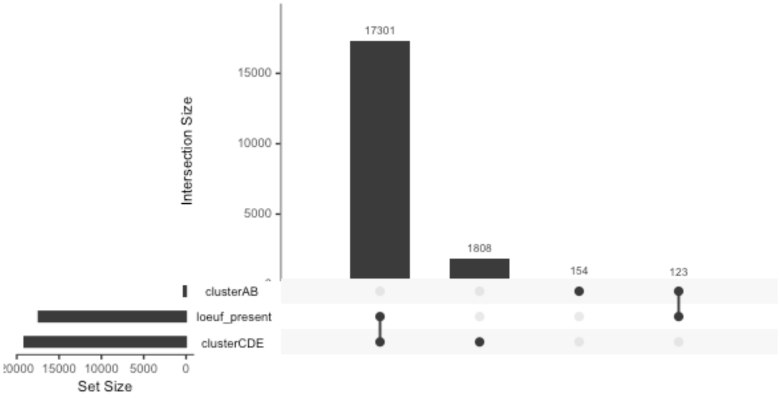

*Fig. S10.6. Upset plot for fracCdsCons clustering and LOEUF. The outlier clusters A and B were combined (clusterAB) and fracCdsCons is invalid. Clusters with valid fracCdsCons were combined (clusterCDE). LOEUF_present indicates whether LOEUF was non-missing for a gene. In summary: 89.2% of PC genes had valid fracCdsCons and LOEUF scores, 9.3% of genes had valid fracCdsCons but missing LOEUF, 0.79% of genes had missing or invalid scores for both, and 0.63% of genes had a LOEUF score alone (and none of these genes were in the most constrained quintile).*

*Fig. S10.7* shows a direct comparison of fracCdsCons with gnomAD LOEUF and pLI. The first two measures are smoothly continuous with no point of rarity at which to separate "constrained" from "not constrained". pLI is nearly bimodal.

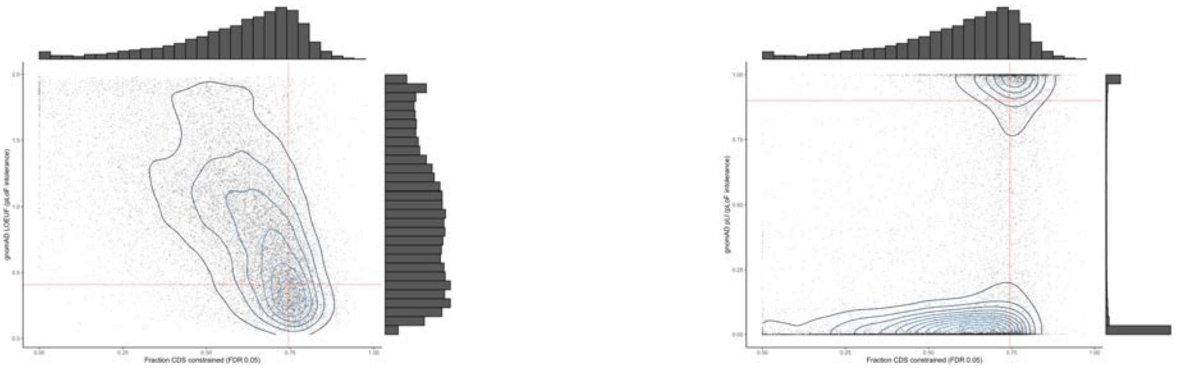

*Fig. S10.7. (A) Left, scatter plot of X-axis=fracCdsCons (fraction of constrained coding bases over mammalian evolution, larger values more constrained) and Y-axis=gnomAD LOEUF (an index of tolerance of pLoF, smaller values are more constrained). Each point is a gene with valid scores (N=17,301). The vertical red line is the 80th percentile for fracConsFdr05 (more constrained to the right), and the horizontal red line is the 20th percentile of LOEUF (more constrained lower). The contours indicate gene density, and the histograms on right and top show the univariate distributions. (B) Right, scatter plot of X-axis=fracCdsCons and Y=gnomAD pLI (an alternative index of tolerance of pLoF, higher values are more constrained), and most genes are near 0 or 1.*

These results are an important validator for both Zoonomia and gnomAD. *Fig. S10.7A* compares gene constraint between Zoonomia fracCdsCons and LOEUF. There was agreement between the gene measures (Spearman r = -0.56). This degree of congruence is notable as these measures are based on markedly divergent samples and phenomena—i.e., constraint over ~100 million years of mammalian evolution vs statistical modeling of pLoF counts in WES catalogs of humans who survived to consent to be included in a gnomAD cohort. In theory, these measures should be related and this empirical confirmation is an important validator for both measures.

However, the correlation is far from perfect, and we highlight that there are advantages and disadvantages to each measure: (a) missingness is lesser for fracCdsCons (see *Fig. S10.6*) and (b) LOEUF should perform better for human- or primate-specific genes as Zoonomia has low power (but, in practice, few of these genes had low LOEUF scores).

We investigated discrepancies between the most constrained quantile on one measure and the least constrained on the other. Genes with high fracCdsCons/low LOEUF constraint compared to high fracConsCds/high LOEUF constraint (i.e., top right vs bottom right corners of *Fig. S10.7A*) had smaller CDS sizes (mean 542 vs 2073 bases) as suggested by the large residual correlation of LOEUF with CDS size. We suspect that this resulted from an issue with the LOEUF model for smaller genes with far fewer expected pLoF variants (83 vs 390) and a large confidence interval (mean 1.3 vs 0.2) and thus high LOEUF score. We suspect that these genes are misclassified by LOEUF. For example, these genes included 3 small HOX genes that are known to be highly constrained (*HOXA9*, *HOXC12*, and *HOXC5*), and gene set analyses were significantly enriched for genes with specific expression in brain cortex, a generally highly constrained subset.

**Constraint between the parts of PC genes:** We evaluated the relationship between constraint of different gene parts in more detail here. *Fig. S10.8* depicts the relationship between the fraction of proximal promoters (50 bases upstream of TSS) under mammalian constraint with the fraction of coding bases under constraint. See *Supplementary Methods, Section 9* for derivation of this measure and comparisons with other gene constraint metrics. There is a strong positive correlation (Spearman rho=0.55) between constraint of CDS (fracCdsCons) and proximal promoter sequences, particularly for genes with any degree of constraint. We observed consistent correlation with constraint in 5'UTR and 3'UTR regions (rho = 0.62 and 0.60, respectively), and (at a lower extent) with constraint in introns (rho = 0.45).

We finally evaluated constraint in gene enhancers. Investigating gene enhancers is challenging as enhancers do not necessarily regulate the closest genes (*66*), and because ENCODE3 did not provide enhancer-gene linking predictions of their enhancer-like signatures (ELS) elements. Here, we used the EpiMap enhancer-gene linking predictions (*67*) to link ENCODE3 proximal and distal ELS (pELS and dELS, respectively). Briefly, EpiMap predicted enhancers in more than 800 cell-types (including ENCODE datasets), and linked these enhancers to their target genes using correlation between enhancer activity and gene expression across cell-types. Using EpiMap enhancer-gene links in all cell-types, we were able to link 33% and 53% of pELS and dELS to at least one gene, respectively. We observed non negligible correlation between dELS constraint gene scores with constraint in CDS and promoters (rho = 0.25 and 0.25, respectively); we note that we observed similar values when considering all EpiMap enhancers (rather than restricting to ENCODE3 cCRE), and lower correlation when considering pELS (*Fig. S10.8*). These data are consistent with the theory that if the function of a gene in mammals requires high constraint of protein structure, then the non-coding sequence in the proximal promoter that regulates its expression also tends to be constrained. As discussed below, for many genes these correlations reflect dosage sensitivity (as assessed by intolerance to predicted loss-of-function, pLoF, variation in large surveys of currently living humans) – if a mammalian gene cannot tolerate pLoF then this may imply a corresponding constraint of its regulatory mechanisms. Gene constraints are available in *table S18*. We note that 489 genes do not have 5'UTR constraint scores, 1,115 do not have Intron constraint scores, 2,293 do not have pELS constraint score, 1,469 do not have dELS constraint score, and 1,367 do not have EpiMap constraint score.

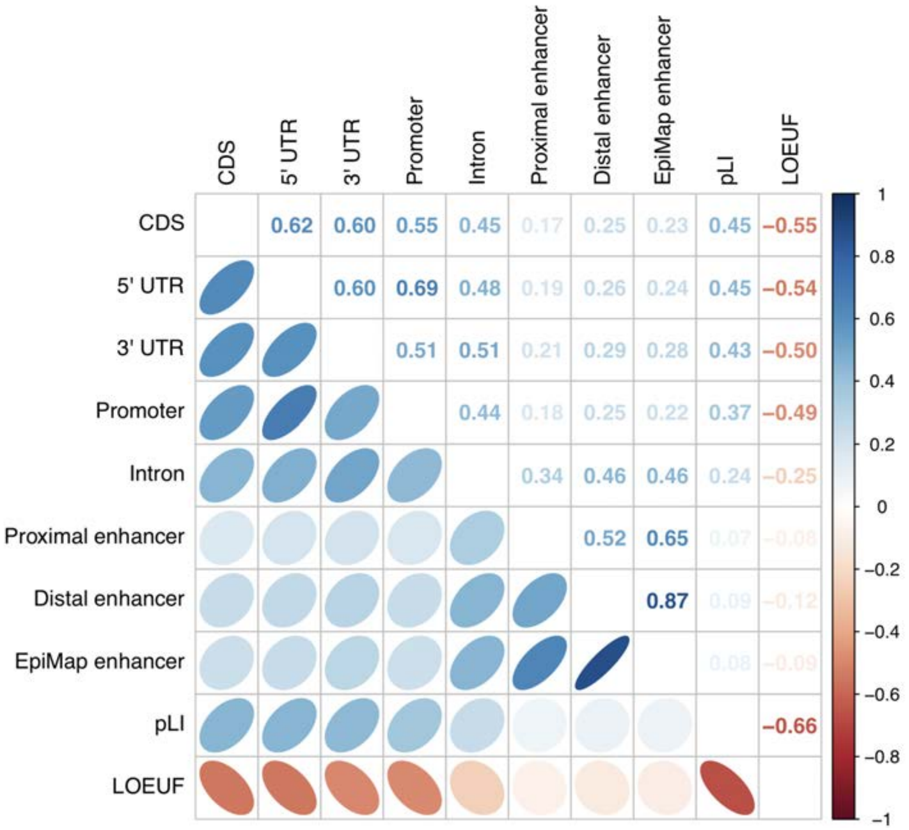

*Fig. S10.8. Spearman correlation of constraint fraction between the parts of PC genes.*

Finally, we evaluated the impact of common SNPs linked to genes in each constraint score decile by using S-LDSC on 63 independent GWAS datasets (see above). We observed that common SNPs linked to genes with constrained promoters and distal enhancers are as enriched in $h^2$ as genes with constrained CDS, suggesting that constraint in regulatory elements can be leveraged in the analyses of human diseases and complex traits (*Fig. S10.5*). We obtained similar conclusions using two SNP-gene linking strategies.

***Section 11: Rare copy number variation and human disease***

By: Jin Szatkiewicz, Jia Wen, Patrick Sullivan

Copy number variants (CNVs) are genomic segments that have fewer or more copies compared to a reference sample. Their sizes can range from 1kb to megabases. CNVs are important drivers of evolution and are notable risk factors for multiple human diseases (*68*–*70*). However, CNVs often occur in high repeat/low mappability regions meaning that detecting their presence and disease significance often carries considerable uncertainty (*71*, *72*). We thus evaluated the salience of mammalian constraint in the analysis of disease-related CNVs.

First, as an initial qualitative check, CNV deletions in a gene desert upstream of *SOX9* can cause the craniofacial condition Pierre Robin sequence by altering the copy number of distal enhancers. Functional analyses of the CNV-deletion regions in Pierre Robin sequence patients suggested a causal role for several small enhancers >1.2 Mb from *SOX9* (*73*). We found that these enhancers were highly constrained: the fraction of constrained bases comprising three small (3.5-6.9 kb) enhancers (1.45, 1.35, and 1.25 Mb from *SOX9*) were all 11-12%, and a functionally important element within one enhancer (the "min2" element (*73*)) was 73% constrained.

Second, we generalized the approach by analyzing TOPMed (*6*) structural variation (SV) calls from WGS. The *inline table* below presents descriptive data. Deletions were the most prevalent and smallest overall. All types of SV were rare, had moderate overlap with open chromatin regions, but had low overlap with CDS and constrained bases were underrepresented compared to the genome (~3.5%).

| *Variable* | *Deletion* | *Duplication* | *Inversion* |
|---|---|---|---|
| Number | 191,846 | 118,033 | 25,359 |
| Size (bp) | 6600 (3837-14888) | 16000 (5561-53900) | 10593 (4721-68332) |
| Allele frequency | 0.000007 (0.000004-0.000018) | 0.000004 (0.000004-0.000018) | 0.000004 (0.000004-0.000014) |
| High mappability | 0.770 (0.635-0.859) | 0.746 (0.612-0.834) | 0.758 (0.629-0.849) |
| ENCODE3 DHS | 0.138 (0.073-0.235) | 0.146 (0.076-0.238) | 0.151 (0.084-0.241) |
| CDS | 0 (0-0) | 0 (0-0.008) | 0 (0-0.007) |
| Mammal constrained | 0.016 (0.007-0.034) | 0.020 (0.009-0.040) | 0.021 (0.009-0.040) |
| Primate constrained | 0.012 (0.003-0.031) | 0.016 (0.004-0.039) | 0.017 (0.004-0.038) |

*TOPMed v5b structural variants (SV), autosomes, filtered for size ≥ 1kb, and call rate > 0.9. The columns show deletions, duplications, and inversions with respect to the reference genome. Values shown are median and interquartile range (p25-p75). Mappability is the fraction of SV bases with unique mapping (> 0.9 in 24 bp windows). ENCODE3 DHS is the fraction of SV bases that overlap DNase hypersensitive sites in any cell or tissue. CDS is the fraction of SV bases that overlaps a coding base (all protein-coding genes, pickAll). Mammal constrained is the fraction of SV bases that are constrained in all mammals (FDR 0.05) and primate constrained is the fraction of SV bases that are constrained in primates.*

However, these SV calls are highly overlapping, and the 335,238 SVs occur in 2,182 genomic regions. A given base can be part of multiple SV that range from very rare to common. To obtain a more detailed view of the impact of SV with respect to base pair constraint, we converted the TOPMed SV calls to base pair annotations. For each base, we recorded the total allele count of all SV intersecting that base (separately by SV type), the phyloP score at that base, and whether it was a CDS base. Over all bases impacted by an SV, the median (IQR) allele counts per base were: deletions 3 (1-9), duplications 7 (2-32), and inversions 5 (2-19).

We then looked at the effects of allele count per base and SV prevalence, constraint, and CDS (*Fig. S11.1*). For SV deletions, duplications, and inversions, the fraction of bases, the fraction constrained, and the fraction intersecting a CDS base were the most prevalent for singleton SV. These fractions decreased with allele counts in the rare range. The patterns differed markedly for bases that were intersected by SVs with a high total allele count; in the common range, deletions had the lowest values whereas duplications and inversion were markedly higher. These results are consistent with deletions being, on average, more deleterious due to exposure of dosage sensitive elements via haploinsufficiency. Duplications and inversions can be disruptive (e.g., via triplosensitivity or disruption of gene or regulatory structure) but may otherwise be relatively tolerated.

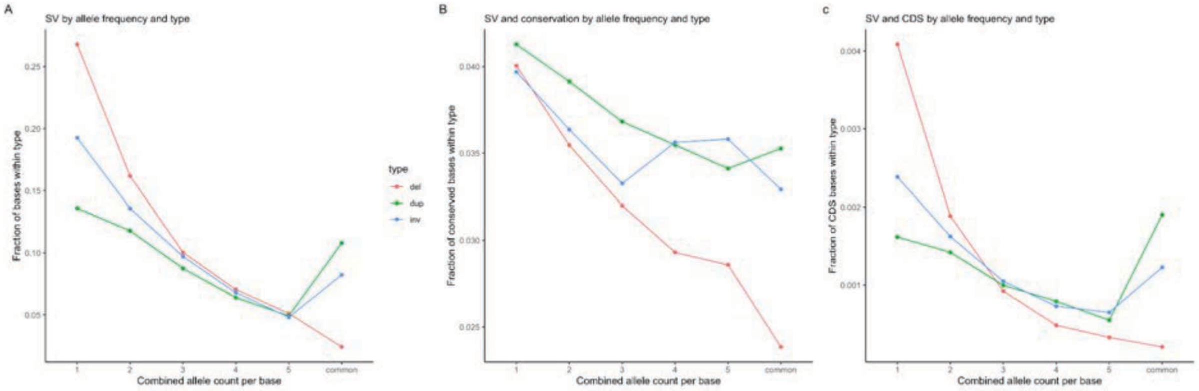

*Fig. S11.1. TOPMed SV converted to the combined allele count per base (X-axis). A value of 1 means that a base was intersected by a single SV with an allele count of 1 in TOPMed. A value of 2 means that a base was intersected by two different SVs each with allele count of 1 or by one SV with an allele count of 2. There is a break between combined allele count of 5 and "common" (frequency ≥ 0.005). Panel A shows the fraction of bases within each SV type (deletion, duplication, or inversion). Singletons are the most frequent within SV type, and decrease with allele count. For the common category, deletions are the lowest but duplication and inversions increase. Panel B shows the fraction of constrained bases (Zoonomia phyloP ≥ 2.270) by allele count and by type. Panel C shows the fraction of CDS bases by allele count and by type. All panels show a qualitatively different relation with allele count by type.*

Finally, we evaluated whether Zoonomia constraint measures were useful in CNV analyses for a complex trait. Schizophrenia has a notable CNV contribution (*74*), and we obtained the CNV call set from the largest published study (*75*) that applied multiple calling algorithms and careful quality control and harmonization procedures to array intensity data from 21,094 cases with schizophrenia and 20,227 controls.

We first replicated the main findings from the paper for increased CNV "burden" in cases vs controls. We then re-annotated all CNVs by including the number of constrained bases encompassed by each CNV, reasoning that this may better capture the potential effects of a CNV on both regulatory and CDS in case-control burden comparisons. As shown in the *inline table* (controlling for five ancestry principal components and genotyping array), we found that the numbers of constrained bases were 4-5 logs more significant as a predictor of case-control differences compared to the traditional measures of CNV length and number of CNVs. Assessing CNV burden of bases constrained over mammalian evolution may better capture the impact of changes in copy number by identifying important genic and non-genetic elements.

| *CNV type* | *CNV burden measure* | *Beta* | *SE* | *Statistic* | *P-value* |
|---|---|---|---|---|---|
| All | Constrained bases | 7.93e-6 | 7.72e-7 | 10.27 | 1.0e-24 |
| | Length | 1.75e-7 | 2.20e-8 | 7.98 | 1.4e-15 |
| | Number of CNVs | 2.87e-2 | 6.64e-3 | 4.33 | 1.5e-5 |
| Deletion | Constrained bases | 1.23e-5 | 1.37e-6 | 9.01 | 2.0e-19 |
| | Length | 3.03e-7 | 3.98e-8 | 7.63 | 2.4e-14 |
| | Number of CNVs | 2.88e-2 | 9.60e-3 | 3.00 | 2.7e-3 |
| Duplication | Constrained bases | 5.90e-6 | 9.98e-7 | 5.91 | 3.4e-9 |
| | Length | 1.16e-7 | 2.65e-8 | 4.39 | 1.2e-5 |
| | Number of CNVs | 2.96e-2 | 9.34e-3 | 3.17 | 1.5e-3 |

***Section 12: Somatic Cancer Driver Genes and Constraint***

By: Sharadha Sakthikumar, Jessika Nordin, Ananya Roy, Kerstin Lindblad-Toh, Karin Forsberg Nilsson

The full set of processes used to identify and functionally explore non-coding constrained driver genes of cancers can be found in *Companion paper #12, Sakthikumar et al*. Key methodologies are included in brief below.

**Medulloblastoma and pilocytic astrocytoma sample set and basic validation.** We downloaded variant calling format files with somatic mutations based on whole genome sequencing data from the International Cancer Genomics Consortium (ICGC, project code PBCA-DE). The data were aligned to the human genome build hg19. A total of 89 pilocytic astrocytoma and 146 medulloblastoma samples were available. A standard analysis to identify significantly mutated genes was performed to validate our data against previous analyses. The GATK's Funcotator module (*76*) was used to functionally classify the medulloblastoma somatic point mutation (SPM) and somatic indel mutations (SIM) as being either coding or non-coding changes. Coding mutations were used for identification of SMGs. We found one SMG in pilocytic astrocytoma and three SMGs in medulloblastoma, consistent with what has been seen previously.

**Identification of non-coding constraint mutations (NCCMs).** SPM and SIM calls were annotated with mammalian phyloP scores to identify NCCMs. PhyloP scores based on hg38 were converted to hg19 coordinates using liftOver (*77*). Previous estimates of the fraction of the human genome that is functional has been between 6-13%. We used an estimate of the top 8% constraint positions (i.e., mammalian phyloP $\geq 1.2$) which we previously used to examine NCCMs in glioblastoma (*78*). Non-coding variants with phyloP $\geq 1.2$ were termed NCCMs. NCCMs located within 5′ or 3′ UTRs, introns, or ±100 kb intergenic flanking regions of protein-coding genes were extracted. The rate of NCCMs, normalized to the length of the queried genomic region for each gene was computed, and those genes with $\geq 2$ NCCMs per 100 kb were termed as candidate driver genes.

**Evaluation of putative functional impact on NCCMs.** To verify if the NCCMs around candidate genes have cis-regulatory changes, their coordinates were annotated with regulatory annotations from UCSC (*77*) and ENCODE. Annotations included transcription factor binding sites (TFBS), regulatory markers (ORegAnno), transcription start sites (TSS), enhancers, and DNase I hypersensitive sites (DHS). sTRAP (*79*) was used to predict if the NCCMs were likely to alter the binding affinity of transcription factors to the mutated versus wild-type sequences. For every NCCM allele and its corresponding wild-type, 20 bp upstream and downstream were used as input sequences. The matrices for the analysis were set to the JASPAR vertebrates database (*79*, *80*), and for the background model, the 'human promoter' option was selected. The JASPAR database (http://jaspar.genereg.net) was used to obtain information about TF binding matrices and motifs of interest. Affinity profiles for the top-ranking matrices with $P \leq 0.05$ were recorded as having significantly more TF binding for the input sequences than what could be expected from a random sequence.

**Electromobility shift assays (EMSA) of NCCMs.** Multiple EMSAs were run for NCCMs in the MB002 medulloblastoma cell line (*81*). All probes and experimental conditions are stated in the *Companion paper #12, Sakthikumar et al*.

**Expression and survival analysis.** To assess gene expression levels between normal cerebellum and medulloblastoma, or between the medulloblastoma subgroups, processed gene expression data from Gene Expression Omnibus (GEO) were used. Dataset GSE124814 contains batch-normalized transcriptional profiles of 1350 medulloblastoma samples and 291 normal cerebellum samples (*82*). Dataset GSE85217 contains 763 medulloblastoma samples that were used for survival analysis (*83*). Based on the order of expression, every increasing expression value is used as a cutoff to create the two groups and the significance difference tested in a log-rank test. The most significant expression cutoff for survival analysis is plotted on a Kaplan Meier curve.

### Supplementary Figures

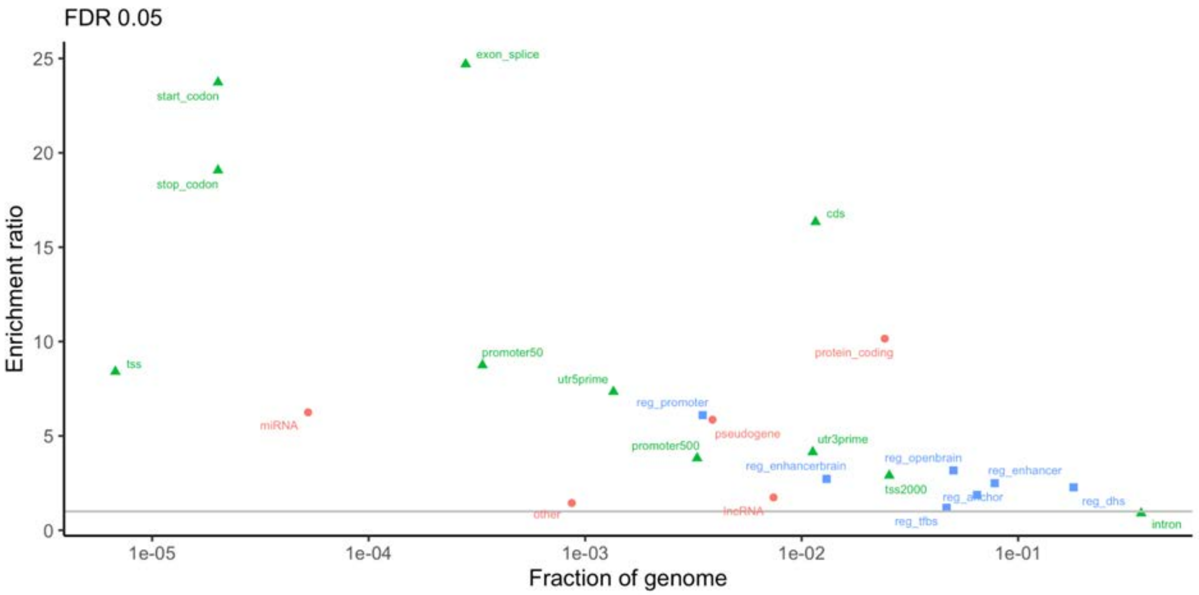

***Fig. S1. Evolutionary constraint in multiple genomic partitions.***

*Enrichment ratios for genomic features. Enrichment ratios are the fraction of a feature that is constrained (5% FDR mammalian phyloP), divided by the fraction of the genome that is constrained. For example, 0.577 of CDS bases are significantly constrained (=19,069,108/33,073,903) and 0.0352 of the genome is significantly constrained (=100,651,377/2,852,623,265) for an enrichment ratio of 16.3 (=0.577/0.352). The horizontal grey line has an enrichment ratio of 1. Genomic features are defined above. Orange=cDNA exons; green=parts of PC genes (pickOne set); and blue=regulatory annotations.*

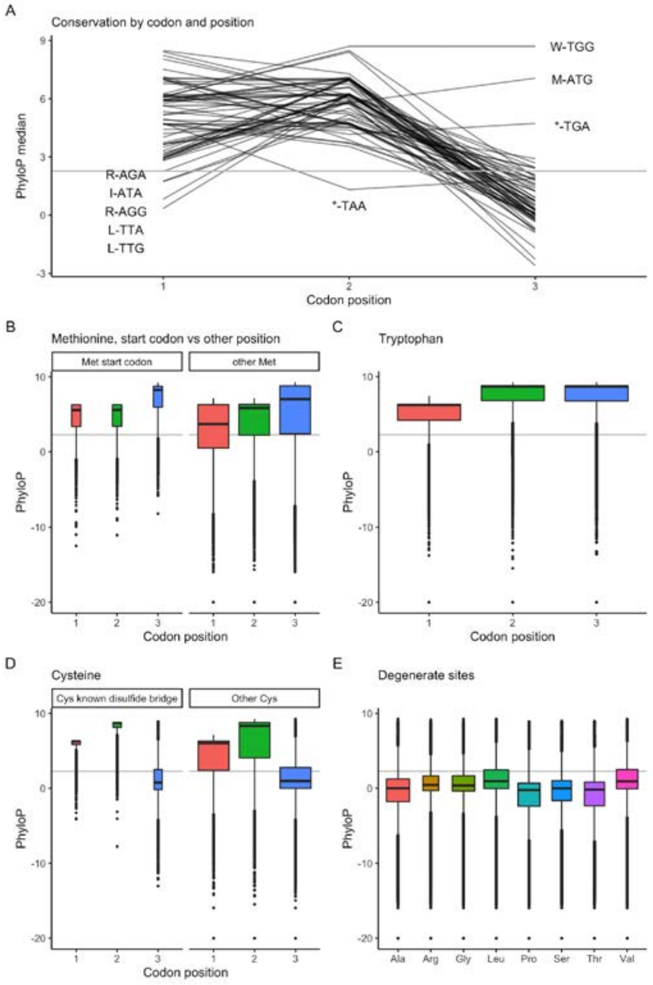

***Fig. S2. Constraint scores captured nuances of the genetic code.***

*(A) Parallel plot of median phyloP scores for all PC CDS by codon and codon position (pickOne subset, 32,127,234 bases and 10,709,078 codons). X-axis=codon position. Y-axis=median phyloP score (grey line at 2.270 is the FDR 0.05 genome-wide cutoff). The plot has 64 lines, one per codon. The median phyloP scores for each position were 4.93, 5.98, and 0.68. However, there were exceptions to these general trends as labeled on the graph. For example, position 3 for tryptophan (W-TGG, median 8.70) and methionine (M-ATG, median 7.06) are highly constrained–these are the two amino acids with a single codon and any third position change is non-synonymous. I=Ile, L=Leu, M=Met, R=Arg, W=Trp, *=Ter. (B) Median phyloP values for Met in the start position (8.7%) versus elsewhere in the CDS (91.3%). As Met has one codon (ATG), any mutation is non-synonymous and is an impactful start-lost if in the first codon. Met is highly constrained but particularly in the start position (median phyloP per position of 5.6, 5.6, and 8.3) compared to Met elsewhere in a protein (median phyloP 3.7, 5.8, and 7.0). The IQR is far smaller in start codon Met (IQR around 2.8) compared to Met in other positions (IQR of 4.1 or greater). (C) Tryptophan (Trp) is the other single codon amino acid. All positions are highly constrained but with variability. Of all transcripts, 93.1% contain a Trp and, for most of these transcripts, all positions in all Trp are highly constrained; for ~25% of transcripts containing ≥ 10 Trp, not all Trp positions were constrained (e.g., 96/98 Trp in MUC16 and 87/95 Trp in DMBT1 were not constrained at all positions). (D) Evaluating cysteine (Cys) residues that are or are not part of a known intra-peptide disulfide bridge (6.85% of 756,942 Cys). Non-synonymous mutations in a Cys can ablate a disulfide bridge and lead to loss-of-function due to disruption in peptide folding. Cys in known disulfide bridges had higher median phyloP values and narrower IQR. (E) Half of all codons are degenerate as positions 1 and 2 determine the amino acid and position 3 can be any base. The box plots show the distribution of phyloP scores in the degenerate third positions of Ala, Arg, Gly, Leu, Pro, Ser, Thr, and Val. The medians show low levels of constraint (Pro -0.22 to Val 0.93). However, there is variability with 18.6% of 5,263,653 degenerate sites being highly constrained.*

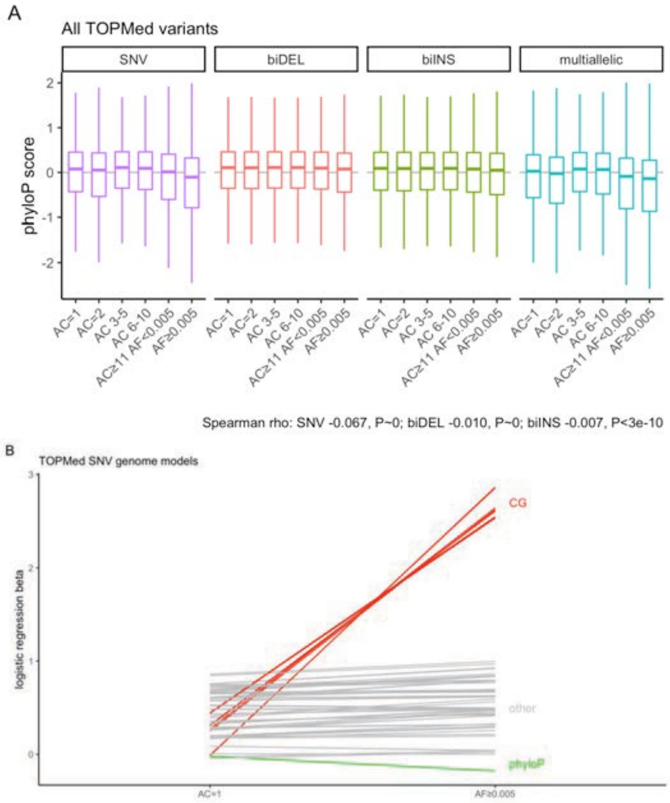

***Fig. S3. Plots of the relation between AC and phyloP score for all TOPMed variants by frequency and type.***

*Variants were classified in 5 AC bins of decreasing rarity plus common variation (AF ≥ 0.005). We included all ancestries as appropriate for mammalian comparisons. (A) Description of genome-wide constraint, variant type, and variant frequency. The boxplot thick line is the median, the rectangle shows 25th–75th percentiles, and whiskers extend 1.5 times the IQR above and below the box. Violin plots were impractical due to the number of variants. AC=allele count, AF=allele frequency. There is a small negative correlation between AC-phyloP for SNVs and near zero correlations for non-SNV types of variation. (B) Parallel plot of modeling results for singleton (AC=1) and common SNVs (AF≥0.005). Because nearly all P-values were smaller than the operating system's minimum numeric value (P < 2.2e-308), the Y-axis shows patterns of beta coefficients between the two logistic regression models. Each model had 64 terms (phyloP score plus 63 base sequences AAC to TTT compared to reference AAA). Green line shows betas for phyloP constraint. Most of the 3 base context terms were similar (grey lines) with the exception of the 8 terms containing the dinucleotide CG (red, GCN or NGC). The modelling was done in this way (i.e., singleton SNVs vs 10x random local positions and common SVS vs 10x random local positions) due to the great imbalance in the numbers of variants: of all 552 million SNVs, 45.6% AC=1 and 2.6% AF ≥ 0.005.*

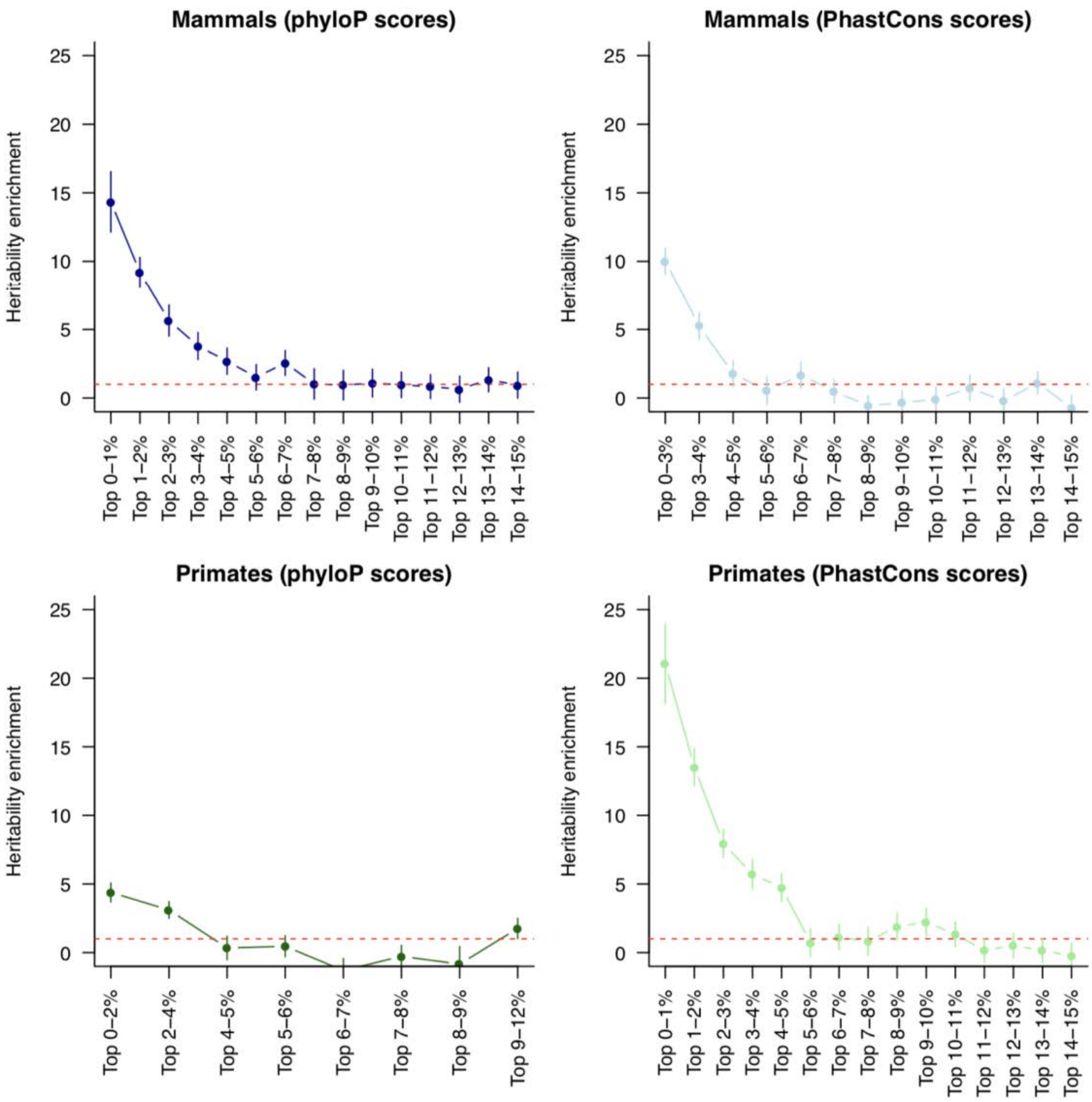

***Fig. S4. Heritability enrichment of SNPs falling in the top percentiles of constraint scores.***

*We report heritability enrichment for constraint annotations based on PhyloP and PhastCons scores computed in either Mammals or Primates. Some annotations contain multiple percentiles due to a large number of bases with similar scores (for example the top 0-3% PhastCons scores in Mammals annotation contain SNPs with a maximum PhastCons score of 1). Results are meta-analyzed across 63 independent GWAS. Error bars represent 95% confidence intervals. Numerical results are reported in table S2.*

A

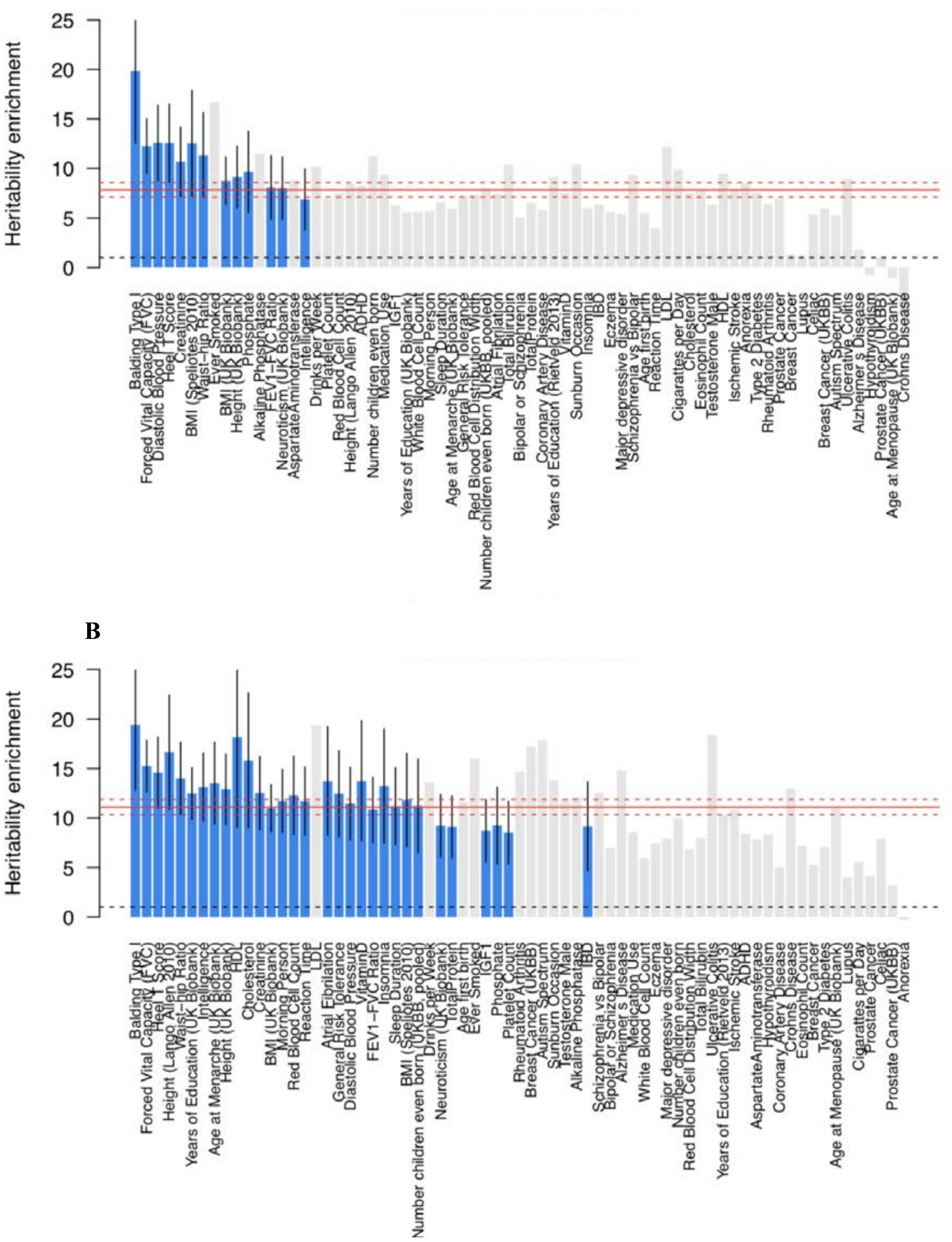

***Fig. S5. Heritability enrichment of the constrained in mammals (A) and constrained in primates (B) annotations in 69 GWAS.***

*Traits are sorted by the lower bound of the 95% confidence interval. Significant enrichment (P < 0.05/69) are plotted in blue, with error bars representing 95% confidence intervals. Red lines represent enrichment and corresponding 95% confidence intervals meta-analyzed across 63 independent GWAS. Numerical results are reported in table S5.*

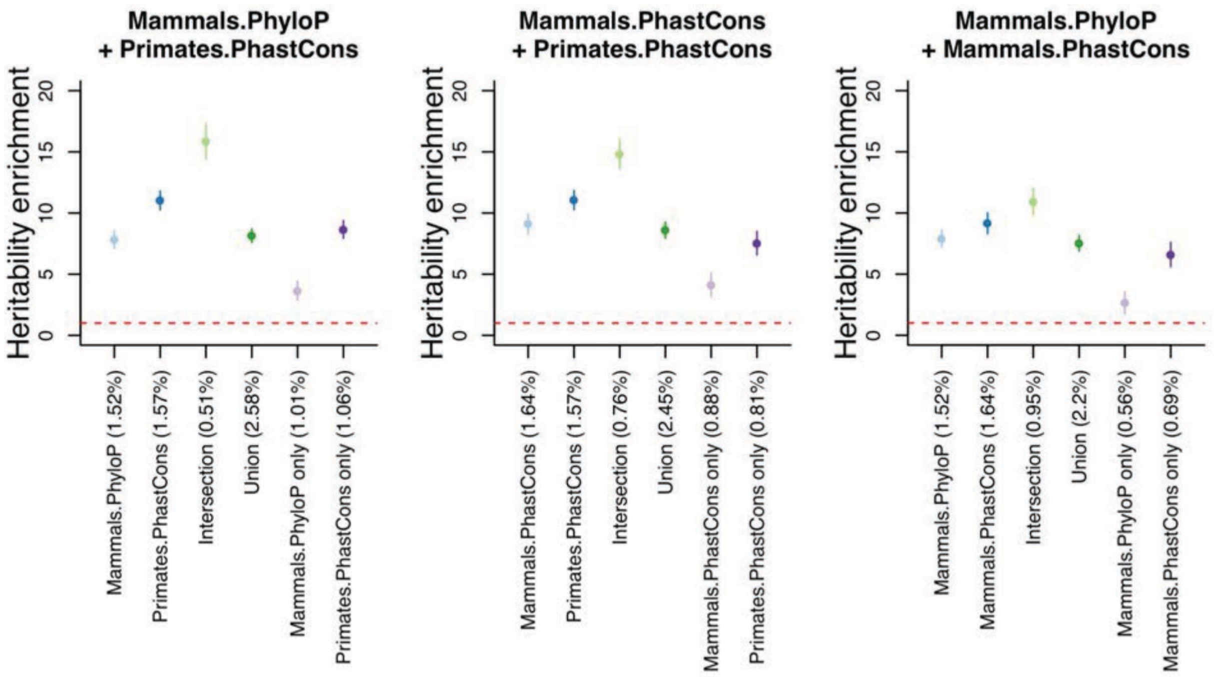

***Fig. S6. Heritability enrichment of pairs of constrained annotations intersected together.***

*We report heritability enrichment of pairs of constrained annotations intersected together. Error bars represent 95% confidence interval. Each constrained annotation represents 3.54% of the genome bases with the highest score (consistent with the FDR 5% threshold measured in phyloP scores in mammals). We observed that annotations derived from phyloP scores in mammals and PhastCons scores in primates (as in main analyses) produce very large heritability enrichment (left panel), that using phastCons scores in mammals rather than phyloP scores in mammals produce similar conclusions (middle panel), and that annotations derived from phyloP scores in mammals and PhastCons scores in mammals do not have significantly different heritability enrichments, but that intersecting the two produces higher enrichment than the annotation derived from phyloP scores in mammals (right panel).*

A

B

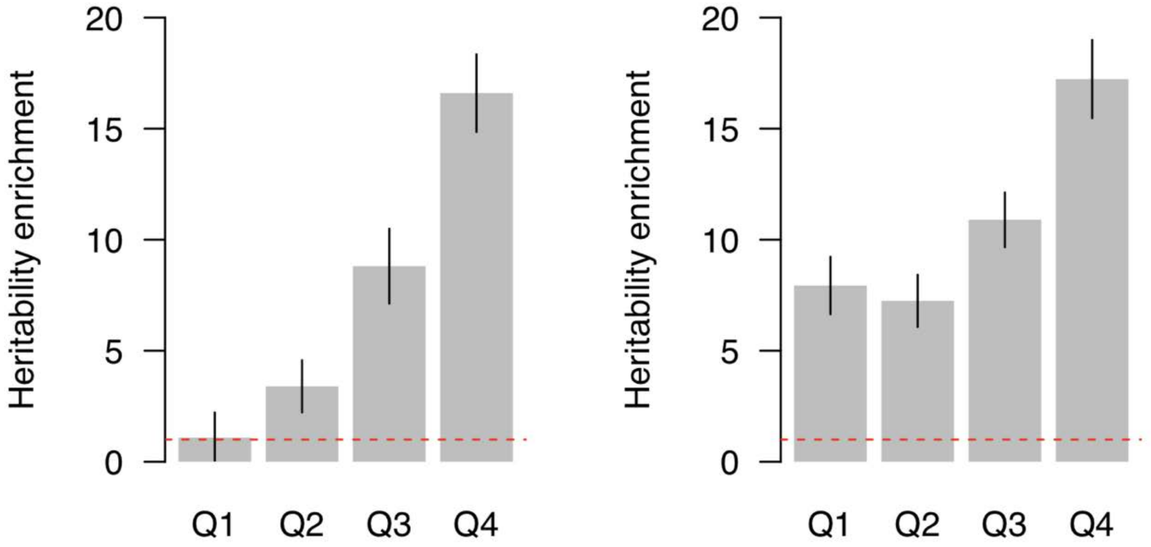

***Fig. S7. Heritability enrichment of the constraint in mammal annotation stratified by quartiles of PhastCons scores in primates (A) and of the constraint in primate annotation stratified by quartiles of phyloP scores in mammals (B).***

*PhastCons score quartile boundaries in regions constraint in mammals are 0.000, 0.038, 0.625, 0.991, and 1.000. PhyloP score quartile boundaries in regions constraint in primates are -20.000, 0.219, 1.170, 2.903, and 8.903. Error bars represent 95% confidence intervals.*

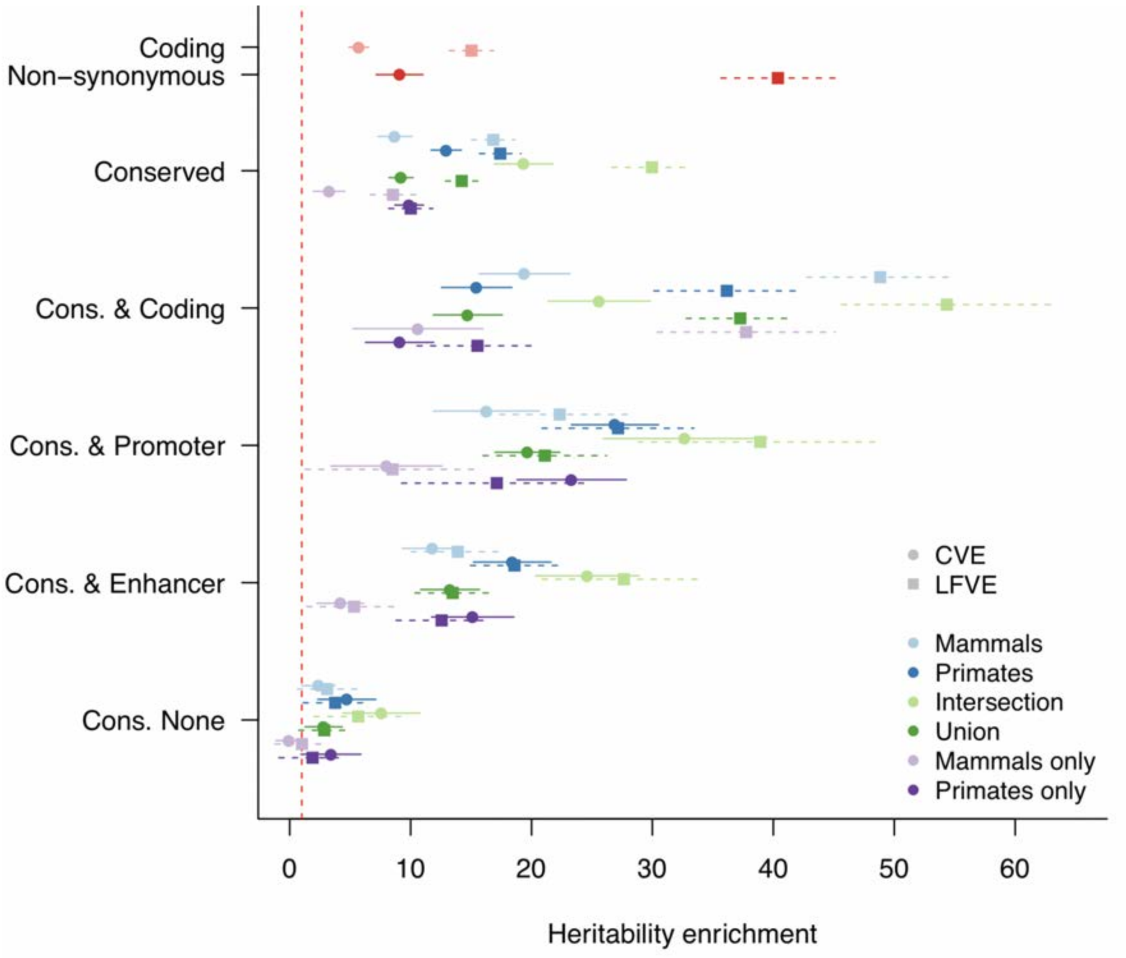

***Fig. S8. Common and low-frequency variant heritability enrichments in constraint regions.***

*We report common variant heritability enrichment (CVE) and low-frequency variant heritability enrichment (LFVE) of constrained in mammals and primates annotations intersected together, and stratified by their genomic function. We plotted coding and non-synonymous enrichments for comparison purposes. The dashed red lines represent a null enrichment of 1. Results are meta-analyzed across 27 independent UK Biobank traits. Error bars represent 95% confidence intervals. Numerical results are reported in table S10.*

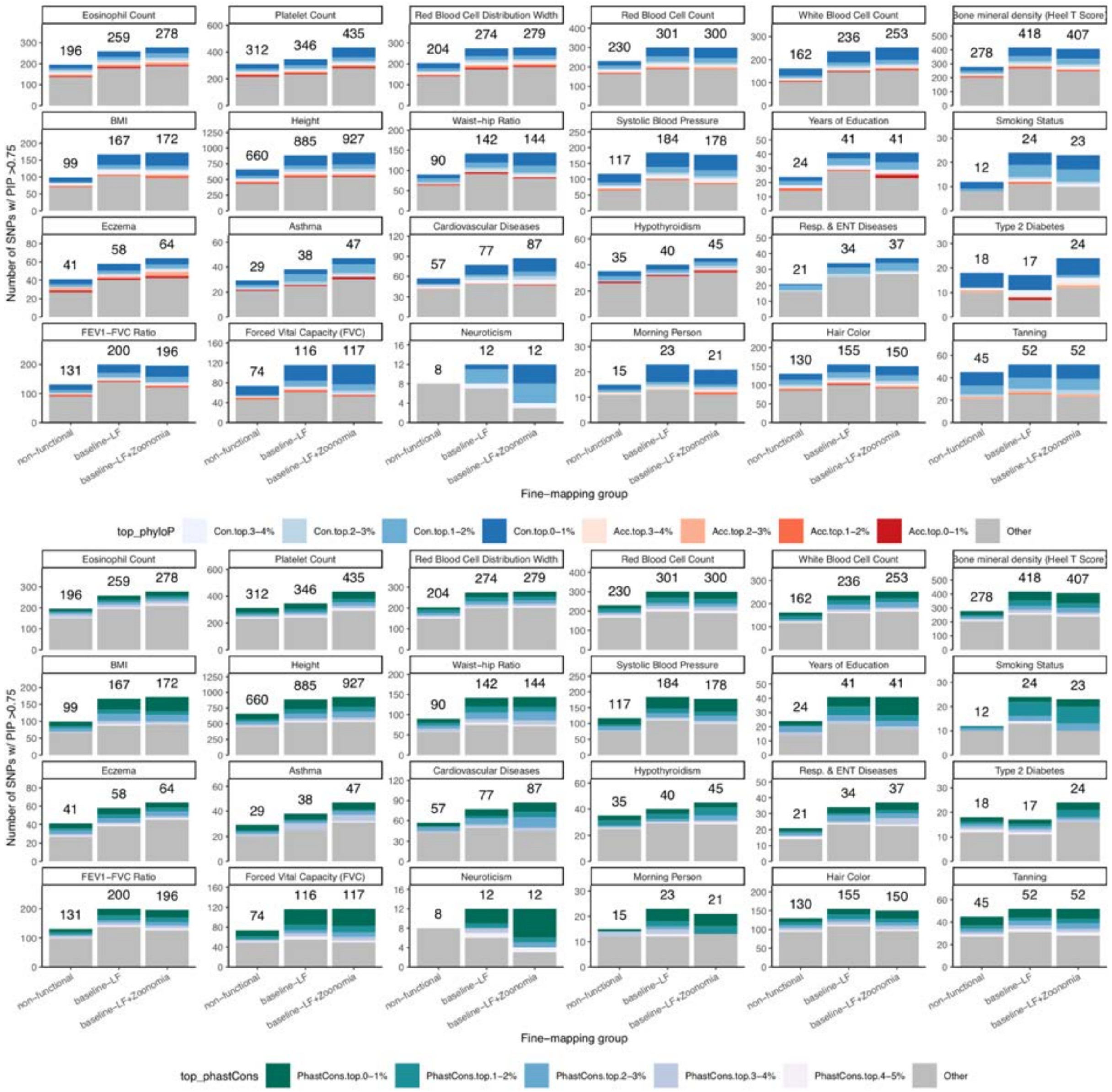

Fig. S9. Functional fine-mapping UKBB GWAS SNPs with Zoonomia constraint scores.

*We report the number of fine-mapped SNPs across UKBB traits that have posterior inclusion probability (PIP) greater than 0.75 using no functional annotation, the baseline-LF model, and the baseline-LF+Zoonomia model. We highlight SNPs that fall into various non-overlapping phyloP annotations (top conserved annotations: blue, top accelerating annotations: red, not in phyloP annotation: gray) (top) and SNPs that fall into various non-overlapping PhastCons annotations (top conserved annotations: colored, not in PhastCons annotation: gray) (bottom).*

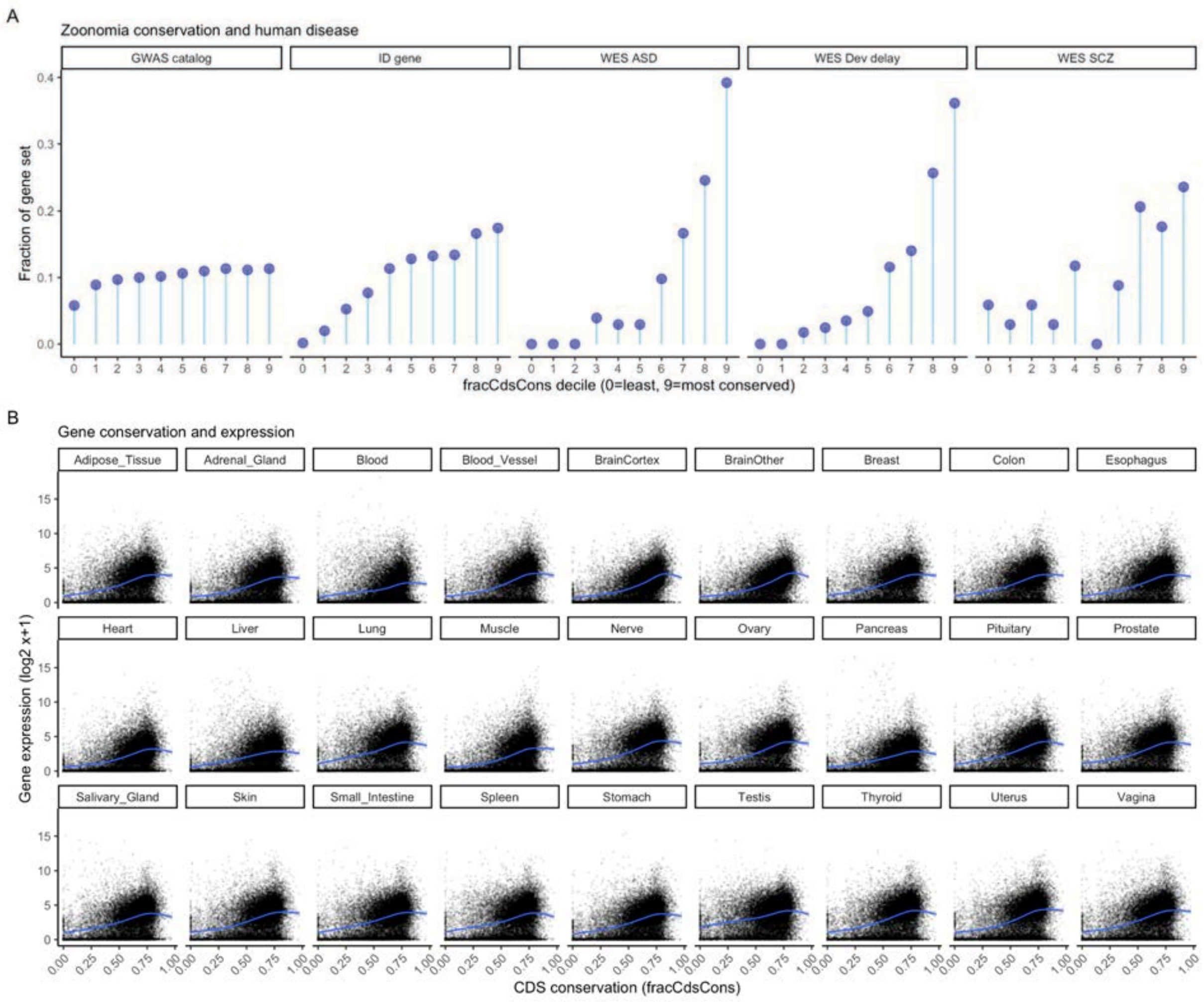

***Fig. S10. Constraint and human disease and gene expression.***

*(A) fracCdsCons deciles versus external gene sets: genes with a genome-wide significant association in the EBI/NHGRI GWAS catalog, genes implicated in intellectual disability (ID), and genes implicated by whole exome association (WES) studies of autism spectrum disorder (ASD), developmental delay, and schizophrenia (SCZ). Although there is some variation, genes with greater mammalian constraint tend to have greater human disease relevance. (B) 27 scatterplots, one for each GTEx tissue, where X=fracCdsCons (0=least and 1=most constrained) and Y=mean gene expression with -$log_2$ transformation. Each point is a gene. The blue line is a loess smoother. Although the Y-intercepts and slopes of the blue lines vary, greater constraint is positively correlated with gene expression in all tissues.*

### Supplementary Tables

***Table S1. GWAS summary statistics used in common variant heritability analyses.***

We report the 69 summary statistics used in this study, including 63 independent GWAS covering 57 independent diseases and complex traits, 11 autoimmune diseases and blood cell traits, and 9 bran diseases.

***Table S2. Heritability partitioning results for common SNPs falling in the top percentiles of phyloP and PhastCons scores in mammals and primates.***

We report the heritability enrichments (with corresponding standard errors and P-values), and S-LDSC regression coefficients (with corresponding standard errors and P-values) for common SNPs falling in the top percentiles of phyloP and PhastCons scores in mammals and primates. Results are meta-analyzed across 63 independent GWAS.

***Table S3. Heritability partitioning results for common SNPs as a function of their distance to a constrained base.***

We report the heritability enrichments (with corresponding standard errors and P-values), for common SNPs as a function of their distance to a constrained base. Results are meta-analyzed across 63 independent GWAS.

***Table S4. Heritability partitioning results for the main S-LDSC analysis.***

We report the heritability enrichments (with corresponding standard errors and P-values), and S-LDSC regression coefficients (with corresponding standard errors and P-values) for our main model including 97 annotations from the baseline-LD model v2.2, 6 ENCODE3 annotations, and 3 zoonomia annotations. Results are meta-analyzed across 63 independent GWAS.

***Table S5. Heritability partitioning results for the constrained in mammals and constrained in primates annotations in 69 GWAS.***

We report the heritability enrichments (with corresponding standard errors and P-values) for the constrained in mammals and constrained in primates annotations in 69 GWAS, as well as the statistical significance between the trait enrichment and the meta-analyzed enrichment.

***Table S6. Heritability partitioning results for constrained annotations in 11 blood and immune traits and 9 brain disorders.***

We report the heritability enrichments (with corresponding standard errors and P-values) for constrained in mammals and primates annotations intersected together and meta-analyzed across 11 blood and immune traits and 9 brain disorders.

***Table S7. Heritability partitioning results for constrained in mammals and primates annotations intersected together and stratified by their genomic function.***

We report the heritability enrichments (with corresponding standard errors and P-values), and S-LDSC regression coefficients (with corresponding standard errors and P-values) for constrained in mammals and primates annotations intersected together and stratified by their genomic function. Results are meta-analyzed across 63 independent GWAS.

***Table S8. S-LDXR results for the 28 main binary annotations of the baseline-LD-X model and 3 constrained annotations.***

We report the enrichment of stratified squared trans-ancestry genetic correlation (with corresponding standard errors and P-values) for the 28 main binary annotations of the baseline-LD-X model and 3 constrained annotations. Results are meta-analyzed across 31 independent traits.

***Table S9. UK Biobank GWAS summary statistics used in common and low-frequency variant heritability analyses.***

We report the 27 summary statistics used to partition common and low-frequency variant heritability. We report their sample size (N), effective sample size (Neff), prevalence (for binary traits), and common and low-frequency variant heritability (with corresponding Z score) estimated using the baseline-LF model. We restricted our analyses to these traits (different the the 63 used for common variant heritability analyses), as low-frequency variant heritability analyses require very large sample size (*48*). These traits are the same as the ones used in ref. (*48*).

***Table S10. Common and low-frequency variant heritability partitioning results for constrained in mammals and primates annotations intersected together and stratified by their genomic function.***

We report the common and low-frequency variant heritability enrichments (with corresponding standard errors and P-values for differences) for constrained in mammals and primates annotations intersected together and stratified by their genomic function. Results are meta-analyzed across 27 independent UK Biobank traits.

***Table S11. Standardized squared effect sizes as a function of allele frequency.***

We report the standardized squared effect sizes in multiple annotations where variants are stratified by their allele frequency. Results are meta-analyzed across 27 independent UK Biobank traits.

***Table S12. Functionally-informed fine-mapping results across 24 UK Biobank traits.***

We report functionally-informed fine-mapping results using 3 sets of annotations for 24 UK Biobank traits. We report trait sample size (N) and effective sample size (Neff), the number of fine-mapped variants with posterior inclusion probability (PIP) > 0.75 and the number and fraction of fine-mapped variants with PIP > 0.75 that are constrained in mammals or primates.

***Table S13. Fine-mapped variants within UNICORNs.***

We report the location and posterior inclusion probability (PIP) of fine-mapped variants within UNannotated Intergenic COnstraint RegioNs (UNICORNs) as well as the trait associated with each variant. Further, we indicate the ranking of SNPs within the primate PhastCons and mammalian phyloP score distributions.

***Table S14. Gene constraint metrics for 19,386 protein-coding genes.***

We report gene constraint metrics (pickOne transcript subset) referred to in the manuscript including the fraction of CDS bases constrained in mammals and in primates. We also include the UMAP/DBSCAN cluster identifiers (A-E), the constraint ventiles, and indicate the genes in the top and bottom ventiles (5%) .

***Table S15. Gene Set Analysis: most/least constrained 5% of genes (GO gene sets).***

We report the results of the hypergeometric gene set analyses of Gene Ontology (GO) pathways for genes in the top or bottom ventiles of mammalian constraint. The background gene set was 19,109 protein-coding genes. We provide data used for P-value calculation.

***Table S16. Gene Set Analysis: most/least constrained 5% of genes (SynGO terms).***

We report the results of the hypergeometric gene set analyses of the expert-curated synaptic gene sets from the SynGO consortium. The background gene set was 19,109 protein-coding genes. We provide data used for P-value calculation.

***Table S17. Heritability partitioning results for common variants linked to constrained gene sets.***

We report the heritability enrichments (with corresponding standard errors and P-values) and gene heritability enrichment (with corresponding standard errors and P-values) for common variants linked to gene sets based on their decile for their constrained gene scores. We considered 6 different constrained scores (fraction of CDS bases constrained, fraction of promoter bases constrained, fraction of ENCODE3 proximal enhancer bases constrained, fraction of ENCODE3 distal enhancer bases constrained, pLI, and LOEUF). We considered two strategies to link SNPs to genes: by using the cS2G map (*65*), and by using the gene-body +/- 100 kb window.

***Table S18. Mammalian gene constraint across different parts of PC genes.***

For each protein coding genes, we report the constraint fraction of their CDS (fracCdsCons), 5' UTR, 3'UTR, promoter, intron, proximal enhancers (pELS), distal enhancers (dELS) and EpiMap enhancers.

***Table S19. ClinVar classes of variants identified as constrained with TOGA.***

Gene alignments generated using TOGA were explored to identify constrained amino acid residues unchanged across mammalian evolutionary history. All sites were compared with known pathogenic/benign records in ClinVar. The number of sites, as well as their ClinVar disease status where present, are displayed.

***Table S20. Naturally occurring adaptive 'pathogenic' variants in non-human mammals.***

Pathogenic residues identified in ClinVar were explored in non-human mammals. Any instances of potentially adaptive 'pathogenic' human variants occurring naturally in other species are documented. All species showing human pathogenic residues are listed.

6. D. Taliun, D. N. Harris, M. D. Kessler, J. Carlson, Z. A. Szpiech, R. Torres, S. A. G. Taliun, A. Corvelo, S. M. Gogarten, H. M. Kang, A. N. Pitsillides, J. LeFaive, S.-B. Lee, X. Tian, B. L. Browning, S. Das, A.-K. Emde, W. E. Clarke, D. P. Loesch, A. C. Shetty, T. W. Blackwell, A. V. Smith, Q. Wong, X. Liu, M. P. Conomos, D. M. Bobo, F. Aguet, C. Albert, A. Alonso, K. G. Ardlie, D. E. Arking, S. Aslibekyan, P. L. Auer, J. Barnard, R. G. Barr, L. Barwick, L. C. Becker, R. L. Beer, E. J. Benjamin, L. F. Bielak, J. Blangero, M. Boehnke, D. W. Bowden, J. A. Brody, E. G. Burchard, B. E. Cade, J. F. Casella, B. Chalazan, D. I. Chasman, Y.-D. I. Chen, M. H. Cho, S. H. Choi, M. K. Chung, C. B. Clish, A. Correa, J. E. Curran, B. Custer, D. Darbar, M. Daya, M. de Andrade, D. L. DeMeo, S. K. Dutcher, P. T. Ellinor, L. S. Emery, C. Eng, D. Fatkin, T. Fingerlin, L. Forer, M. Fornage, N. Franceschini, C. Fuchsberger, S. M. Fullerton, S. Germer, M. T. Gladwin, D. J. Gottlieb, X. Guo, M. E. Hall, J. He, N. L. Heard-Costa, S. R. Heckbert, M. R. Irvin, J. M. Johnsen, A. D. Johnson, R. Kaplan, S. L. R. Kardia, T. Kelly, S. Kelly, E. E. Kenny, D. P. Kiel, R. Klemmer, B. A. Konkle, C. Kooperberg, A. Köttgen, L. A. Lange, J. Lasky-Su, D. Levy, X. Lin, K.-H. Lin, C. Liu, R. J. F. Loos, L. Garman, R. Gerszten, S. A. Lubitz, K. L. Lunetta, A. C. Y. Mak, A. Manichaikul, A. K. Manning, R. A. Mathias, D. D. McManus, S. T. McGarvey, J. B. Meigs, D. A. Meyers, J. L. Mikulla, M. A. Minear, B. D. Mitchell, S. Mohanty, M. E. Montasser, C. Montgomery, A. C. Morrison, J. M. Murabito, A. Natale, P. Natarajan, S. C. Nelson, K. E. North, J. R. O'Connell, N. D. Palmer, N. Pankratz, G. M. Peloso, P. A. Peyser, J. Pleiness, W. S. Post, B. M. Psaty, D. C. Rao, S. Redline, A. P. Reiner, D. Roden, J. I. Rotter, I. Ruczinski, C. Sarnowski, S. Schoenherr, D. A. Schwartz, J.-S. Seo, S. Seshadri, V. A. Sheehan, W. H. Sheu, M. B. Shoemaker, N. L. Smith, J. A. Smith, N. Sotoodehnia, A. M. Stilp, W. Tang, K. D. Taylor, M. Telen, T. A. Thornton, R. P. Tracy, D. J. Van Den Berg, R. S. Vasan, K. A. Viaud-Martinez, S. Vrieze, D. E. Weeks, B. S. Weir, S. T. Weiss, L.-C. Weng, C. J. Willer, Y. Zhang, X. Zhao, D. K. Arnett, A. E. Ashley-Koch, K. C. Barnes, E. Boerwinkle, S. Gabriel, R. Gibbs, K. M. Rice, S. S. Rich, E. K. Silverman, P. Qasba, W. Gan, NHLBI Trans-Omics for Precision Medicine (TOPMed) Consortium, G. J. Papanicolaou, D. A. Nickerson, S. R. Browning, M. C. Zody, S. Zöllner, J. G. Wilson, L. A. Cupples, C. C. Laurie, C. E. Jaquish, R. D. Hernandez, T. D. O'Connor, G. R. Abecasis, Sequencing of 53,831 diverse genomes from the NHLBI TOPMed Program. *Nature*. **590**, 290–299 (2021).

7. M. Lek, K. J. Karczewski, E. V. Minikel, K. E. Samocha, E. Banks, T. Fennell, A. H. O'Donnell-

Luria, J. S. Ware, A. J. Hill, B. B. Cummings, T. Tukiainen, D. P. Birnbaum, J. A. Kosmicki, L. E. Duncan, K. Estrada, F. Zhao, J. Zou, E. Pierce-Hoffman, J. Berghout, D. N. Cooper, N. Deflaux, M. DePristo, R. Do, J. Flannick, M. Fromer, L. Gauthier, J. Goldstein, N. Gupta, D. Howrigan, A. Kiezun, M. I. Kurki, A. L. Moonshine, P. Natarajan, L. Orozco, G. M. Peloso, R. Poplin, M. A. Rivas, V. Ruano-Rubio, S. A. Rose, D. M. Ruderfer, K. Shakir, P. D. Stenson, C. Stevens, B. P. Thomas, G. Tiao, M. T. Tusie-Luna, B. Weisburd, H.-H. Won, D. Yu, D. M. Altshuler, D. Ardissino, M. Boehnke, J. Danesh, S. Donnelly, R. Elosua, J. C. Florez, S. B. Gabriel, G. Getz, S. J. Glatt, C. M. Hultman, S. Kathiresan, M. Laakso, S. McCarroll, M. I. McCarthy, D. McGovern, R. McPherson, B. M. Neale, A. Palotie, S. M. Purcell, D. Saleheen, J. M. Scharf, P. Sklar, P. F. Sullivan, J. Tuomilehto, M. T. Tsuang, H. C. Watkins, J. G. Wilson, M. J. Daly, D. G. MacArthur, Exome Aggregation Consortium, Analysis of protein-coding genetic variation in 60,706 humans. *Nature*. **536**, 285–291 (2016).

8. K. J. Karczewski, L. C. Francioli, G. Tiao, B. B. Cummings, J. Alföldi, Q. Wang, R. L. Collins, K. M. Laricchia, A. Ganna, D. P. Birnbaum, L. D. Gauthier, H. Brand, M. Solomonson, N. A. Watts, D. Rhodes, M. Singer-Berk, E. M. England, E. G. Seaby, J. A. Kosmicki, R. K. Walters, K. Tashman, Y. Farjoun, E. Banks, T. Poterba, A. Wang, C. Seed, N. Whiffin, J. X. Chong, K. E. Samocha, E. Pierce-Hoffman, Z. Zappala, A. H. O'Donnell-Luria, E. V. Minikel, B. Weisburd, M. Lek, J. S. Ware, C. Vittal, I. M. Armean, L. Bergelson, K. Cibulskis, K. M. Connolly, M. Covarrubias, S. Donnelly, S. Ferriera, S. Gabriel, J. Gentry, N. Gupta, T. Jeandet, D. Kaplan, C. Llanwarne, R. Munshi, S. Novod, N. Petrillo, D. Roazen, V. Ruano-Rubio, A. Saltzman, M. Schleicher, J. Soto, K. Tibbetts, C. Tolonen, G. Wade, M. E. Talkowski, Genome Aggregation Database Consortium, B. M. Neale, M. J. Daly, D. G. MacArthur, The mutational constraint spectrum quantified from variation in 141,456 humans. *Nature*. **581**, 434–443 (2020).

14. D. Earl, N. Nguyen, G. Hickey, R. S. Harris, S. Fitzgerald, K. Beal, I. Seledtsov, V. Molodtsov, B. J. Raney, H. Clawson, J. Kim, C. Kemena, J.-M. Chang, I. Erb, A. Poliakov, M. Hou, J. Herrero, W. J.

Kent, V. Solovyev, A. E. Darling, J. Ma, C. Notredame, M. Brudno, I. Dubchak, D. Haussler, B. Paten, Alignathon: a competitive assessment of whole-genome alignment methods. *Genome Res.* **24**, 2077–2089 (2014).

22. ENCODE Project Consortium, J. E. Moore, M. J. Purcaro, H. E. Pratt, C. B. Epstein, N. Shoresh, J. Adrian, T. Kawli, C. A. Davis, A. Dobin, R. Kaul, J. Halow, E. L. Van Nostrand, P. Freese, D. U. Gorkin, Y. Shen, Y. He, M. Mackiewicz, F. Pauli-Behn, B. A. Williams, A. Mortazavi, C. A. Keller, X.-O. Zhang, S. I. Elhajjajy, J. Huey, D. E. Dickel, V. Snetkova, X. Wei, X. Wang, J. C. Rivera-Mulia, J. Rozowsky, J. Zhang, S. B. Chhetri, J. Zhang, A. Victorsen, K. P. White, A. Visel, G. W. Yeo, C. B. Burge, E. Lécuyer, D. M. Gilbert, J. Dekker, J. Rinn, E. M. Mendenhall, J. R. Ecker, M. Kellis, R. J. Klein, W. S. Noble, A. Kundaje, R. Guigó, P. J. Farnham, J. M. Cherry, R. M. Myers, B. Ren, B. R. Graveley, M. B. Gerstein, L. A. Pennacchio, M. P. Snyder, B. E. Bernstein, B. Wold, R. C. Hardison, T. R. Gingeras, J. A. Stamatoyannopoulos, Z. Weng, Expanded encyclopaedias of DNA elements in the human and mouse genomes. *Nature*. **583**, 699–710 (2020).

31. D. Wang, S. Liu, J. Warrell, H. Won, X. Shi, F. C. P. Navarro, D. Clarke, M. Gu, P. Emani, Y. T. Yang, M. Xu, M. J. Gandal, S. Lou, J. Zhang, J. J. Park, C. Yan, S. K. Rhie, K. Manakongtreecheep, H. Zhou, A. Nathan, M. Peters, E. Mattei, D. Fitzgerald, T. Brunetti, J. Moore, Y. Jiang, K. Girdhar, G. E. Hoffman, S. Kalayci, Z. H. Gümüş, G. E. Crawford, PsychENCODE Consortium, P. Roussos, S. Akbarian, A. E. Jaffe, K. P. White, Z. Weng, N. Sestan, D. H. Geschwind, J. A. Knowles, M. B. Gerstein, Comprehensive functional genomic resource and integrative model for the human brain. *Science*. **362** (2018), doi:10.1126/science.aat8464.

37. P. Akbari, A. Gilani, O. Sosina, J. A. Kosmicki, L. Khrimian, Y.-Y. Fang, T. Persaud, V. Garcia, D. Sun, A. Li, J. Mbatchou, A. E. Locke, C. Benner, N. Verweij, N. Lin, S. Hossain, K. Agostinucci, J. V. Pascale, E. Dirice, M. Dunn, Regeneron Genetics Center, DiscovEHR Collaboration, W. E. Kraus, S. H. Shah, Y.-D. I. Chen, J. I. Rotter, D. J. Rader, O. Melander, C. D. Still, T. Mirshahi, D. J. Carey, J. Berumen-Campos, P. Kuri-Morales, J. Alegre-Díaz, J. M. Torres, J. R. Emberson, R. Collins, S. Balasubramanian, A. Hawes, M. Jones, B. Zambrowicz, A. J. Murphy, C. Paulding, G. Coppola, J. D. Overton, J. G. Reid, A. R. Shuldiner, M. Cantor, H. M. Kang, G. R. Abecasis, K. Karalis, A. N. Economides, J. Marchini, G. D. Yancopoulos, M. W. Sleeman, J. Altarejos, G. Della Gatta, R. Tapia-Conyer, M. L. Schwartzman, A. Baras, M. A. R. Ferreira, L. A. Lotta, Sequencing of 640,000 exomes identifies variants associated with protection from obesity. *Science*. **373** (2021), doi:10.1126/science.abf8683.

46. B. K. Bulik-Sullivan, Schizophrenia Working Group of the Psychiatric Genomics Consortium, P.-R. Loh, H. K. Finucane, S. Ripke, J. Yang, N. Patterson, M. J. Daly, A. L. Price, B. M. Neale, LD Score regression distinguishes confounding from polygenicity in genome-wide association studies. *Nature*

*Genetics*. **47** (2015), pp. 291–295.

55. D. Griesemer, J. R. Xue, S. K. Reilly, J. C. Ulirsch, K. Kukreja, J. Davis, M. Kanai, D. K. Yang, S. B. Montgomery, C. D. Novina, R. Tewhey, P. C. Sabeti, Genome-Wide Functional Screen of 3'UTR Variants Uncovers Causal Variants for Human Disease and Evolution. *SSRN Electronic Journal*, , doi:10.2139/ssrn.3762769.

64. F. Koopmans, P. van Nierop, M. Andres-Alonso, A. Byrnes, T. Cijsouw, M. P. Coba, L. N. Cornelisse, R. J. Farrell, H. L. Goldschmidt, D. P. Howrigan, N. K. Hussain, C. Imig, A. P. H. de Jong, H. Jung, M. Kohansalnodehi, B. Kramarz, N. Lipstein, R. C. Lovering, H. MacGillavry, V. Mariano, H. Mi, M. Ninov, D. Osumi-Sutherland, R. Pielot, K.-H. Smalla, H. Tang, K. Tashman, R. F. G. Toonen, C. Verpelli, R. Reig-Viader, K. Watanabe, J. van Weering, T. Achsel, G. Ashrafi, N. Asi, T. C. Brown, P. De Camilli, M. Feuermann, R. E. Foulger, P. Gaudet, A. Joglekar, A. Kanellopoulos, R. Malenka, R. A. Nicoll, C. Pulido, J. de Juan-Sanz, M. Sheng, T. C. Südhof, H. U. Tilgner, C. Bagni, À. Bayés, T. Biederer, N. Brose, J. J. E. Chua, D. C. Dieterich, E. D. Gundelfinger, C. Hoogenraad, R. L. Huganir, R. Jahn, P. S. Kaeser, E. Kim, M. R. Kreutz, P. S. McPherson, B. M. Neale, V. O'Connor, D. Posthuma, T. A. Ryan, C. Sala, G. Feng, S. E. Hyman, P. D. Thomas, A. B. Smit, M. Verhage, SynGO: An Evidence-Based, Expert-Curated Knowledge Base for the Synapse. *Neuron*. **103**, 217–234.e4 (2019).

65. S. Gazal, O. Weissbrod, F. Hormozdiari, K. Dey, J. Nasser, K. Jagadeesh, D. Weiner, H. Shi, C. Fulco, L. O'Connor, B. Pasaniuc, J. M. Engreitz, A. L. Price, Combining SNP-to-gene linking strategies to pinpoint disease genes and assess disease omnigenicity. *medRxiv*, 2021.08.02.21261488 (2021).

66. A. Sanyal, B. R. Lajoie, G. Jain, J. Dekker, The long-range interaction landscape of gene promoters. *Nature*. **489**, 109–113 (2012).

67. C. A. Boix, B. T. James, Y. P. Park, W. Meuleman, M. Kellis, Regulatory genomic circuitry of human disease loci by integrative epigenomics. *Nature*. **590**, 300–307 (2021).

68. H. Innan, F. Kondrashov, The evolution of gene duplications: classifying and distinguishing between

models. *Nat. Rev. Genet.* **11**, 97–108 (2010).

75. C. R. Marshall, D. P. Howrigan, D. Merico, B. Thiruvahindrapuram, W. Wu, D. S. Greer, D. Antaki, A. Shetty, P. A. Holmans, D. Pinto, M. Gujral, W. M. Brandler, D. Malhotra, Z. Wang, K. V. F. Fajarado, M. S. Maile, S. Ripke, I. Agartz, M. Albus, M. Alexander, F. Amin, J. Atkins, S. A. Bacanu, R. A. Belliveau Jr, S. E. Bergen, M. Bertalan, E. Bevilacqua, T. B. Bigdeli, D. W. Black, R. Bruggeman, N. G. Buccola, R. L. Buckner, B. Bulik-Sullivan, W. Byerley, W. Cahn, G. Cai, M. J. Cairns, D. Campion, R. M. Cantor, V. J. Carr, N. Carrera, S. V. Catts, K. D. Chambert, W. Cheng, C. R. Cloninger, D. Cohen, P. Cormican, N. Craddock, B. Crespo-Facorro, J. J. Crowley, D. Curtis, M. Davidson, K. L. Davis, F. Degenhardt, J. Del Favero, L. E. DeLisi, D. Dikeos, T. Dinan, S. Djurovic, G. Donohoe, E. Drapeau, J. Duan, F. Dudbridge, P. Eichhammer, J. Eriksson, V. Escott-Price, L. Essioux, A. H. Fanous, K.-H. Farh, M. S. Farrell, J. Frank, L. Franke, R. Freedman, N. B. Freimer, J. I. Friedman, A. J. Forstner, M. Fromer, G. Genovese, L. Georgieva, E. S. Gershon, I. Giegling, P. Giusti-Rodríguez, S. Godard, J. I. Goldstein, J. Gratten, L. de Haan, M. L. Hamshere, M. Hansen, T. Hansen, V. Haroutunian, A. M. Hartmann, F. A. Henskens, S. Herms, J. N. Hirschhorn, P. Hoffmann, A. Hofman, H. Huang, M. Ikeda, I. Joa, A. K. Kähler, R. S. Kahn, L. Kalaydjieva, J. Karjalainen, D. Kavanagh, M. C. Keller, B. J. Kelly, J. L. Kennedy, Y. Kim, J. A. Knowles, B. Konte, C. Laurent, P. Lee, S. H. Lee, S. E. Legge, B. Lerer, D. L. Levy, K.-Y. Liang, J. Lieberman, J. Lönnqvist, C. M. Loughland, P. K. E. Magnusson, B. S. Maher, W. Maier, J. Mallet, M. Mattheisen, M. Mattingsdal, R. W. McCarley, C. McDonald, A. M. McIntosh, S. Meier, C. J. Meijer, I. Melle, R. I. Mesholam-Gately, A. Metspalu, P. T. Michie, L. Milani, V. Milanova, Y. Mokrab, D. W. Morris, B. Müller-Myhsok, K. C. Murphy, R. M. Murray, I. Myin-Germeys, I. Nenadic, D. A. Nertney, G. Nestadt, K. K. Nicodemus, L. Nisenbaum, A. Nordin, E. O'Callaghan, C. O'Dushlaine, S.-Y. Oh, A. Olincy, L. Olsen, F. A. O'Neill, J. Van Os, C. Pantelis, G. N. Papadimitriou, E. Parkhomenko, M. T. Pato, T. Paunio, Psychosis Endophenotypes International Consortium, D. O. Perkins, T. H. Pers, O. Pietiläinen, J. Pimm, A. J. Pocklington, J. Powell, A. Price, A. E. Pulver, S. M. Purcell, D. Quested, H. B. Rasmussen, A. Reichenberg, M. A. Reimers, A. L. Richards, J. L. Roffman, P. Roussos, D. M. Ruderfer, V. Salomaa, A. R. Sanders, A. Savitz, U. Schall, T. G. Schulze, S. G. Schwab, E. M. Scolnick, R. J. Scott, L. J. Seidman, J. Shi, J. M. Silverman, J. W. Smoller, E. Söderman, C. C. A. Spencer, E. A. Stahl, E. Strengman, J. Strohmaier, T. S. Stroup, J. Suvisaari, D. M. Svrakic, J. P. Szatkiewicz, S. Thirumalai, P. A. Tooney, J. Veijola, P. M. Visscher, J. Waddington, D. Walsh, B. T. Webb, M. Weiser, D. B.

Wildenauer, N. M. Williams, S. Williams, S. H. Witt, A. R. Wolen, B. K. Wormley, N. R. Wray, J. Q. Wu, C. C. Zai, R. Adolfsson, O. A. Andreassen, D. H. R. Blackwood, E. Bramon, J. D. Buxbaum, S. Cichon, D. A. Collier, A. Corvin, M. J. Daly, A. Darvasi, E. Domenici, T. Esko, P. V. Gejman, M. Gill, H. Gurling, C. M. Hultman, N. Iwata, A. V. Jablensky, E. G. Jönsson, K. S. Kendler, G. Kirov, J. Knight, D. F. Levinson, Q. S. Li, S. A. McCarroll, A. McQuillin, J. L. Moran, B. J. Mowry, M. M. Nöthen, R. A. Ophoff, M. J. Owen, A. Palotie, C. N. Pato, T. L. Petryshen, D. Posthuma, M. Rietschel, B. P. Riley, D. Rujescu, P. Sklar, D. St Clair, J. T. R. Walters, T. Werge, P. F. Sullivan, M. C. O'Donovan, S. W. Scherer, B. M. Neale, J. Sebat, CNV and Schizophrenia Working Groups of the Psychiatric Genomics Consortium, Contribution of copy number variants to schizophrenia from a genome-wide study of 41,321 subjects. *Nat. Genet.* **49**, 27–35 (2017).

83. F. M. G. Cavalli, M. Remke, L. Rampasek, J. Peacock, D. J. H. Shih, B. Luu, L. Garzia, J. Torchia, C. Nor, A. S. Morrissy, S. Agnihotri, Y. Y. Thompson, C. M. Kuzan-Fischer, H. Farooq, K. Isaev, C. Daniels, B.-K. Cho, S.-K. Kim, K.-C. Wang, J. Y. Lee, W. A. Grajkowska, M. Perek-Polnik, A. Vasiljevic, C. Faure-Conter, A. Jouvet, C. Giannini, A. A. Nageswara Rao, K. K. W. Li, H.-K. Ng, C. G. Eberhart, I. F. Pollack, R. L. Hamilton, G. Y. Gillespie, J. M. Olson, S. Leary, W. A. Weiss, B. Lach, L. B. Chambless, R. C. Thompson, M. K. Cooper, R. Vibhakar, P. Hauser, M.-L. C. van Veelen, J. M. Kros, P. J. French, Y. S. Ra, T. Kumabe, E. López-Aguilar, K. Zitterbart, J. Sterba, G. Finocchiaro, M. Massimino, E. G. Van Meir, S. Osuka, T. Shofuda, A. Klekner, M. Zollo, J. R. Leonard, J. B. Rubin, N. Jabado, S. Albrecht, J. Mora, T. E. Van Meter, S. Jung, A. S. Moore, A. R.

Hallahan, J. A. Chan, D. P. C. Tirapelli, C. G. Carlotti, M. Fouladi, J. Pimentel, C. C. Faria, A. G. Saad, L. Massimi, L. M. Liau, H. Wheeler, H. Nakamura, S. K. Elbabaa, M. Perezpeña-Diazconti, F. Chico Ponce de León, S. Robinson, M. Zapotocky, A. Lassaletta, A. Huang, C. E. Hawkins, U. Tabori, E. Bouffet, U. Bartels, P. B. Dirks, J. T. Rutka, G. D. Bader, J. Reimand, A. Goldenberg, V. Ramaswamy, M. D. Taylor, Intertumoral Heterogeneity within Medulloblastoma Subgroups. *Cancer Cell*. **31**, 737–754.e6 (2017).
